# *De novo* chemo-optogenetics through rational small-molecule design and TRAP display

**DOI:** 10.1101/2025.04.10.648175

**Authors:** Tomoki Miyazaki, Tomoshige Fujino, Tatsuyuki Yoshii, Mamoru Funane, Naoya Murata, Chung Nguyen Kim, Satoru Nagatoishi, Kouhei Tsumoto, Gosuke Hayashi, Hiroshi Murakami, Shinya Tsukiji

## Abstract

Optical control of biomolecules is a powerful technique for biological research, but current tools derived from natural proteins face significant limitations. Engineering these proteins to meet researchers’ needs remains challenging. In this study, we present a novel approach to the *de novo* creation of tailored chemo-optogenetic tools. This method combines the rational design of synthetic photo-switchable molecules with *in vitro* selection of artificial protein binders that specifically recognize a particular conformation of these molecules. We demonstrate the utility of this approach by generating artificial proteins that bind exclusively to the *cis*-form of designed azobenzene derivatives. The newly created pairs enable sustained protein activation with a single light pulse and allow for precise, reversible, and repeatable switching of cellular processes using two distinct wavelengths of light in mammalian cells. This *de novo* technique provides tailored synthetic photoswitch–protein binder pairs, expanding the potential for optical biomolecule manipulation.

## Introduction

Mammalian cells possess advanced mechanisms for the spatiotemporal regulation of protein activities across a wide variety of biological and signaling processes.^1–3^ A fundamental and quantitative understanding of living cells requires methods to precisely manipulate specific protein activities in a controlled manner. Demand is growing for experimental tools that facilitate the precise control of protein activity in living cells using light, as the wavelength, intensity, timing, spatial location, and frequency of light can be finely tuned.

Over the past two decades, researchers have harnessed photoreceptor proteins from plants and cyanobacteria to develop optogenetic tools that enable light-dependent regulation of protein activity in living cells.^4–7^ While undoubtedly valuable, these tools have limitations. Flavoprotein-based systems, such as Cry2–CIB1^8^ and iLID-SspB,^9^ induce rapid protein association in response to blue light. However, their complex dissociation occurs spontaneously through thermal deactivation after illumination ends, and thus they require continuous light to maintain the associated (active) state. Phytochrome-based systems, such as PhyB-PIF^10^ and BPhp1-PpsR2,^11^ enable bidirectional control of protein association and dissociation using red and far-red light, respectively, without thermal dissociation. Nevertheless, their broad applicability is constrained by the large size of the phytochrome domain (e.g., 99 kDa in the case of PhyB), making it particularly challenging, or even impossible, to engineer photoreceptor-based tools that simultaneously meet all the needs of researchers.

Synthetic photo-responsive molecules offer an alternative chemical approach for the optical control of biological processes. Recent advancements in orthogonal small molecule/protein binder (or protein tag) pairs, such as benzylguanine/SNAP-tag,^12^ chloroalkane/HaloTag,^13^ trimethoprim/*Escherichia coli* dihydrofolate reductase (eDHFR),^14^ and AP1867/FKBP12(F36V),^15^ have spurred the development of hybrid chemical and genetic strategies. This approach, known as chemo-optogenetics, leverages synthetic photo-functionalized derivatives of small-molecule ligands to regulate the activity of genetically tag-fused proteins of interest with light.^16–18^ Given that synthetic organic chemistry allows for the creation of diverse photo-responsive molecules with tailored photochemical properties, chemo-optogenetics holds great promise as a versatile platform for the optical manipulation of biomolecules. However, the field remains in its infancy, and current chemo-optogenetic tools are largely limited to light-triggered protein dimerization or dissociation systems utilizing photocaged chemical dimerizers consisting of two small-molecule ligands for protein tags.^19–25^ While effective, these methods do not enable repeatable control of protein activity. To overcome this limitation, a photochromic ligand for eDHFR was recently developed, enabling reversible protein dimerization and dissociation with two distinct wavelengths of light.^26,27^ However, the system retains relatively high binding affinity even in the dark state and achieves only a 10-fold increase in affinity upon illumination, presenting practical challenges.

A fundamental limitation of current chemo-optogenetics is its exclusive reliance on the redesign and modification of existing small-molecule ligands originally developed for protein tags (e.g., through caging and azologization) (**Fig. 1a**). This strategy applies only to a limited set of ligands: certain ligands, such as chloroalkane (for HaloTag) and AP1867 [for FKBP12(F36V)], cannot be easily modified. Furthermore, this strategy severely restricts the scope of modifications that can be introduced while preserving the desired protein-binding properties of the ligands.

**Figure 1.**
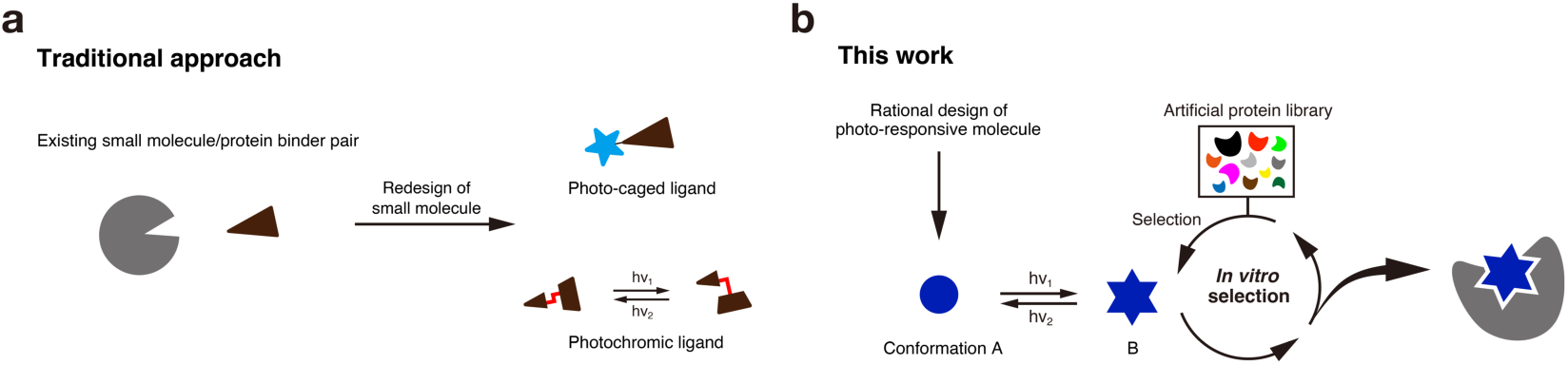
Approaches for the development of chemo-optogenetic tools. (**a**) Traditional approach. (**b**) *De novo* approach proposed in this work.

We here present a novel approach to expanding chemo-optogenetics by integrating the rational design of synthetic photo-responsive molecules with the *in vitro* selection of artificial protein binders. These binders are engineered to specifically recognize a chosen conformation of the molecules using TRAP (<u>tr</u>anscription–translation coupled with <u>a</u>ssociation of <u>p</u>uromycin linker) display (**Fig. 1b**),^28,29^ an advanced mRNA display technique that accelerates selection speed.^30,31^ This strategy facilitates the *de novo* creation of custom-designed pairs of synthetic photo-responsive molecules and artificial protein binders for chemo-optogenetic applications. To demonstrate the proposed technique, we developed artificial protein binders that selectively recognize the *cis*-form of an azobenzene ligand, whose *trans*–*cis* isomerization can be controlled with two different wavelengths of visible light. One binder exhibited high nanomolar affinity for the *cis*-form while showing no detectable binding to the *trans*-form. We further utilized this newly created pair to establish a chemo-optogenetic platform that enables (i) sustained protein-association induction with a single light pulse and (ii) precise, reversible, and repeatable control of protein association and dissociation using dual-wavelength light in mammalian cells. This light-switchable system is a practical tool for manipulating various cellular processes, such as protein kinase and lipid signaling, changes in polarized cell morphology, and gene expression, with precise temporal and spatial control. These results demonstrate the utility and potential of our *de novo* approach, which merges rational small-molecule design with *in vitro* protein selection, paving the way for the creation of customized pairs of synthetic photo-responsive small molecules and artificial protein binders for optical biomolecule manipulation.

## Results

### Basic strategy

An ideal yet unrealized chemo-optogenetic tool would be a light-switchable small molecule–protein binder pair possessing the following properties: (i) minimal to no binding in the dark; (ii) high-affinity association upon light irradiation; (iii) thermal stability of the complex, with no dissociation after irradiation ceases; (iv) efficient dissociation of the complex by light irradiation at a different wavelength; and (v) repeatable bidirectional association and dissociation triggered by dual-wavelength light irradiation. Additionally, it is preferable for the protein binder to be relatively small. In this study, we aimed to develop such a pair *de novo* by combining the rational design of a small molecule with the *in vitro* selection of artificial protein binders using TRAP display. To create a reversibly light-switchable small molecule–protein binder pair, we employed an azobenzene derivative as a small-molecule photoswitch and isolated artificial protein binders that specifically recognize the *cis*-form of the azobenzene ligand.

### Design and characterization of azobenzene ligands

We selected the *ortho*-tetrafluoroazobenzene (*o*-F_4_-azobenzene) diacid structure as our small-molecule scaffold.^32,33^ Azobenzenes have been employed in the development of various small-molecule drugs whose activities can be modulated by light.^34–37^ However, standard (non-fluorinated) azobenzenes often face limitations, such as the need for UV light for *trans*→*cis* photoisomerization, low thermal stability of the *cis*-form (which is prone to rapid isomerization back to the *trans*-form), and incomplete reversibility.^32,33^ In contrast, *o*-F_4_-azobenzene diacid derivatives undergo *trans*→*cis* and *cis*→*trans* isomerizations upon irradiation with visible blue-to-green and violet light, respectively. The *cis*-form is thermally stable and exhibits negligible reverse isomerization in the dark.^33^ Additionally, *cis*→*trans* photoisomerization occurs with nearly quantitative efficiency, while *trans*→*cis* photoisomerization proceeds with moderate efficiency.^32^ Leveraging these properties, we rationally designed Et-FAzo (**1**) and Ca-FAzo (**2**) as small-molecule photoswitches (**Fig. 2a**). In these designs, one end of the *o*-F_4_-azobenzene diacid unit was functionalized with either an ethyl group (Et-FAzo) or a carboxylic acid group (Ca-FAzo) to serve as a potential recognition site for artificial protein binders, while the other end was attached to a flexible linker for further modification. Compounds **1** and **2** were synthesized as detailed in **Schemes S1** and **S2**.

**Figure 2.**
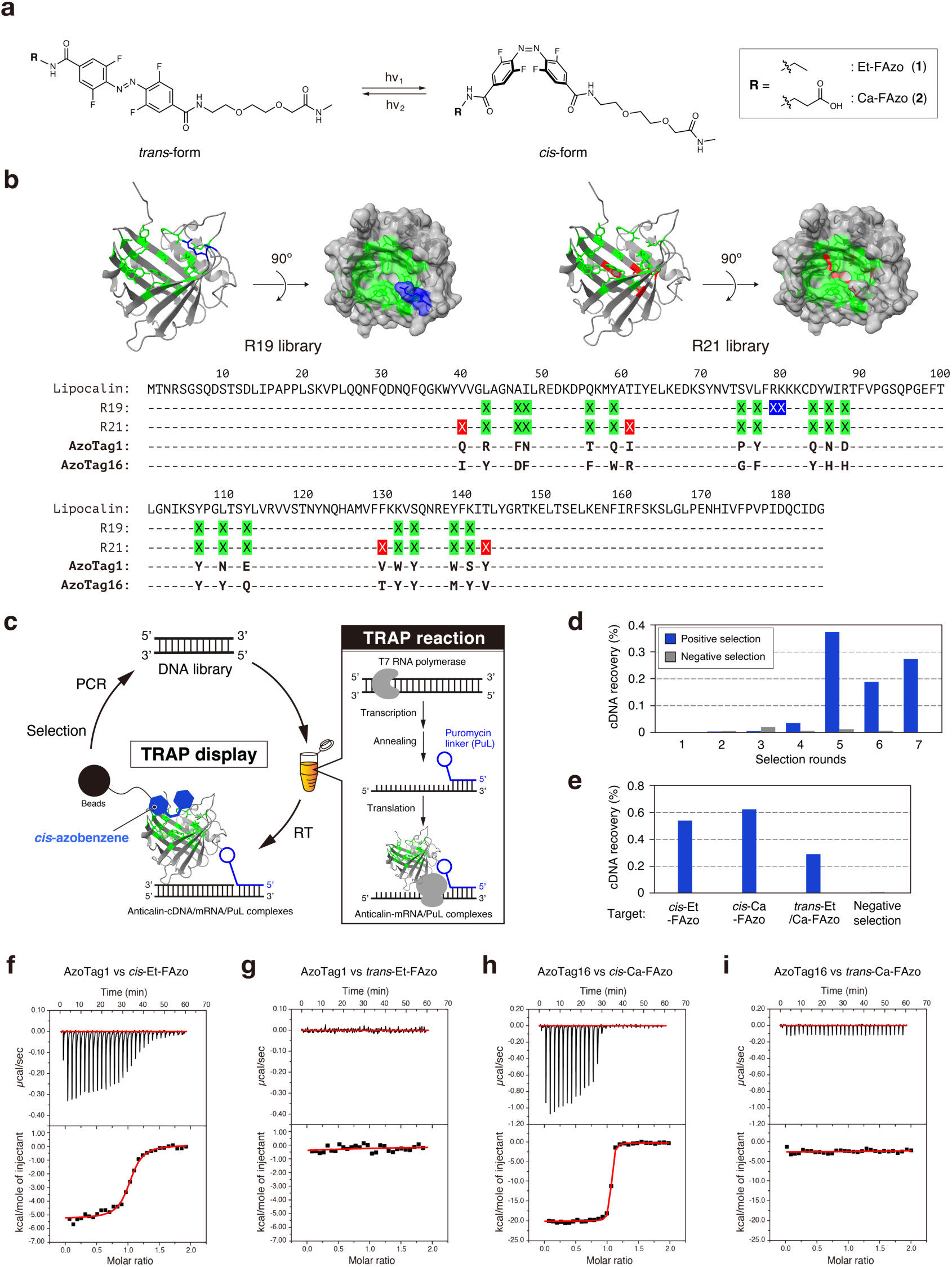
*De novo* creation of *cis*-azobenzene ligand/artificial protein binder pairs. (**a**) Chemical structures of Et-FAzo (**1**) and Ca-FAzo (**2**) ligands. (**b**) Design of anticalin libraries. (Top) Randomized positions in anticalin libraries displayed on the neutrophil gelatinase-associated lipocalin structure (PDB: 1L6M). Green indicates positions randomized in both the R19 and R21 libraries, while blue and red indicate positions unique to the R19 and R21 libraries, respectively. (Bottom) Sequence alignment of the lipocalin, the R19 and R21 anticalin libraries, and AzoTag1 and AzoTag16 obtained through *in vitro* selection. Randomized amino acids are represented as X, following the same color scheme as in the top panel. (**c**) Schematic illustration of the TRAP display. Anticalin/mRNA/PuL complexes were synthesized in the TRAP reaction solution by adding the DNA library. After reverse transcription (RT), anticalin clones that bind to the *cis*-form of FAzo ligands were selectively recovered, and the cDNA of the clones was amplified by PCR for the next selection round. (**d**) Progress of the *in vitro* selection. After each round, the recovered cDNA was quantified using real-time PCR. The blue and gray bars represent the recoveries from positive and negative selections, respectively. (**e**) cDNA recovery rates in the eighth (final) round against different FAzo ligands. (**f**–**i**) Affinity measurements of AzoTag1 for *cis*-Et-FAzo (**f**) and *trans*-Et-FAzo (**g**), and AzoTag16 for *cis*-Ca-FAzo (**h**) and *trans*-Ca-FAzo (**i**). The top panels show titration kinetics, while the bottom ones display the integrated binding isotherms. The binding enthalpy (Δ*H*) and dissociation constant (*K*_d_) were determined by nonlinear regression of the integrated data using the 1:1 binding model. Thermodynamic properties are provided in **Supplementary Table 3.**

To characterize their photochemical and photophysical properties, we isolated the pure *trans-* and *cis*-isomers of **1** and **2** using reversed-phase HPLC (**Supplementary Fig. 1**). The UV/Vis absorption spectra and molar extinction coefficients of the *trans-* and *cis-* isomers were consistent with those reported for diamidated *o*-fluoroazobenzene derivatives (**Supplementary Fig. 2** and **Supplementary Table 1**).^33^ No thermal *cis→trans* isomerization was observed for at least 50 h under dark conditions at either room temperature or 37°C (**Supplementary Fig. 3**). The percentages of the *cis*-form at photostationary states (PSS*_cis_*) upon irradiation with various wavelengths of light are summarized in **Supplementary Fig. 4** and **Supplementary Table 1**. Under green light (516 nm), the *cis-*form predominated (PSS*_cis_*_,516_: 80% for **1**, 81% for **2**), while violet light (400 nm) favored the *trans-*form (PSS*_cis_*_,400_: 5% for **1**, 7% for **2**). Moreover, both compounds demonstrated reversible photoisomerization when exposed to alternating blue and violet light, maintaining stability through multiple cycles (**Supplementary Fig. 5**). These findings indicate that compounds **1** and **2** possess the necessary photochemical and photophysical properties to function as small-molecule photoswitches for generating artificial protein binders.

### Artificial protein library design and *in vitro* selection

Anticalins are small engineered proteins derived from the lipocalin family, consisting of approximately 180 amino acids with a molecular weight of around 21 kDa.^38,39^ Their β-barrel structure contains a cavity that serves as a binding pocket for small molecules.^40–42^ Accordingly, we chose the lipocalin scaffold to isolate artificial protein binders (anticalins) that can recognize the *cis-*forms of the synthetic photoswitches **1** and **2**.

**Figure 2** summarizes our design of the artificial protein library and the *in vitro* selection strategies. The R19 library, which contains randomized amino acid residues at 19 positions facing the cavity within the β-barrel structure, was prepared based on a report by Gebauer and colleagues (**Fig. 2b**).^43^ The newly constructed R21 library included randomized residues at 21 positions, extending to areas near the bottom of the cavity (**Fig. 2b**). Both DNA libraries were created using a codon mix rich in Tyr, Trp, and Phe, as these aromatic amino acids were anticipated to enhance binding to azobenzene molecules through stacking interactions.

We performed *in vitro* selection experiments using TRAP display to isolate anticalin clones that bind to the *cis*-form of Et-FAzo and Ca-FAzo photoswitches (**Fig. 2c** and **Supplementary Fig. 6**). In the first selection round, we conducted *in vitro* transcription reactions for the R19 and R21 DNA libraries separately from the *in vitro* translation in order to maximize the diversity of the anticalin library. The R19 and R21 mRNA libraries, hybridized with a puromycin linker (PuL)-conjugated oligonucleotide, were translated into anticalin-mRNA/PuL complexes in a reconstituted *E. coli in vitro* translation mixture.^44^ After reverse transcription, we achieved over 10^13^ diversity in the anticalin-cDNA/mRNA/PuL library (**Supplementary Fig. 7**). Beads immobilized with Et-FAzo and Ca-FAzo (**Supplementary Fig. 8**) were mixed and irradiated with 500 nm light to photoisomerize the azobenzene ligands into the *cis*-form prior to selection. We incubated the anticalin-cDNA/mRNA/PuL library with the beads and collected the anticalin complexes bound to the *cis*-Et/Ca-FAzo‒immobilized beads. Bound cDNA was eluted by heating the beads to 95°C, followed by PCR amplification. Starting from the second selection round, we included a negative selection step in each round using beads immobilized with a *p*-hydroxybenzoyl-D-tyrosine (HBY) derivative (**Supplementary Fig. 8**) to eliminate undesired anticalins. From the third round onward, DNA was directly added to the TRAP system (**Fig. 2c**) to expedite the selection process. In the fifth round the cDNA recovery rate increased (**Fig. 2d**), so in the sixth and seventh round selections, after capturing the *cis*-Et/Ca-FAzo-binding anticalin complexes on the beads, we eluted the cDNA by irradiating the beads with 410 nm light to photoisomerize the azobenzene ligands from *cis*- to *trans*-form. This approach allowed for the selective elution of anticalin complexes that exhibit *cis*-Et-FAzo or *cis*-Ca-FAzo ligand-specificity and photo-reversibility. In the eighth (final) selection round, we targeted three types of beads: *cis*-Et-FAzo-, *cis*-Ca-FAzo-, and a mixture of *trans*-Et-FAzo- and *trans*-Ca-FAzo‒ immobilized beads. Each collected cDNA was analyzed by next-generation sequencing (**Supplementary Table 2**). The clones most frequently obtained from the selection against *cis*-Et-FAzo and *cis*-Ca-FAzo ligands were identified and named AzoTag1 and AzoTag16, respectively. These two clones demonstrated the highest ratio of cDNA recovery for the *cis-*form compared to the *trans*-form of the azobenzene ligands, indicating their strong selectivity for the *cis*-form.

### Thermodynamic characterization of the Et-FAzo/AzoTag1 and Ca-FAzo/AzoTag16 binding pairs

Isothermal titration calorimetry (ITC) experiments were conducted to assess the binding affinity, binding stoichiometry, and thermodynamic parameters of the Et-FAzo/AzoTag1 and Ca-FAzo/AzoTag16 interactions. Recombinant AzoTag1 and AzoTag16 proteins were expressed in *E. coli* as fusion proteins with NusA (Nus-tag) and a His-tag, and were purified to high purity using affinity chromatography (**Supplementary Fig. 9**). Representative titration curves are presented in **Fig. 2f–i**, with results summarized in **Supplementary Table 3**. For the titration of the *cis*-Et-FAzo ligand with AzoTag1, the dissociation constant (*K*_d_) and binding stoichiometry (*N*) were found to be 804.7 ± 22.6 nM and approximately 1, respectively (**Fig. 2f**). In contrast, no significant binding heat was detected during the titration of the *trans*-Et-FAzo ligand with AzoTag1 (**Fig. 2g**). Similarly, for the titration of the *cis*-Ca-FAzo ligand with AzoTag16, the *K*_d_ and *N* were determined to be 14.03 ± 0.02 nM and approximately 1, respectively (**Fig. 2h**), while no measurable binding heat was observed for the *trans*-Ca-FAzo ligand (**Fig. 2i**).

These findings demonstrate that AzoTag1 and AzoTag16 bind to their corresponding ligands in the *cis-*form with a 1:1 stoichiometry and that both interactions are exothermic. The *cis*-Ca-FAzo/AzoTag16 pair exhibits a higher affinity than the *cis*-Et-FAzo/AzoTag1 pair, as indicated by the Gibbs free energy (Δ*G*) values (AzoTag1: −8.31 ± 0.02 kcal mol^-^ _1_; AzoTag16: −10.71 ± 0.01 kcal mol^-1^). While both pairs exhibit enthalpy-driven binding based on the Δ*H* parameters (AzoTag1: −4.85 ± 0.52 kcal mol^-1^; AzoTag16: −19.31 ± 0.43 kcal mol^-1^), the *cis*-Et-FAzo/AzoTag1 pair shows a greater entropy contribution compared to the *cis*-Ca-FAzo/AzoTag16 pair, as reflected in the −TΔ*S* parameters (AzoTag1: −3.47 ± 0.53 kcal mol^-1^; AzoTag16: 8.59 ± 0.44 kcal mol^-1^).

### Evaluation of Et-FAzo/AzoTag1 and Ca-FAzo/AzoTag16 pairs in mammalian cells

We next aimed to evaluate the binding and photoswitching functions of the *de novo*-created Et-FAzo/AzoTag1 and Ca-FAzo/AzoTag16 pairs in mammalian cells. To achieve this, we designed a protein dimerization system using chimeric FAzo ligands conjugated to the HaloTag ligand (HTL) and HaloTag expressed on the inner leaflet of the plasma membrane (PM) (**Fig. 3a**). This system enables the observation of AzoTag binding to the FAzo ligand through a localization change, where AzoTag translocates from the cytoplasm to the PM. Prior to the assay, Et-FAzo-HTL (**8**) and Ca-FAzo-HTL (**9**) were synthesized by conjugating the respective FAzo ligands to HTL via a linker (**Fig. 3b**). The membrane permeability and intracellular HaloTag-conjugation efficiency of each dimerizer were then assessed using the chloroalkane penetration assay (CAPA).^45^ Et-FAzo-HTL exhibited high membrane permeability and efficient HaloTag conjugation, whereas Ca-FAzo-HTL showed low membrane permeability (**Supplementary Fig. 10**). To address this, the carboxylate group of Ca-FAzo-HTL was modified with an intracellular esterase-cleavable acetoxymethyl group, resulting in Ca(AM)-FAzo-HTL (**10**) (**Fig. 3b**). This modification significantly enhanced both the membrane permeability and HaloTag conjugation efficiency (**Supplementary Fig. 10**), making Ca(AM)-FAzo-HTL suitable for subsequent experiments.

**Figure 3.**
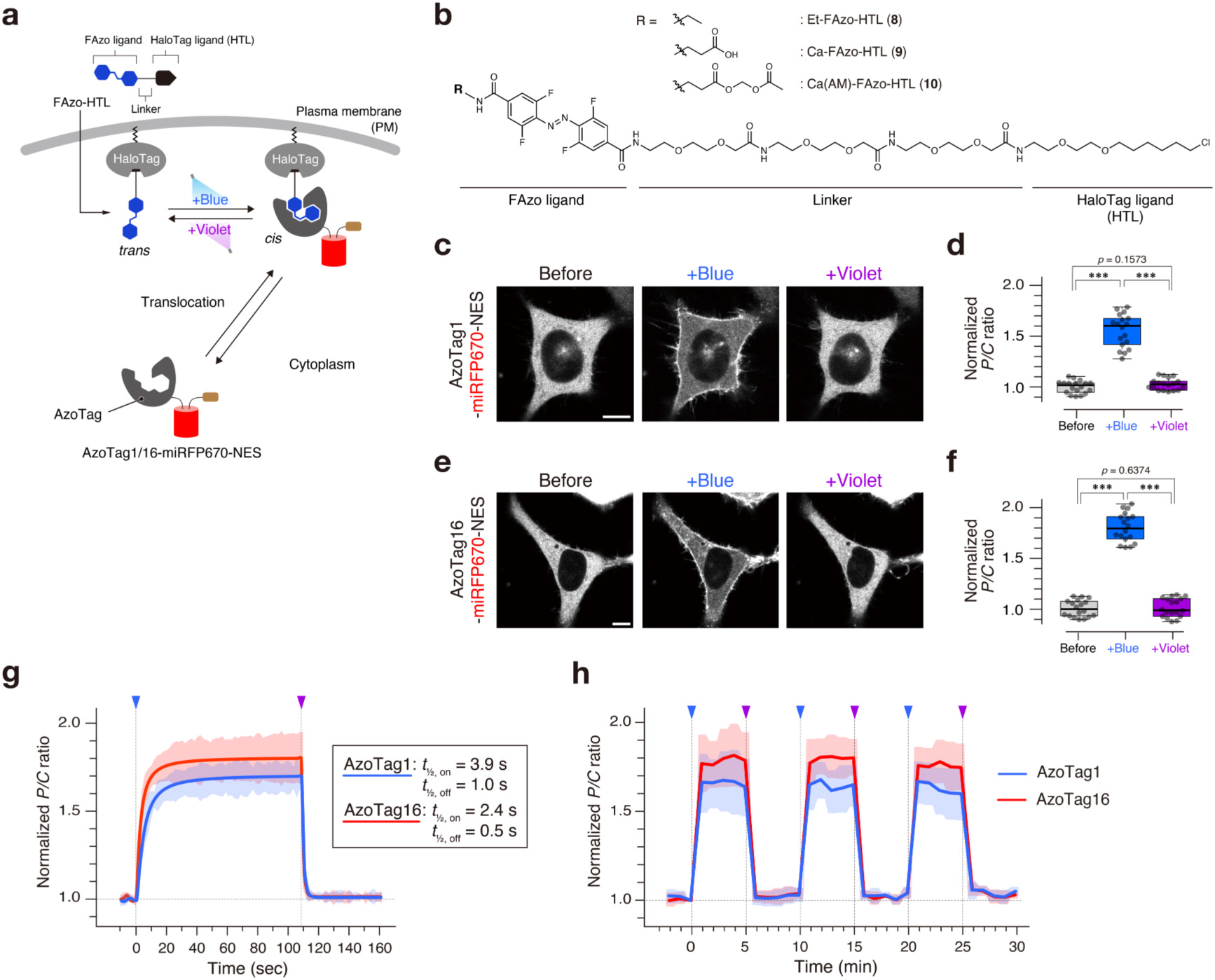
Intracellular applications of Et-FAzo/AzoTag1 and Ca-FAzo/AzoTag16 pairs. (**a**) Schematic illustration of the light-controlled AzoTag-HaloTag dimerization system. (**b**) Chemical structures of AzoTag-HaloTag dimerizers: Et-FAzo-HTL (**8**), Ca-FAzo-HTL (**9**), and Ca(AM)-FAzo-HTL (**10**). (**c**,**e**) Representative confocal fluorescence images of HeLa cells coexpressing AzoTag1-miRFP670-NES with Et-FAzo-HaloTag^PM^ (**c**) or AzoTag16-miRFP670-NES with Ca-FAzo-HaloTag^PM^ (**e**). Images were acquired before and 2 min after irradiation with blue light (483 nm, “+Blue”) and violet light (400 nm, “+Violet”). Scale bars, 10 μm. (**d**,**f**) Quantification of protein translocation efficiencies. The normalized plasma membrane-to-cytoplasm fluorescence intensity ratios (*P/C* ratios) of AzoTag1-(**d**) and AzoTag16-miRFP670-NES (**f**) are presented as box plots (n = 18 cells from three independent experiments per condition). \*\*\**p* < 0.001, Student’s *t*-test. (**g**) Kinetics of light-triggered forward and reverse protein translocation. The confocal fluorescence imaging experiments were performed as described in (**c**,**e**). The normalized *P/C* ratios are presented as the mean ± SD (n = 5 cells from three independent experiments per construct) and plotted as a function of time. Arrowheads indicate irradiation with 483 nm (blue) and 400 nm (violet) light. The half-maximal translocation times for association (*t*_1/2,on_) and dissociation (*t*_1/2_,_off_) were determined from the fitting curves using a single exponential function. For time-lapse movies, see **Supplementary Video 1**. (**h**) Repetitive photo-switching of protein translocation. The confocal fluorescence imaging experiments were performed as described in (**c**,**e**), and data are depicted as described in (**g**) (n = 10 cells from three independent experiments per construct). For time-lapse movies, see **Supplementary Video 2**.

For the dimerization assay (**Fig. 3a**), AzoTag1 and AzoTag16 were fused to miRFP670 and a nuclear export signal (NES) sequence,^46^ creating AzoTag1-miRFP670-NES and AzoTag16-miRFP670-NES constructs for cytosolic expression. HaloTag was fused to the C-terminal sequence of KRas4B (HaloTag^PM^) to localize it to the inner leaflet of the PM. The AzoTag1 and AzoTag16 fusion constructs were coexpressed with HaloTag^PM^ in HeLa cells at equimolar ratios using a self-cleaving P2A peptide sequence.^47^ The successful expression of both AzoTag fusion proteins was confirmed through confocal fluorescence imaging (**Fig. 3c,e**). In the absence of their respective FAzo ligands, the AzoTag fusion proteins were diffusely distributed throughout the cytoplasm.

To assess the light-dependent binding between Et-FAzo and AzoTag1 in cells, we prepared Et-FAzo-conjugated HaloTag^PM^ by incubating cells with *trans*-Et-FAzo-HTL (10 µM) for 30 min, followed by washing to remove excess unbound ligand. Notably, the localization of AzoTag1-miRFP670-NES remained unchanged in the absence of light irradiation (**Fig. 3c,e**, and **Supplementary Fig. 11**). However, when irradiated with blue light (483 nm), which induces *trans*→*cis* photoisomerization, AzoTag1-miRFP670-NES translocated from the cytoplasm to the PM, significantly increasing the normalized fluorescence intensity ratio (*P/C*) between the PM and cytoplasm (1.58) (**Fig. 3d**). Subsequent violet light irradiation (400 nm) caused complete relocalization of AzoTag1-miRFP670-NES from the PM back to the cytoplasm, reducing the *P/C* value almost to its pre-irradiation level (**Fig. 3d**). No light-induced translocation of AzoTag1-miRFP670-NES was observed in the cells not treated with Et-FAzo-HTL (**Supplementary Fig. 12**). These results demonstrate that AzoTag1 binds specifically to the *cis*-form of the Et-FAzo ligand in mammalian cells, and that the association and dissociation of AzoTag1 and Et-FAzo can be reversibly controlled by photoswitching the *trans*/*cis* state of the Et-FAzo ligand.

We performed the same assay for the Ca-FAzo/AzoTag16 pair using HaloTag^PM^ and Ca(AM)-FAzo-HTL. AzoTag16-miRFP670-NES showed no binding to the *trans*-form of Ca-FAzo-conjugated HaloTag^PM^ but specifically bound to its *cis*-form after blue light irradiation, yielding a normalized *P/C* value of 1.83 (**Fig. 3e,f** and **Supplementary Fig. 11**). Nearly complete reverse translocation of AzoTag16-miRFP670-NES from the PM to the cytoplasm was also observed upon subsequent violet light irradiation (**Fig. 3f**), demonstrating high reversible binding through light switching.

We further evaluated the intracellular association and dissociation kinetics of the Et-FAzo/AzoTag1 and Ca-FAzo/AzoTag16 pairs. The half-times for blue-light-induced association (*t*_1/2,on_) and violet-light-induced dissociation (*t*_1/2,off_) were calculated to be 3.9 ± 0.2 s and 1.0 ± 0.2 s for the Et-FAzo/AzoTag1 pair, and 2.4 ± 0.1 s and 0.5 ± 0.1 s for the Ca-FAzo/AzoTag16 pair, respectively (**Fig. 3g** and **Supplementary Video 1**). These results indicate that both pairs undergo rapid, light-triggered association and dissociation on the scale of seconds. Additionally, the light-switching of association and dissociation could be repeatedly and precisely controlled (**Fig. 3h** and **Supplementary Video 2**).

Taken together, the results showed that *de novo*-created Et-FAzo/AzoTag1 and Ca-FAzo/AzoTag16 pairs function as light-switchable modules that interact in a *cis*-form‒ specific manner in mammalian cells. Notably, both pairs exhibit all-or-none binding behavior: no binding occurs prior to light irradiation (when the FAzo ligand is in the *trans*-form), high-affinity binding is induced by blue light irradiation (*trans*→*cis* photoisomerization), and near-complete dissociation occurs upon violet light irradiation (*cis*→*trans* reverse photoisomerization). Although both pairs demonstrate excellent light-switching properties, we selected the Ca-FAzo/AzoTag16 pair for subsequent experiments because of its slightly higher *cis*-form binding affinity, superior P/C value, and faster association‒dissociation kinetics upon photoswitching.

### Reversible dual light control of protein kinase signaling

We next investigated whether the dual light-switchable Ca-FAzo/AzoTag16 pair could serve as a novel chemo-optogenetic platform for the precise and reversible control of intracellular processes. To this end, we utilized the AzoTag16/Ca-FAzo-HaloTag dimerization system constructed above, as light-switchable protein dimerization provides a versatile tool for various applications in chemical biology and synthetic biology.

We first focused on cRaf, a protein kinase that regulates various cellular functions, including growth, survival, and differentiation.^48^ When cRaf is recruited to the PM, it undergoes activation via autophosphorylation, which triggers the activation of downstream MEK and ERK.^48^ To control the PM localization and activity of cRaf, we fused full-length cRaf to the C-terminus of AzoTag16-miRFP680 (AzoTag16-miRFP680-cRaf) (**Fig. 4a**).^49^ This protein was coexpressed with HaloTag^PM^ in HeLa cells, followed by conjugation of HaloTag^PM^ with Ca(AM)-FAzo-HTL. The activity of the cRaf/ERK pathway was monitored using an ERK reporter fused to mScarlet-I (mScarlet-ERK),^50^ which translocates from the cytoplasm to the nucleus upon ERK activation.

**Figure 4.**
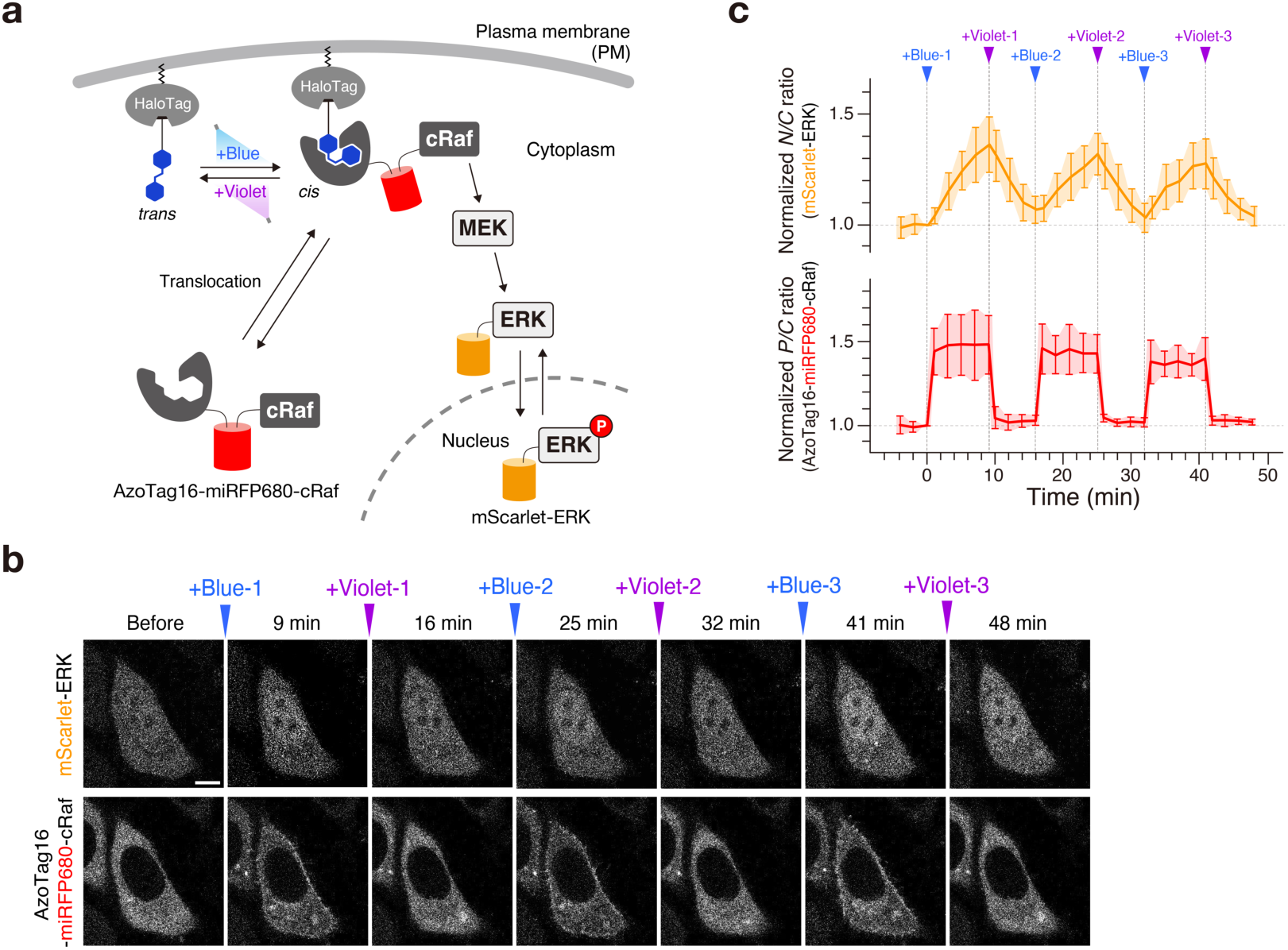
Reversible dual-light control of cRaf/ERK signaling. (**a**) Schematic illustration of the experimental setup using AzoTag16-fused cRaf. (**b**) Representative time-lapse confocal fluorescence images of HeLa cells coexpressing AzoTag16-miRFP680-cRaf, Ca-FAzo-HaloTag^PM^, and mScarlet-ERK before and after sequential irradiation with blue light (483 nm, “+Blue”) and violet light (400 nm, “+Violet”). Images were acquired at the indicated time points. Scale bar, 10 μm. For time-lapse movies, see **Supplementary Video 4**. (**c**) Time course of cRaf translocation and ERK activation. The normalized *P/C* ratios of AzoTag16-miRFP680-cRaf are plotted as a function of time to evaluate cRaf translocation (bottom). The normalized nucleus-to-cytoplasm fluorescence intensity ratios (*N/C* ratios) of mScarlet-ERK are plotted as a function of time to assess ERK activity (top). Arrowheads indicate irradiation with 483 nm (blue) and 400 nm (violet) light. Data are presented as the mean ± SD (n = 10 cells from three independent experiments).

Prior to light irradiation, AzoTag16-miRFP680-cRaf exhibited a cytoplasmic distribution, while mScarlet-ERK was also predominantly localized in the cytoplasm (**Fig. 4b**), indicating low ERK activity. Upon brief irradiation with blue light for 30 s, AzoTag16-miRFP680-cRaf translocated to the PM (**Fig. 4b**). Following this translocation, mScarlet-ERK moved to the nucleus, indicating ERK activation (**Fig. 4b**). The nuclear translocation of mScarlet-ERK reached a plateau approximately 20 min after light irradiation, and the PM localization of AzoTag16-miRFP680-cRaf, along with the ERK activity, remained stable for at least 2 h without additional light exposure (**Supplementary Fig. 13** and **Supplementary Video 3**). Such ERK activation was not observed when the same experiment was performed using AzoTag16-miRFP670-NES, which lacks the cRaf protein (**Supplementary Fig. 14**), or in the absence of light irradiation (**Supplementary Fig. 15**). These results demonstrate that the AzoTag16/Ca-FAzo-HaloTag^PM^ system enables unidirectional, sustained PM recruitment and activation of AzoTag16-fused cRaf via blue light-triggered *trans*→*cis* isomerization.

Furthermore, by alternately irradiating cells with blue and violet light, inducing *trans*→*cis* and *cis*→*trans* isomerization of the Ca-FAzo ligand, respectively, we achieved reversible and repeated control of the PM localization of AzoTag16-miRFP680-cRaf (**Fig. 4b,c** and **Supplementary Video 4**). This dual light-based manipulation synchronously switched ERK activity on and off, demonstrating its ability to control synthetic ERK signal oscillations.

### Dual light control of spatially localized lipid signaling

The light-switchable AzoTag16/Ca-FAzo-HaloTag^PM^ system also proved useful for controlling lipid signaling. Phosphatidylinositol 3-kinase (PI3K) is a lipid kinase that phosphorylates phosphatidylinositol 4,5-bisphosphate [PI(4,5)P_2_] to produce phosphatidylinositol 3,4,5-trisphosphate [PI(3,4,5)P_3_] upon PM recruitment.^51^ We generated a synthetic light-switchable PI3K by fusing AzoTag16-miRFP680 with the inter-Src homology 2 (iSH2) domain derived from the p85 subunit of PI3K (AzoTag16-miRFP680-p85_iSH2_) (**Supplementary Fig. 16a**).^49^ The iSH2 domain spontaneously forms a complex with the endogenous p110 subunit of PI3K in cells.^52^ AzoTag16-miRFP680-p85_iSH2_ was coexpressed in HeLa cells with HaloTag^PM^ and a PI(3,4,5)P_3_ biosensor consisting of the PH domain of Akt fused to mScarlet-I (mScarlet-PH_Akt_).^53^ Following conjugation with Ca(AM)-FAzo-HTL, we were able to reversibly and repeatedly control the acute production and depletion of PI(3,4,5)P_3_ by alternating blue and violet light irradiation across the entire dish (**Supplementary Figs. 16–18** and **Supplementary Video 5**).

Locally generated PI(3,4,5)P_3_ at the PM is crucial for regulating cell membrane dynamics and polarized cell morphology.^54^ We thus attempted to artificially control the local production of PI(3,4,5)P_3_ in Cos-7 cells using the light-switchable PI3K (**Fig. 5a**). Confined light scanning of a local area of the cell with a 488 nm laser rapidly induced directed membrane protrusion—specifically, lamellipodium formation—around the irradiated PM region. This protrusion was accompanied by membrane retraction on the opposite side of the cell (**Fig. 5b,c**). The light-induced local membrane protrusion typically continued for 30 min (**Supplementary Fig. 19** and **Supplementary Video 6**). However, when the entire cell was irradiated with a 405 nm laser during the membrane protrusion process, the local membrane protrusion was halted (**Fig. 5b,c**). We were also able to repeatedly switch the directed membrane protrusion on and off in a user-defined manner by combining local 488 nm laser irradiation with whole-cell 405 nm laser irradiation (**Fig. 5b,c** and **Supplementary Video 7**). The directional lamellipodium formation was not observed when the entire cell was uniformly irradiated (**Supplementary Fig. 20**). Additionally, no lamellipodium formation occurred with AzoTag16-miRFP670-NES, which lacks the iSH2 domain (**Supplementary Fig. 21**). These results demonstrate that the AzoTag16/Ca-FAzo-HaloTag^PM^-based PI3K activation tool enables rapid, reversible, and repeatable production of PI(3,4,5)P_3_, allowing for precise control of the polarized cell morphology with high spatiotemporal resolution.

**Figure 5.**
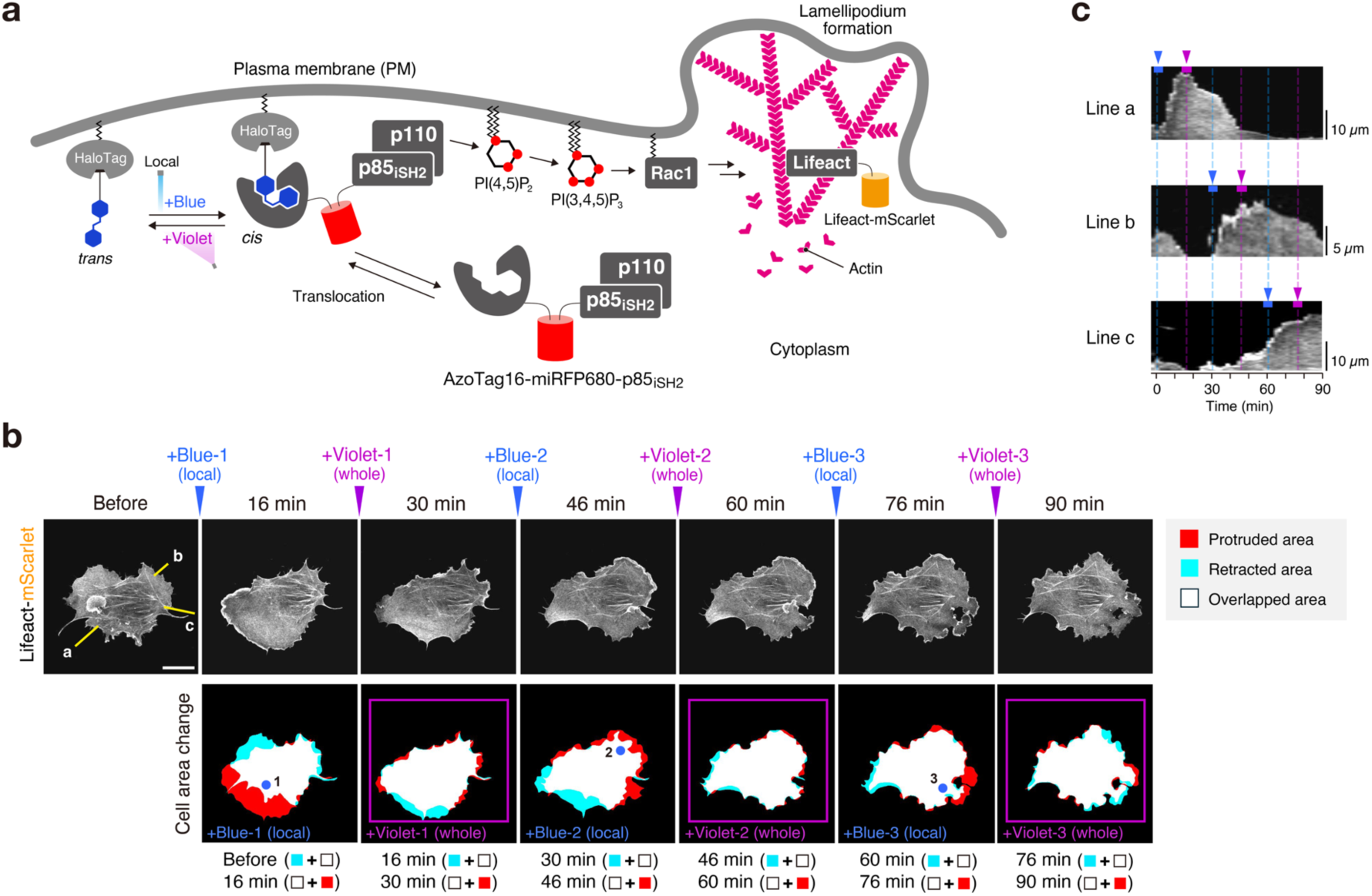
Reversible spatially localized PI3K activation and lamellipodia formation. (**a**) Schematic illustration of the experimental setup using AzoTag16-fused PI3K. (**b**) Representative time-lapse confocal fluorescence images of a Cos-7 cell coexpressing AzoTag16-miRFP680-p85_iSH2_, Ca-FAzo-HaloTag^PM^, and Lifeact-mScarlet before and after sequential local and whole-cell irradiation with 488 nm laser (“+Blue”) and 405 nm laser (“+Violet”) light, respectively. The +Blue images were acquired 16 min after 488 nm laser irradiation, while the +Violet images were acquired 14 min after 405 nm laser irradiation. The blue circles 1–3 indicate the regions of local 488 nm laser irradiation, while the violet squares indicate the regions of 405 nm laser irradiation. Scale bar, 20 μm. For time-lapse movies, see **Supplementary Video 7**. (**c**) Kymographs of cell edges. The kymographs were generated along the yellow lines a–c in (**b**) and aligned as a function of time. Arrowheads indicate irradiation with 488 nm laser light (blue) at the indicated site and with 405 nm laser light (violet) at the whole cell.

### Light-controlled gene expression via a CRISPR-dCas activation system

We next examined whether sustained interaction of the Ca-FAzo/AzoTag16 pair upon light irradiation could be used to control gene expression. Here, we utilized the CRISPR-dCas activation system—which consists of a transcription factor fused to nuclease-dead Cas9 (dCas9)—to regulate gene expression at specific target sites via guide RNA (gRNA).^55–57^ We designed and constructed a light-responsive CRISPR-dCas system by fusing dCas9 with AzoTag16 (dCas9-AzoTag16) and a transcriptional activator VPR with HaloTag (HaloTag-VPR) (**Fig. 6a**). The two constructs and the gRNA expression plasmid were transiently coexpressed with a reporter gene encoding mAzamiGreen (mAG)^56,57^ in HeLa cells. After 24 h, the cells were treated with Ca(AM)-FAzo-HTL and exposed to blue light for 2 min (**Fig. 6b**). Following an additional 24-h incubation in the dark to allow for gene expression, we observed a significant (∼10-fold) increase in the fraction of mAG-positive cells among the transfected population (**Fig. 6c,d**). In contrast, no gene expression induction was observed in cells that were either not treated with Ca(AM)-FAzo-HTL (**10**) or not exposed to light. These results demonstrate that the AzoTag16/Ca-FAzo pair can effectively control gene expression in mammalian cells when combined with the CRISPR-dCas activation system. Notably, this system induced gene expression with a single, brief light exposure, unlike most existing light-induced gene expression systems, which typically require continuous light irradiation.

**Figure 6.**
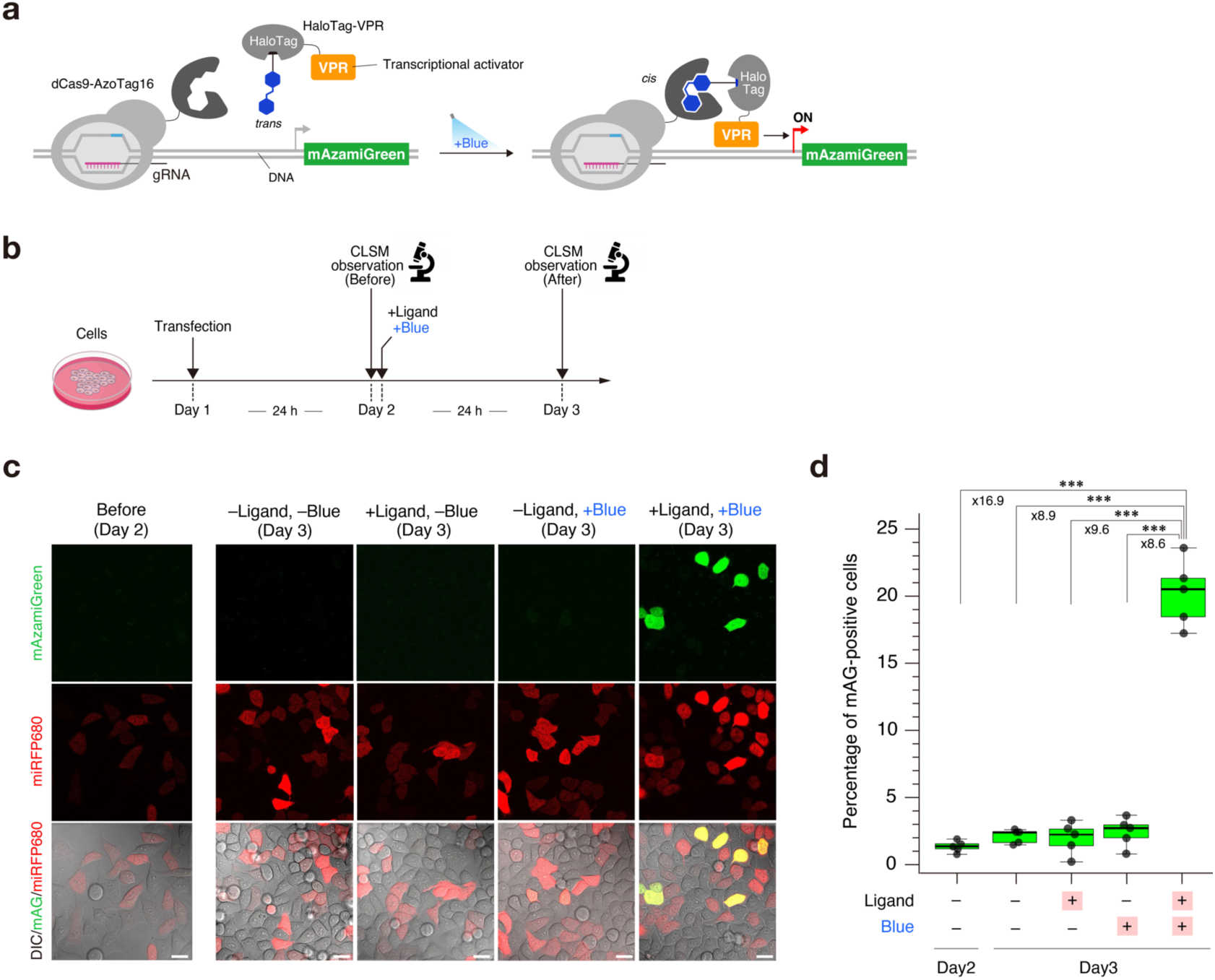
Optical regulation of gene expression by the CRISPR-dCas system. (**a**) Schematic illustration of the experimental setup using the AzoTag16-based transcription activation system. (**b**) Timeline of the light-controlled gene expression experiment. HeLa cells were cotransfected with dCas9-AzoTag16, HaloTag-VPR, miRFP680 (transfection marker), and an mAG-encoding reporter gene on Day 1, observed by confocal laser scanning microscopy (CLSM) on Day 2, treated with Ca(AM)-FAzo-HTL and blue light (483 nm), and then observed by CLSM on Day 3. (**c**) Representative confocal fluorescence images showing light-dependent mAG expression. Control experiments were performed as indicated. Images were acquired under identical conditions. Scale bars, 20 μm. (**d**) Quantification of light-induced gene expression. mAG-positive cells among transfected (miRFP680-expressing) cells under the indicated conditions are presented as box plots (n > 2,500 cells from five independent experiments per condition). \*\*\**p* < 0.001, Student’s *t*-test.

## Discussion

We presented a strategy to create tailor-designed chemo-optogenetic tools *de novo* through rational small-molecule design and *in vitro* protein selection. Using this approach, we generated artificial anticalin-based proteins that specifically bind to the *cis*-form of rationally designed *o*-F_4_-azobenzene derivatives. The Et-FAzo/AzoTag1 and Ca-FAzo/AzoTag16 pairs developed in this study exhibit dual light-dependent binding behaviour with an all-or-none response that is characterized by: (i) no binding prior to light irradiation (with ligands in the *trans*-form); (ii) high-affinity binding in the nanomolar range upon blue light-mediated *trans*→*cis* photoisomerization; (iii) high thermal stability of the complexes after cessation of light irradiation; (iv) near-complete dissociation upon violet light-induced *cis*→*trans* reverse photoisomerization; and (v) repeatable association‒dissociation switching controlled by dual light stimuli. Although light-switchable small molecule–protein pairs were previously developed using caging^21,22^ or azologization^26,27,58,59^ strategies applied to existing small-molecule ligands, none exhibit the full set of properties described above. In this regard, azologization has been recognized as an effective strategy for generating reversibly light-switchable (photochromic) ligands.^18,34–37^ However, the photochemical properties and light-dependent protein-binding characteristics of the resulting molecules cannot be predicted prior to synthesis, making it a challenge to develop light-switchable small molecules with desired properties, even with extensive optimization. In contrast, we here demonstrated that light-switchable small molecule/artificial protein pairs with ideal properties can be created directly by using well-characterized azobenzene molecules as ligands for TRAP display-based protein selection. This underscores the advantage of our *de novo* strategy over conventional approaches that modify established small molecule/natural protein pairs, providing a new conceptual framework for developing light-responsive molecular systems.

We also demonstrated that the *de novo-*created FAzo/AzoTag pairs function in mammalian cells, enabling both sustained protein association with a single light pulse and dual-light mediated, rapid (within seconds), reversible, and repeatable control of protein association when combined with HaloTag. The Ca-FAzo/AzoTag16 pair was further utilized to achieve spatiotemporal control over various biological processes, including protein kinase signaling, lipid modification, polarized cell morphology changes, and gene expression. Additionally, we showcased the broad applicability of this pair for controlling protein localization at various cellular sites (**Supplementary Fig. 22**). These findings highlight the potential of FAzo/AzoTag pairs as a versatile chemo-optogenetic platform for reversible, all-or-none switching of molecular activities at specific locations with high spatiotemporal precision. Notably, these pairs offer distinct advantages as complementary tools to the optogenetic PhyB/PIF system. AzoTag proteins (21 kDa) are considerably smaller than the PhyB domain (99 kDa), and FAzo ligands respond to violet and blue light, whereas PhyB is activated by red light and inactivated by far-red light. In addition to their applications in cell biology, FAzo/AzoTag pairs show promise as light-switchable molecular components for the bottom-up construction of artificial functional molecular systems, such as synthetic cells.^60–62^

In addition to establishing the utility of FAzo/AzoTag pairs, this work significantly expands the scope of *in vitro* selection for engineering functional artificial protein binders that target small molecules. The mRNA display technique and its variant, TRAP display, are powerful tools for efficiently discovering and isolating artificial protein binders from a protein library based on their binding specificity.^29,63–66^ We envision that extending our TRAP display-based protein selection platform to include other photo-switchable small molecules, such as red-shifted azobenzene,^67^ arylazopyrazole,^68^ diarylethene,^69^ and spiropyran derivatives,^70^ will enable the *de novo* creation of a diverse range of light-dependent small molecule/artificial protein binder pairs. These pairs could serve as chemo-optogenetic modules for manipulating intracellular molecular processes with high spatiotemporal resolution using various light stimuli. This advancement will further broaden the potential of *de novo* chemo-optogenetics for biological and biomedical applications.

## Acknowledgements

We thank Hirohide Saito (The University of Tokyo) for providing the mAG-encoding reporter and gRNA expression plasmids. We are grateful to Kai Tahara, Masaru Yoshikawa, and Natsumi Fukaya (Nagoya Institute of Technology) for their technical assistance. This work was supported by JSPS Grants-in-Aid for Scientific Research (KAKENHI) [JP21H05226 (S.T.), JP21K05270 (T.F.), and JP23H05456 (H.M.)], a JST PRESTO grant [JPMJPR178B (T.Y.)], and an AMED grant [JP21zf0127004 (H.M.)], and by funds from the Toyota Riken Scholar Project (T.Y.) and the OptoBioTechnology Research Center at Nagoya Institute of Technology (S.T.). Partial support was also provided by the Platform Project for Supporting Drug Discovery and Life Science Research [Basis for Supporting Innovative Drug Discovery and Life Science Research (BINDS)] from AMED [JP21am0101094 (K.T.)].

## Author contributions

T.F., T.Y., H.M., and S.T. initiated and designed the project. T.M. and T.Y. synthesized the compounds and conducted all experiments except for the *in vitro* selection experiments. T.F., M.F., and N.M. performed the *in vitro* selection experiments. C.N.K. assisted with the protein expression and affinity measurement experiments. S.N. and K.T. provided assistance with the ITC experiments. H.M. and S.T. supervised the project. T.M., T.F., T.Y., G.H., H.M., and S.T. analyzed the data. T.M., T.F., S.N., S.T., and H.M. wrote the manuscript. All authors discussed and commented on the manuscript.

## Methods

### Synthesis

Detailed procedures for synthesizing and characterizing the compounds can be found in the **Supplementary Methods**.

### Photochemical properties of FAzo ligands

Et-FAzo (**1**) and Ca-FAzo (**2**) ligands were purified into their *trans*- and *cis*-forms as described in the **Supplementary Methods**. The purified *trans*- and *cis*-Et-FAzo ligands were each dissolved in DMSO to prepare 10 mM stock solutions and stored at –20°C. The purified *trans*- and *cis*-Ca-FAzo ligands were also dissolved in DMSO to prepare 6.25 mM stock solutions, which were protected from light and stored at –20°C.

In all experiments, *trans*-Et-FAzo, *cis*-Et-FAzo, *trans*-Ca-FAzo, and *cis*-Ca-FAzo (50 µM) were dissolved in HEPES buffer (20 mM HEPES, 150 mM NaCl, pH 7.5) with either 0.5% DMSO for Et-FAzo or 0.8% DMSO for Ca-FAzo. Ultraviolet-visible absorption spectra were measured at 25°C using a Hitachi U-3900H spectrophotometer and a quartz cell with a 1 cm optical path length. Light irradiation was performed using a 300 W xenon light source (MAX-303; Asahi Spectra) equipped with bandpass filters of various wavelengths (360 ± 6 nm, Semrock FF01-360/12-25; 380 ± 7 nm, Semrock FF02-380/14-25; 400 ± 20 nm, Semrock FF01-400/40-25; 420 ± 5 nm, Semrock FF01-420/10-25; 438 ± 12 nm, Semrock FF02-438/24-25; 483 ± 16 nm, Semrock FF01-483/32-25; and 514 ± 15 nm, Semrock FF01-514/30-25).

To determine the PSS*_cis_* values, 1 mL solutions of *trans*-Et-FAzo and *trans*-Ca-FAzo (50 µM) in HEPES buffer were irradiated with the designated wavelengths for 5 min at 25°C. The *trans*- and *cis*-forms of the FAzo ligands were separated and quantified by reversed-phase HPLC using a Hitachi LaChrom Elite system with a YMC-Pack C4 column (10 × 250 mm) for Et-FAzo and a YMC-Pack ODS-A column (10 × 250 mm) for Ca-FAzo. Elution was performed with a linear gradient from 40% to 45% MeCN/H_2_O containing 0.1% trifluoroacetic acid (TFA) over 30 min for Et-FAzo and from 40% to 42% MeCN/H_2_O containing 0.1% TFA over 30 min for Ca-FAzo. Detection occurred at 440 nm for Et-FAzo and 442 nm for Ca-FAzo, corresponding to the isosbestic points of the *trans*- and *cis*-FAzo ligands. The PSS*_cis_* values were calculated by integrating the peak area ratios of the *trans*- and *cis*-forms.

To evaluate the thermal stability of the FAzo ligands in the *cis*-form, 5 mL solutions of *cis*-Et-FAzo and *cis*-Ca-FAzo (50 µM) in HEPES buffer were maintained in the dark at 25°C or 37°C, with their absorption spectra measured at specified time points over 48 h.

In the repetitive photo-switching experiments, 1 mL solutions of *trans*-Et-FAzo and *trans*-Ca-FAzo (50 µM) in HEPES buffer were alternately irradiated with 483 nm and 400 nm light, and their absorption spectra were recorded before and after each irradiation step.

### Preparation of ligand-immobilized magnetic beads

The ligand derivatives and the methods for preparing ligand-immobilized beads used for *in vitro* selection are shown in **Supplementary Fig. 8**. Detailed procedures for the preparation of ligand-immobilized beads can be found in the **Supplementary Methods**.

### Preparation of anticalin mRNA libraries

Two anticalin libraries, R19 and R21, were prepared, each featuring different positions of randomized residues. Those positions in the R19 library matched those reported by Gebauer et al.^43^ The R21 library was newly designed based on R19. For randomized residues, we used a codon mix containing 20% Tyr, 10% Trp, 10% Phe, and 3.75% each of all other amino acids except Cys. We focused on the ratio of aromatic amino acids because we expected them to establish stacking interactions with azobenzenes. The ratio of Tyr was particularly increased because it is known to be rich in the complementarity-determining region (CDR)-H3 loop of immunoglobulin G (IgG).^71^

DNAs coding the anticalin libraries were prepared by ligating four DNA fragments— A, B, C, and D—from the 5’ end of the anticalin gene. Fragments B and C contained randomized residues in their sequences. Oligonucleotides used for the preparation of anticalin libraries are listed in **Supplementary Table 4**.

To prepare fragment B of the R19 library (fragment B-R19), LcncoR19F1.F58 (2 µM), LcncoR19F2.F49-P (2 µM), and LcncoR19F3.F81-P (2 µM) were ligated using T4 DNA ligase (75 µL) with the aid of Lcnan0.R21-NH_2_ (5 µM) and Lcnan1.R20-NH_2_ (5 µM). For fragment C of R19 (fragment C-R19), LcncoR19F4.F75-P (2 µM) and LcncoR19F5.F90-P (2 µM) were ligated (75 µL) with the aid of Lcnan3.R20-NH_2_ (5 µM). For fragment B of R21 (fragment B-R21), LcncoR21F1.F58 (2 µM), LcncoR21F2.F49-P (2 µM), and LcncoR21F3.F81-P were ligated (75 µL) with the aid of Lcnan0.R21-NH_2_ (5 µM) and Lcnan1.R20-NH_2_ (5 µM). For fragment C of R21 (fragment C-R21), LcncoR19F4.F75-P (2 µM) and LcncoR21F5.F90-P (2 µM) were ligated (75 µL) with the aid of Lcnan3.R20-NH_2_ (5 µM). Following the ligation, 75 µL of each reaction mixture was added to a PCR reaction pre-mix [15 mL; 10 mM Tris-HCl (pH 8.4), 100 mM KCl, 0.1% (v/v) Triton X-100, 2% (v/v) dimethyl sulfoxide, 2 mM MgSO_4_, and 0.2 mM of each deoxynucleotide triphosphate (dNTP)] containing 6 nM Pfu-S DNA polymerase^29^ along with the following primer pairs: 0.375 µM LcnBsaI.F33 and 0.375 µM LcnBsaI2.R33 for fragments B-R19 and B-R21, and 0.375 µM LcnBsaI2.F37 and 0.375 µM LcnBsaI.R32 for fragments C-R19 and C-R21. DNAs were amplified by seven PCR cycles.

Fragments A and D did not contain randomized residues and were used for both the R19 and R21 libraries. Both fragments were prepared using two rounds of PCR from the lipocalin gene (**Supplementary Table 4**). For fragment A, LcnPri1.F62 and LcnBsaIFront.R33 were used for the first PCR, and T7exG.F23 and 0.5 µM LcnBsaIFront.R33 were used for the second. As primers for fragment D, LcnBsaIEnd.F33 and LcnG5S-4.R40 were used for the first PCR and LcnBsaIEnd.F33 and LcnG5S-4.R40 for the second.

The amplified fragments A, B, C, and D were purified by phenol/chloroform extraction and isopropanol precipitation. The products were digested with Bsa I (New England Biolabs) following the manufacturer’s protocol, and the DNA products were again purified by phenol/chloroform extraction and isopropanol precipitation. The resultant fragments were ligated to synthesize full-length DNA products (1 µM of each fragment, totaling 200 µL for each library). Fragments A, B-R19, C-R19, and D were used for library R19, while fragments A, B-R21, C-R21, and D were used for library R21. The ligated DNAs were PCR-amplified using primers T7exG.F23 and G5S-4Gan21-3.R42 (60 mL for each, 5 cycles). The products were purified by phenol/chloroform extraction and isopropanol precipitation and then dissolved in 600 µL of 10 mM Tris-acetate (pH 7.6).

The R19 and R21 DNA library templates were transcribed using *in vitro* runoff transcription [22 mL for each; 40 mM Tris-HCl (pH 8.0), 1 mM spermidine, 0.01% Triton X-100, 10 mM dithiothreitol (DTT), 25 mM MgCl_2_, 5 mM each NTP, 1 µM T7 RNA polymerase, and 100 µL of the PCR product] at 37°C for 6 h. The resulting mRNA was purified by isopropanol precipitation followed by polyacrylamide gel electrophoresis (PAGE) purification.

The complex of mRNA and hexachlorofluorescein-labelled puromycin-linker (HEX-PuL) was prepared (62.5 µL) by annealing mRNA (4 µM) and HEX-PuL (4 µM) in an annealing buffer [25 mM HEPES-KOH (pH 7.8), 200 mM potassium acetate] by heating the solution to 95°C for 2 min and then cooling it to 25°C. The resulting mRNA/HEX-PuL complexes from the R19 and R21 libraries were combined (125 µL in total). This mixture was used for the first round of selection without purification.

### *In vitro* selection of anticalins against *cis*-Et-FAzo and *cis*-Ca-FAzo ligands

For the first selection round, an *E. coli* reconstituted cell-free translation mixture (500 µL, with all components listed in **Supplementary Table 5**) containing mRNA/HEX-PuL complexes from the combined R19 and R21 libraries (1 µM in total) was prepared and incubated at 37℃ for 30 min. Subsequently, a solution containing 41.7 µL of 200 mM ethylenediaminetetraacetic acid (EDTA) (pH 8.0), 41.1 µL of RT buffer [46.6 mM Tris-HCl (pH 8.4), 70 mM KCl, 20 mM MgCl_2_, 4.67 mM DTT], and 66.7 µL of 5 mM dNTPs was added and the reaction mixture was incubated at 42℃ for 3 min. Then, 10 µL of 100 µM Lcn.R26 and 27.5 µL of 28.7 µM MMLV reverse transcriptase (RNaseH–, prepared in our lab)^29^ were added and the mixture was incubated at 42℃ for 15 min, generating the anticalin-mRNA/cDNA complex library. The buffer was exchanged for HBST buffer using a Zeba Spin Desalting Column (Thermo Fisher Scientific).

A 12 µL volume of each of the four types of Dynabeads M270—(i) Et-FAzo-beads/– PEG, (ii) Et-FAzo-beads/+PEG, (iii) Ca-FAzo-beads/–PEG, (iv) Ca-FAzo-beads/+PEG—was suspended in HBST buffer containing 345 pmol each of Et-FAzo and Ca-FAzo. This mixed suspension was irradiated with 500 nm light for 30 min using fluorescence spectrometry (FP-8500; JASCO) to convert the azobenzene to its *cis* form. An anticalin-mRNA/cDNA complex library in HBST buffer (690 µL total) was then added to the beads, and the suspension was mixed at 25℃ for 20 min. The beads were washed three times with HBST buffer and resuspended in 690 µL of 1× PCR dNTPs buffer [10 mM Tris-HCl (pH 8.4), 100 mM KCl, 0.1% (v/v) Triton X-100, 2 mM MgSO_4_, 0.2 mM each dNTP]. The suspension was heated at 95℃ for 5 min to elute cDNAs from the beads, and the cDNAs were quantified by SYBR green-based qPCR using T7SD8M2.F44 and Lcn Realtime.R20 as primers. The input anticalin-mRNA/cDNA complex library aliquoted before the selection was also quantified. cDNA recovery (%) was calculated by dividing the amount of recovered cDNA by the amount of input cDNA. The eluted cDNAs were amplified by PCR (14 cycles, 1350 µL total) using T7SD8M2.F44 and G5S-4Gan21-3.R42 as primers. The amplified DNAs were purified by phenol/chloroform extraction and isopropanol precipitation and then finally dissolved in 75 µL of 10 mM Tris-acetate (pH 7.8). A 10 µL aliquot of the resulting DNA pool was transcribed using *in vitro* runoff transcription, and the mRNA was purified by isopropanol precipitation. The mRNA and HEX-PuL complex was prepared (10 µL) by annealing mRNA (4 µM) and HEX-PuL (4 µM) in an annealing buffer, heating the solution to 95°C for 2 min and then cooling it to 25°C. This complex was used for the second selection round without purification.

For the second round, a cell-free translation mix (9 µL) was prepared by adding 2.25 µL of the 4 µM mRNA/HEX-PuL complex and at 37℃ for 30 min. Then, 1.8 µL of 100 mM EDTA (pH 8.0) was added and the mixture was incubated at 42℃ for 3 min. Subsequently, 5.4 µL of 3× RT mix [150 mM Tris-HCl (pH 8.4), 9 mM MgCl_2_, 225 mM KCl, 16.5 mM DTT, 1.5 mM dNTPs, 7.5 µM GSSG3R.R26, 66 mM MgCl_2_, and 3.4 µM MMLV reverse transcriptase] was added and the reaction mixture was incubated at 42℃ for 15 min. The buffer was then exchanged for HBST buffer using a Zeba Spin Desalting Column. A 9 µL TRAP reaction solution was added to 9 µL (suspension volume; the supernatant was removed just before use) of HBY-immobilized M280 beads, and the suspension was mixed at 25℃ for 10 min to remove nonspecific anticalin clones. The supernatant was collected and added again to the HBY-immobilized M280 beads. This procedure was repeated three times. For negative selection, the collected supernatant (9 µL) was added to the HBY-immobilized M270 beads and mixed at 25℃ for 10 min, and the supernatant was recovered for positive selection. The remaining beads were washed three times with HBST buffer and resuspended in 50 µL of 1× PCR dNTPs for real-time PCR analysis. For positive selection, 1 µL of 5 µM each of Et-FAzo-biotin (**6**) and Ca-FAzo-biotin (**7**) in HBST buffer, irradiated with 500 nm (*cis*-forming) light for 30 min (FP-8500; JASCO), was added to 9 µL of the supernatant. The solution was incubated at 25℃ for 10 min. The Et-FAzo-biotin (**6**) and Ca-FAzo-biotin (**7**) were collected using Dynabeads M270 Streptavidin. The beads obtained from positive selection were washed three times with HBST buffer and resuspended in 50 µL of 1× PCR dNTPs. The solution was heated at 95℃ for 5 min to elute cDNAs from the beads, and the cDNAs were quantified by qPCR using T7SD8M2.F44 and Lcn Realtime.R20 as primers. cDNAs obtained from positive selection were amplified by PCR using T7SD8M2.F44 and G5S-4Gan21-3.R42 as primers and then purified by phenol/chloroform extraction and isopropanol precipitation.

From the third to fifth rounds of selection, purified cDNAs were directly added to the TRAP reaction mixture containing T7 RNA polymerase (1 µM) and HEX-PuL (1 µM) (**Supplementary Table 5**). The negative and positive selection procedures following the TRAP reaction were the same as those used in the second round.

For the sixth and seventh selection rounds, the procedures were similar to those for the third round, with the exception of the elution method for cDNAs in positive selection. The beads were resuspended in 16 µL of HBST buffer and irradiated with 410 nm (*trans*-forming) light for 10 min to collect cDNAs encoding anticalins that dissociate from the beads upon *cis→trans* photoisomerization. The supernatant (16 µL) was recovered, and 34 µL of 1.5× PCR dNTPs [15 mM Tris-HCl (pH 8.4), 150 mM KCl, 0.15% (v/v) Triton X-100, 3 mM MgSO_4_, 0.3 mM each dNTP] was added. This solution was used for qPCR and the amplification of cDNAs.

For the eighth selection round, the TRAP reaction volume was scaled to 27 µL. The procedures were the same as in the third round, except that positive selection was performed separately against the two target molecules, Et-FAzo-biotin and Ca-FAzo-biotin in the *cis*-form (without mixing), as well as against their mixture in the *trans*-form. Three solutions were prepared for the positive selection: 5 µM Et-FAzo-biotin irradiated with 500 nm (*cis*-forming) light for 10 min, 5 µM Ca-FAzo-biotin irradiated with 500 nm (*cis*-forming) light for 10 min, and a mixture of 5 µM Et-FAzo-biotin and 5 µM Ca-FAzo-biotin irradiated with 410 nm (*trans*-forming) light for 10 min. One microliter of each of these three solutions was added to 9 µL of the supernatant from the negative selection and used for positive selection, resulting in a total volume of 10 µL per reaction.

### Next-generation sequencing of the cDNA encoding anticalins

Three types of cDNA pools recovered in the eighth round were analysed by next-generation sequencing (Ion Torrent; Thermo Fisher Scientific). The frequency (%) of each sequence was calculated by dividing the number of reads for each sequence by the total number of reads. The *cis*/*trans* ratio was then determined by dividing the frequency of each sequence binding to the *cis*-target molecule by the frequency of the same sequence binding to the *trans*-target molecule. The best anticalin clones against *cis*-Et-FAzo and *cis*-Ca-FAzo ligands were selected based on their frequencies and *cis*/*trans* ratios. Next-generation sequencing results are provided in **Supplementary Table 6**.

### Protein expression and purification

The DNA and amino acid sequences of the AzoTag1 and AzoTag16 fusion proteins used for *in vitro* experiments are listed in the **Supplementary Sequences**. The pQCSoHis plasmid was used as a vector backbone.^72^

For *in vitro* experiments, AzoTag1 was expressed as a C-terminal NusA-His6-tag fusion protein (AzoTag1-NusA-His6) and purified using a method similar to that described previously.^29^ The plasmid encoding the fusion protein, pT5-AzoTag1-NusA-His6 (**Supplementary Figure 23**), was transformed into *E. coli* strain BL21(DE3). The transformants were initially cultured in 5 mL of LB broth containing 100 μg/mL ampicillin and 5% glucose at 37°C. The culture was then transferred to 400 mL of LB broth with the same antibiotics and 5% glucose, and the cells were grown at 37°C until the OD_660_ reached 0.6. The culture was cooled on ice for 30 min, and isopropyl-*β*-D-thiogalactoside (IPTG) was added to a final concentration of 500 μM to induce protein expression. The cells expressing AzoTag1-NusA-His6 were further cultured at 18°C for 24 h.

The cells were harvested by centrifugation, resuspended in buffer A (300 mM AcOK, 1 M KCl, 20 mM HEPES-KOH, 5 mM imidazole, pH 7.8), and disrupted by sonication on ice using an ultrasonic homogenizer (UH-50; SMT). The lysate was cleared by centrifugation and filtered through a sterilized cellulose acetate membrane (pore size: 0.20 µm) (ADVANTEC). The filtered supernatant was subjected to affinity purification using a HisTrap FF column (Cytiva) according to the manufacturer’s protocol on an ÄKTA start system (Cytiva). Bound proteins were washed sequentially with buffer B (300 mM AcOK, 1 M KCl, 20 mM HEPES-KOH, 5 mM imidazole, 2 mM ATP, 10 mM MgSO4, 0.2 mM DTT, pH 7.8) and buffer C (300 mM AcOK, 20 mM HEPES-KOH, 10 mM imidazole, 0.2 mM DTT, pH 7.8), and eluted with buffer D (300 mM AcOK, 500 mM imidazole, 0.2 mM DTT, pH 7.8). The eluted protein solution was dialyzed (MWCO: 5–8 kDa) against buffer E (20 mM HEPES-NaOH, pH 7.5, 150 mM NaCl). The protein concentration was determined by UV spectroscopy using a molar extinction coefficient at 280 nm of 64,455 M^−1^ cm^−1^.

Similarly, AzoTag16 was expressed as a C-terminal NusA-His6-tag fusion protein (AzoTag16-NusA-His6) using the plasmid pT5-AzoTag16-NusA-His6 (**Supplementary Figure 23**) and purified in the same manner. Its concentration was determined using a molar extinction coefficient at 280 nm of 63,425 M^−1^ cm^−1^.

## Isothermal titration calorimetry

The binding affinities and binding thermodynamic parameters of AzoTag1 and AzoTag16 for their respective ligands were measured using isothermal titration calorimetry with a MicroCal iTC200 instrument. All experiments were performed in the same buffer used for dialysis (20 mM HEPES-NaOH, pH 7.5, 150 mM NaCl). For the titration of the AzoTag1/Et-FAzo pair, the cell was loaded with 80 µM AzoTag1, and the syringe was filled with 960 µM Et-FAzo (either the pure *trans*- or *cis*-form, with 0.5% DMSO). Similarly, for the AzoTag16/Ca-FAzo pair, the cell was loaded with 50 µM AzoTag16, and the syringe was filled with 600 µM Ca-FAzo (also in either the pure *trans*- or *cis*-form, with 0.8% DMSO).

An initial injection of 0.5 μL of ligand was followed by 30 injections of 1.0 μL at 2-min intervals at 25°C, with the syringe stirred at 1,000 rpm. Data were analysed using ORIGIN software (MicroCal) based on a one-site binding model to extract thermodynamic parameters, including *K*_a_ (1/*K*_d_), ΔH, and stoichiometry (*N*). Values for Δ*G* and Δ*S* were calculated using the equations Δ*G* = –RTln*K*_a_ and Δ*G* = Δ*H* – TΔ*S*. Each experiment was performed in triplicate to determine the mean and standard deviation for each measurement.

### Plasmids used in mammalian cell experiments

The domain structures of the fusion proteins constructed in this study are summarized in **Supplementary Figure 23**, and their DNA and amino acid sequences are listed in the **Supplementary Sequences**. The vector backbones used were pmCherry-N1 (Clontech), pmCherry-C1 (Clontech), and pCAGGS (provided by Jun-ichi Miyazaki, Osaka University).^73^ The following plasmids were either purchased or kindly provided by the following companies or researchers and were used directly or as PCR templates to construct expression plasmids: pFN22K (HaloTag7 CMV*d1* Flexi vector) from Promega; pmiRFP670-N1 (Addgene plasmid #79987)^74^ and pmiRFP680-N1 (Addgene plasmid #136557)^75^ from Vladislav Verkhusha (Albert Einstein College of Medicine); pCAGGS-HaloTag-KRas4B(CT) (Addgene plasmid #214276),^72^ pPBpuro-cRaf-mNG-eDHFR(69K6) (Addgene plasmid #172107),^49^ and pPBpuro-mNG-eDHFR(69K6)-p85iSH2 (Addgene plasmid #172103)^49^ from our group; SP-dCas9-VPR (Addgene plasmid #63798)^76^ from George Church (Harvard University); pPBpuro-TRE-mAG^56^ and pU6-sgRNA^56^ from Hirohide Saito (The University of Tokyo); and EMTB-mScarletI (Addgene plasmid #137801)^77^ and FRB-ECFP(W66A)-Giantin (Addgene plasmid #67903)^78^ from Dorus Gadella (University of Amsterdam). DNA sequences encoding organelle-targeting signals for mitochondria (TOMM20), the nucleus (NLS), and the endoplasmic reticulum (Cb5) were synthesized by a commercial vendor. All expression plasmids were generated using standard cloning procedures and the NEBuilder HiFi DNA assembly system (New England Biolabs). All PCR-amplified sequences were verified by DNA sequencing.

### Cell culture

HeLa and Cos-7 cells were obtained from the Cell Resource Center for Biomedical Research at the Institute of Development, Aging and Cancer, Tohoku University. All cells were cultured in Dulbecco’s modified Eagle’s medium (DMEM) supplemented with 10% heat-inactivated fetal bovine serum (FBS), 100 U/mL penicillin, and 100 µg/mL streptomycin [DMEM(+)] at 37°C under a humidified 5% CO_2_ atmosphere. For transient expression experiments, cells were transfected using 293fectin (Thermo Fisher Scientific) according to the manufacturer’s instructions.

### Live-cell imaging

Confocal fluorescence imaging was conducted using an Olympus IX83/FV3000 confocal laser-scanning microscope equipped with a UPLSAPO60XO/1.35 NA oil objective (Olympus), a Z drift compensator system (IX3-ZDC2, Olympus), and a stage-top incubator (Tokai Hit). Fluorescence images were captured using lasers at 488 nm (for mAzamiGreen), 561 nm (for mCherry, mScarlet-I, and tetramethylrhodamine), and 640 nm (for miRFP670 and miRFP680) at 37°C under a 5% CO_2_ atmosphere. For all imaging experiments, phenol red- and serum-free DMEM supplemented with 100 U/mL penicillin and 100 µg/mL streptomycin [DMEM(–)] was used. Fluorescence images were analysed using the Fiji distribution package of ImageJ.^79^

### Stock preparation

For cell experiments, Et-FAzo-HTL (**8**), Ca-FAzo-HTL (**9**), and Ca(AM)-FAzo-HTL (**10**) were each dissolved in DMSO to prepare 20 mM stock solutions. The solutions were then irradiated with 400 nm light for 5 min to convert the FAzo-HTL ligands to their *trans*-form, protected from light, and stored at –20°C.

### CAPA assay

HeLa cells were plated at 1.0 × 10^5^ cells/dish in 35-mm glass-bottom dishes (IWAKI) and cultured for 24 h at 37°C in 5% CO_2_. The cells were transfected with pCMV-HaloTag-miRFP670 (1.0 µg) using 293fectin. After 12 h, 25 µM biliverdin, a cofactor for the fluorescent protein miRFP670,^80,81^ was added to the medium. Following an additional 12 h incubation, the medium was replaced with DMEM(–), and all subsequent steps were performed in the dark. FAzo ligands [10 µM Et-FAzo-HTL (**8**), 10 µM Ca-FAzo-HTL (**9**), and 5 µM Ca(AM)-FAzo-HTL (**10**)] were added and the medium was incubated at 37°C for 30 min. Unbound ligands were removed by washing the cells twice with fresh DMEM(–). The cells were then stained with 5 µM TMR-HTL (HaloTag TMR ligand; Promega) at 37°C for 30 min. After washing, fluorescence images were acquired using confocal microscopy.

### Evaluation of AzoTag1/Et-FAzo and AzoTag16/Ca-FAzo pairs in live cells

HeLa cells were plated at 1.0 × 10^5^ cells/dish in 35-mm glass-bottom dishes (IWAKI) and cultured for 24 h at 37°C in 5% CO_2_. To evaluate the AzoTag1/Et-FAzo pair, the cells were transfected with pCAGGS-AzoTag1-miRFP670-NES-P2A-HaloTag-KRas4B(CT) (1.0 µg) using 293fectin. After 12 h, 25 µM biliverdin, a cofactor for the fluorescent protein miRFP670, was added to the medium. Following an additional 12 h incubation, the medium was replaced with DMEM(–), and all subsequent steps were performed in the dark. FAzo ligands [10 µM Et-FAzo-HTL and 5 µM Ca(AM)-FAzo-HTL] were added to the medium and the dishes were incubated at 37°C for 30 min. Unbound ligands were removed by washing the cells twice with fresh DMEM(–), and the cells were imaged using time-lapse confocal fluorescence microscopy. Blue (483 nm) and violet (400 nm) light were delivered from above the dish using the xenon light source described above, with bandpass filters of 483 ± 16 nm (6.0 mW/cm²) and 400 ± 20 nm (5.0 mW/cm²), respectively.

The AzoTag16/Ca-FAzo pair was evaluated in the same manner, except that cells were transfected with pCAGGS-AzoTag16-miRFP670-NES-P2A-HaloTag-KRas4B(CT) (1.0 µg).

For the evaluation of binding and dissociation kinetics, the entire field of view was irradiated using 405 nm [laser intensity: 5.0 mW; scan time: 1.2 s per frame (50 × 50 µm)] and 488 nm [laser intensity: 6.0 mW; scan time: 1.2 s per frame (50 × 50 µm)] lasers incorporated into the confocal laser scanning microscope.

### Reversible dual-light control of synthetic AzoTag16-fused cRaf

HeLa cells were plated at 1.0 × 10^5^ cells/dish in 35-mm glass-bottom dishes (IWAKI) and cultured for 24 h at 37°C in 5% CO_2_. The cells were cotransfected with pCAGGS-AzoTag16-miRFP680-cRaf (0.4 µg), pCAGGS-HaloTag-KRas4B(CT) (0.4 µg), and pCAGGS-MEK1-P2A-mScarlet-I-ERK(K57R) (0.2 µg) at a 2:2:1 ratio using 293fectin. After 12 h, 25 µM biliverdin was added to the medium. Following an additional 12 h incubation, the medium was replaced with DMEM(–), and all subsequent steps were performed in the dark. Ca(AM)-FAzo-HTL (5 µM) was added to the medium and the dishes were incubated at 37°C for 30 min. Unbound ligand was removed by washing the cells twice with fresh DMEM(–), and the cells were imaged using time-lapse confocal fluorescence microscopy. Blue (483 nm) and violet (400 nm) light were delivered from above the dish using the xenon light source described above. Control experiments were performed in the same manner.

### Reversible dual-light-control of synthetic AzoTag16-fused PI3K

HeLa cells were plated at 1.0 × 10^5^ cells/dish in 35-mm glass-bottom dishes (IWAKI) and cultured for 24 h at 37°C in 5% CO_2_. The cells were cotransfected with pCAGGS-AzoTag16-miRFP680-p85iSH2 (0.4 µg), pCAGGS-HaloTag-KRas4B(CT)^72^ (0.4 µg), and pCAGGS-mScarlet-I-PH_Akt_ (0.2 µg) at a 2:2:1 ratio using 293fectin. After 12 h, 25 µM biliverdin was added to the medium. Following an additional 12 h incubation, the medium was replaced with DMEM(–), and all subsequent steps were performed in the dark. Ca(AM)-FAzo-HTL (5 µM) was added to the medium and the dishes were incubated at 37°C for 30 min. Unbound ligand was removed by washing the cells twice with fresh DMEM(–), and the cells were imaged using time-lapse confocal fluorescence microscopy. Blue (483 nm) and violet (400 nm) light were delivered from above the dish using the xenon light source described above. Control experiments were performed in the same manner.

### Dual-light control of spatially localized lipid signaling

A 1.0 mg/mL fibronectin solution (Corning) was diluted to 50 µg/mL in Dulbecco’s phosphate-buffered saline (D-PBS). *Two hundred microliters of the diluted solution* was spread over the 12-mm glass surface of 35-mm glass-bottom dishes (IWAKI) and incubated at room temperature for 1 h. The solution was removed and the dish was dried at room temperature for 1 h. Once dried, the dish was washed with D-PBS and prepared for subsequent experiments.

Cos-7 cells were plated at 0.5 × 10^5^ cells/dish on fibronectin-coated 35-mm glass-bottom dishes and incubated for 24 h at 37°C in 5% CO_2_. The cells were cotransfected with pCAGGS-AzoTag16-miRFP680-p85iSH2 (0.4 µg), pCAGGS-HaloTag-KRas4B(CT)^72^ (0.4 µg), and pCMV-Lifeact-m-Scarlet-I (0.2 µg) in a 2:2:1 ratio using 293fectin. After 12 h, 25 µM biliverdin was added to the medium. Following another 12 h of incubation, the medium was replaced with DMEM(–), and all subsequent steps were carried out in the dark. Ca(AM)-FAzo-HTL (5 µM) was added to the medium and incubated at 37°C for 60 min. Unbound ligand was washed away by rinsing the cells twice with fresh DMEM(–), and the cells were imaged using time-lapse confocal fluorescence microscopy. For local photoactivation, selected regions were illuminated by scanning with a 488 nm laser [laser intensity: 6.0 mW; scan time: 30 ms per frame; covering a circular area with a 5 µm diameter]. For photoinactivation, the entire field of view was irradiated by scanning with a 405 nm laser (laser intensity: 5.0 mW; scan time: 1.2 s per frame; covering 100 × 110 µm). Control experiments were conducted in the same manner.

### Light-controlled gene expression using an AzoTag16-based CRISPR-activation system

HeLa cells were plated at 1.0 × 10^5^ cells/dish in 35-mm glass-bottom dishes and cultured for 24 h at 37°C in 5% CO_2_. The cells were cotransfected with pCMV-dCas9-AzoTag16 (0.125 µg), pCMV-HaloTag-VPR (0.125 µg), TRE-pCMV_mini_-mAzamiGreen^56^ (0.125 µg), and pU6-sgRNA^56^ (0.125 µg) in a 1:1:1:1 ratio using 293fectin. Additionally, miRFP680 was coexpressed as an expression marker using pmiRFP680-N1^75^ (0.1 µg). After 12 h, 25 µM biliverdin was added to the medium. Following another 12 h of incubation, the medium was replaced with DMEM(–), and all subsequent steps were performed in the dark. Pre-light-irradiation images of mAzamiGreen fluorescence were acquired before light illumination. Ca(AM)-FAzo-HTL (5 µM) was then added to the medium and the dishes were incubated at 37°C for 30 min. Unbound ligand was removed by washing the cells twice with fresh DMEM(–), after which the entire dish was irradiated with blue light (483 nm) for 2 min using the xenon light source described above. The culture medium was then replaced with DMEM(+) and incubated for 24 h at 37°C, after which post-light-irradiation images of mAzamiGreen fluorescence were acquired. Control experiments were conducted in the same manner.

### Light-controlled protein relocalization to various organelles

HeLa cells were plated at 1.0 × 10^5^ cells/dish in 35-mm glass-bottom dishes (IWAKI) and cultured for 24 h at 37°C. The cells were cotransfected with pCAGGS-AzoTag16-miRFP670 (0.5 µg) and one of the following plasmids (0.5 µg) at a 1:1 ratio using 293fectin: pCMV-TOMM20-mCherry-HaloTag (targeting mitochondria), pCMV-mCherry-HaloTag-NLS (targeting the nucleus), pCMV-HaloTag-mCherry-Cb5 (targeting the ER), pCMV-EMTB-HaloTag-mCherry (targeting microtubules), and pCMV-HaloTag-mCherry-Giantin (targeting the Golgi). After 12 h, 25 µM biliverdin was added to the medium. Following another 12 h of incubation, the medium was replaced with DMEM(–), and all subsequent steps were performed in the dark. Ca(AM)-FAzo-HTL (5 µM) was added to the medium and the dish was incubated at 37°C for 30 min. Unbound ligand was removed by washing the cells twice with fresh DMEM(–), and the cells were imaged using time-lapse confocal fluorescence microscopy. Blue (483 nm) and violet (400 nm) light were delivered from above the dish using the xenon light source described above.

