## Supplementary Information for "*De novo* chemo-optogenetics through rational small-molecule design and TRAP display"

[1]*Department of Life Science and Applied Chemistry, Graduate School of Engineering, Nagoya Institute of Technology, Nagoya 466-8555, Japan*
[2]*Department of Nanopharmaceutical Sciences, Nagoya Institute of Technology, Nagoya 466-8555, Japan*
[3]*Department of Biomolecular Engineering, Graduate School of Engineering, Nagoya University, Nagoya 464-8603, Japan*
[4]*PRESTO, Japan Science and Technology Agency (JST), Kawaguchi, Saitama 332-0012, Japan*
[5]*Medical Device Development and Regulation Research Center, School of Engineering, The University of Tokyo, Tokyo 113-8654, Japan*
[6]*Department of Bioengineering, School of Engineering, The University of Tokyo, Tokyo 113-8654, Japan*
[7]*Department of Chemistry and Bioengineering, School of Engineering, The University of Tokyo, Tokyo 113-8654, Japan*
[8]*Institute of Nano-Life Systems, Institutes of Innovation for Future Society, Nagoya University, Nagoya 464-8603, Japan*
[9]*Research Institute for Quantum and Chemical Innovation, Institutes of Innovation for Future Society, Nagoya University, Nagoya 464-8603, Japan*
[10]*OptoBioTechnology Research Center, Nagoya Institute of Technology, Nagoya 466-8555, Japan*
[#]*These authors contributed equally to this work.*

#### Supplementary Figures

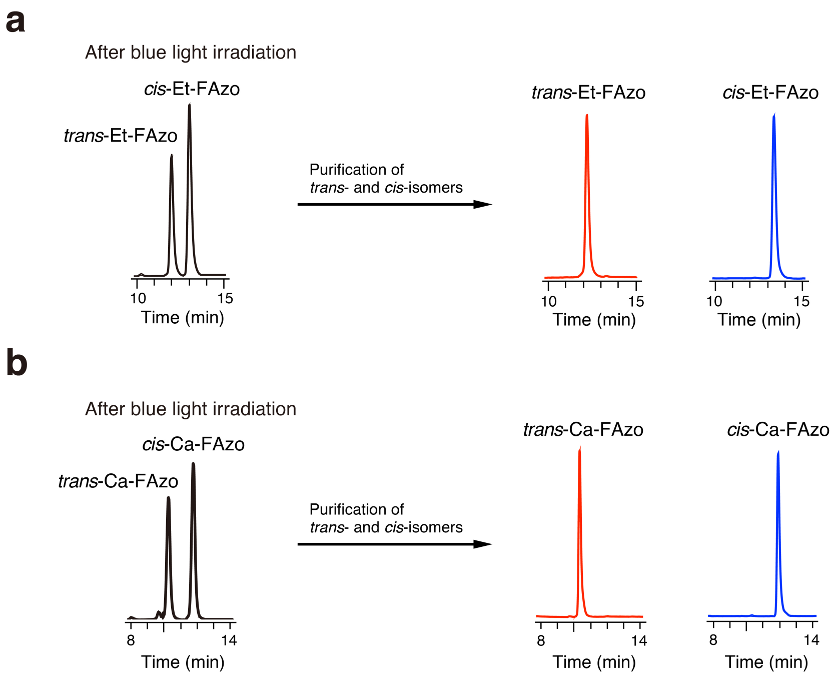


**Supplementary Figure 1. Purification of *trans*- and *cis*-isomers of Et-FAzo and Ca-FAzo ligands.** (**a,b**) Chromatograms recorded before and after purification of the *trans*- and *cis*-forms of Et-FAzo (**a**) and Ca-FAzo (**b**). Solutions of Et-FAzo and Ca-FAzo in MeCN/$H_2O$ (1:1) were irradiated at 483 nm for 5 min and then separated by reversed-phase HPLC on a C4 column for Et-FAzo or an ODC column for Ca-FAzo. Elution was performed with a linear gradient from 40% to 45% MeCN/$H_2O$ containing 0.1% trifluoroacetic acid (TFA) for Et-FAzo and from 40% to 42% MeCN/$H_2O$ containing 0.1% TFA for Ca-FAzo over 30 min. Detection was performed at 440 nm for Et-FAzo and 442 nm for Ca-FAzo.

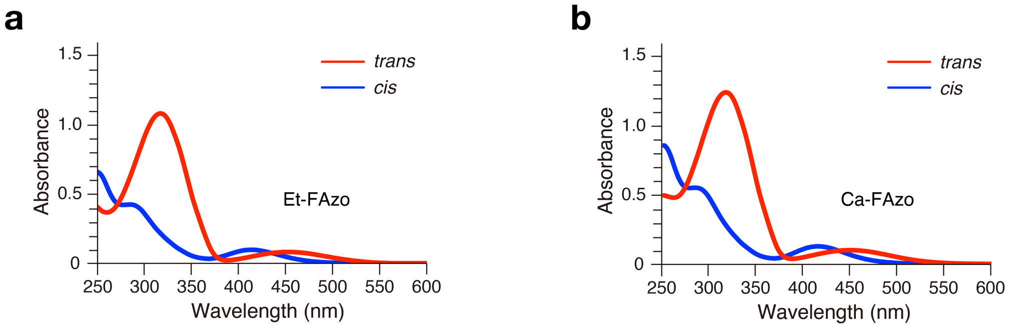


**Supplementary Figure 2. Absorption properties of *trans*- and *cis*-isomers of Et-FAzo and Ca-FAzo ligands. (a,b)** Absorption spectra of purified *trans*- (red) and *cis*- (blue) forms of Et-FAzo (**a**) and Ca-FAzo (**b**) (50 µM) in 20 mM HEPES, 150 mM NaCl, pH 7.5, with 0.5% DMSO for Et-FAzo or 0.8% DMSO for Ca-FAzo.

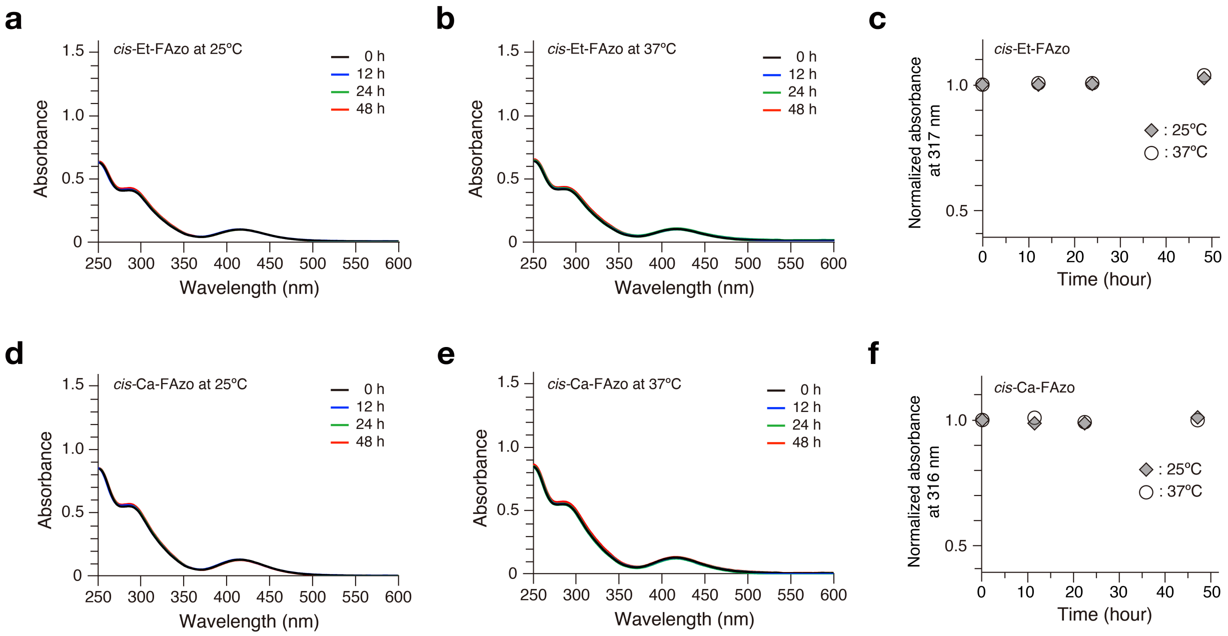


**Supplementary Figure 3. Thermal stability of *cis*-Et-FAzo and *cis*-Ca-FAzo ligands.**

(**a**–**c**) Evaluation of *cis*-Et-FAzo. (**d**–**f**) Evaluation of *cis*-Ca-FAzo. Absorption spectra of *cis*-Et-FAzo and *cis*-Ca-FAzo solutions (50 µM) in the buffer described below were measured after incubation at 25°C (**a**,**d**) or 37°C (**b**,**e**) in the dark for the indicated times. The normalized absorbances at 317 nm for Et-FAzo and 316 nm for Ca-FAzo are plotted as a function of incubation time (**c**,**f**).

Buffer: 20 mM HEPES, 150 mM NaCl, pH 7.5, with 0.5% DMSO for Et-FAzo or 0.8% DMSO for Ca-FAzo.

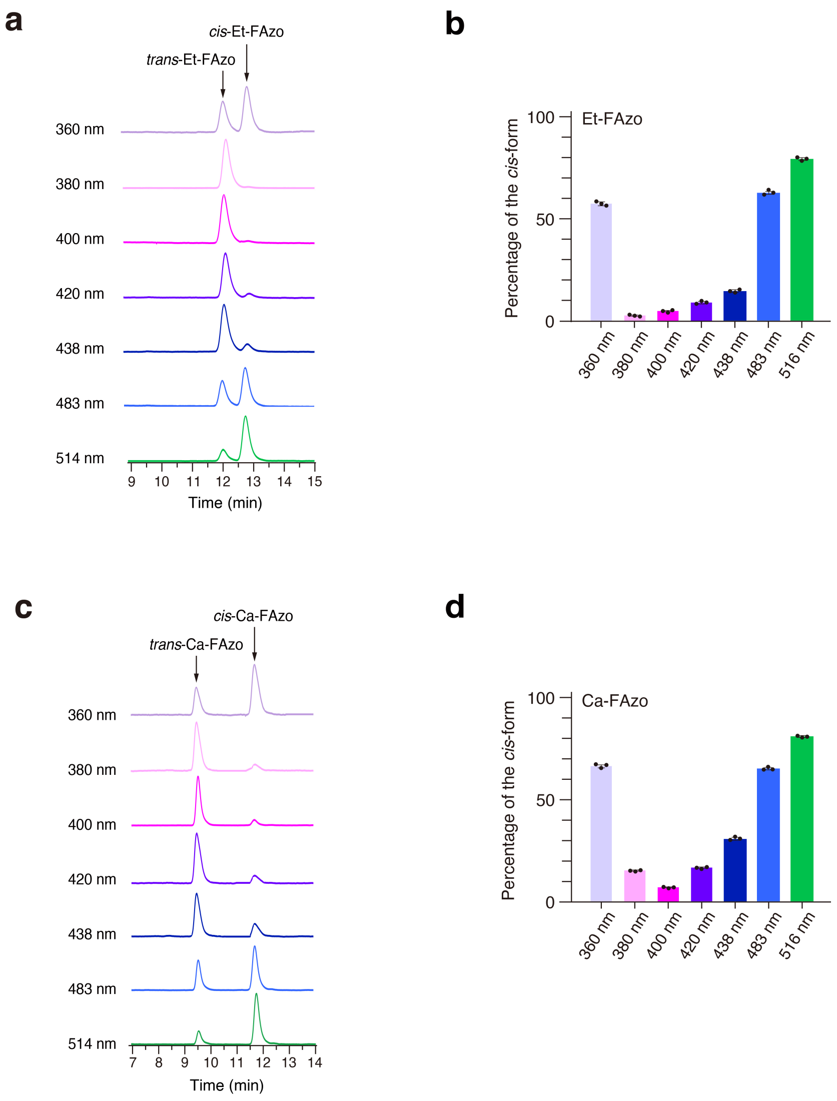


**Supplementary Figure 4. Measurement of photostationary states of Et-FAzo and Ca-FAzo ligands.** (**a**,**b**) Evaluation of Et-FAzo. (**c**,**d**) Evaluation of Ca-FAzo. *trans*-Et-FAzo and *trans*-Ca-FAzo solutions (50 µM) in the buffer described below were irradiated with the indicated wavelengths of light for 5 min. The solutions were separated by reversed-phase HPLC (**a**,**c**) as described in **Supplementary Figure 1**. Detection was performed at 440 nm for Et-FAzo and 442 nm for Ca-FAzo, corresponding to the isosbestic points of the *trans*- and *cis*-FAzo ligands. The percentages of the *cis*-form ($PSS_{cis}$) were determined by integrating the peak area ratios of the *trans*- and *cis*-forms and are presented as the mean ± SD from three independent experiments per condition (**b**,**d**). The results are also summarized in **Supplementary Table 1**.

Buffer: 20 mM HEPES, 150 mM NaCl, pH 7.5, with 0.5% DMSO for Et-FAzo or 0.8% DMSO for Ca-FAzo.

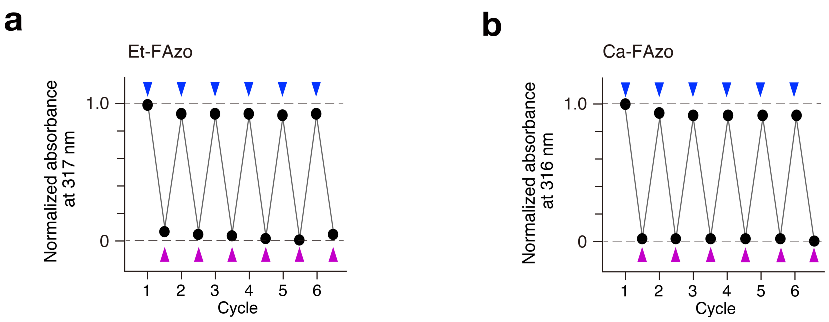


**Supplementary Figure 5. Repetitive photo-switching properties of Et-FAzo and Ca-FAzo ligands.** Absorption spectra of *trans*-Et-FAzo (**a**) and *trans*-Ca-FAzo (**b**) solutions (50 µM) in the buffer described below were recorded before and after repeated irradiation with 483 nm (blue) and 400 nm (violet) light for 5 min. The normalized absorbances at 317 nm for Et-FAzo and 316 nm for Ca-FAzo are plotted as a function of cycle number.

Buffer: 20 mM HEPES, 150 mM NaCl, pH 7.5, with 0.5% DMSO for Et-FAzo or 0.8% DMSO for Ca-FAzo.

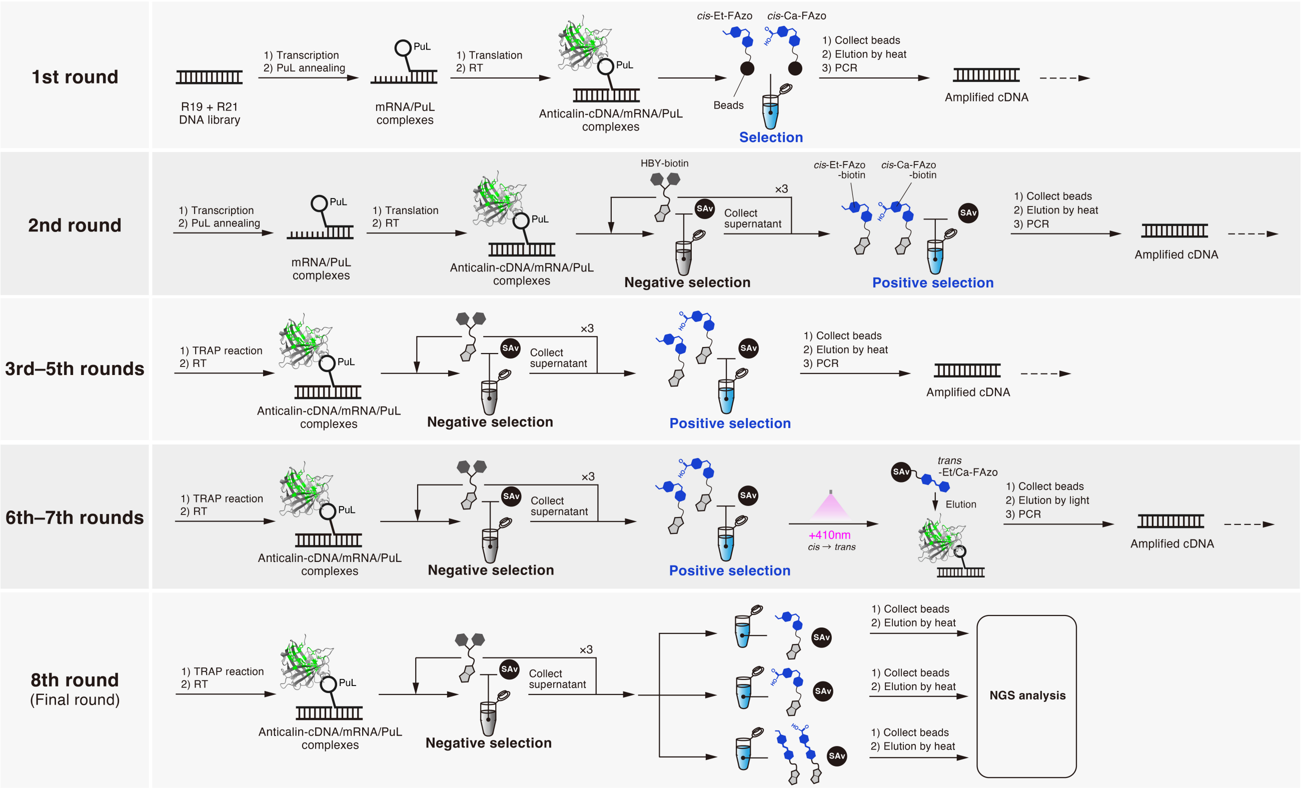


**Supplementary Figure 6. Schematic illustration of the overall *in vitro* selection procedures.**

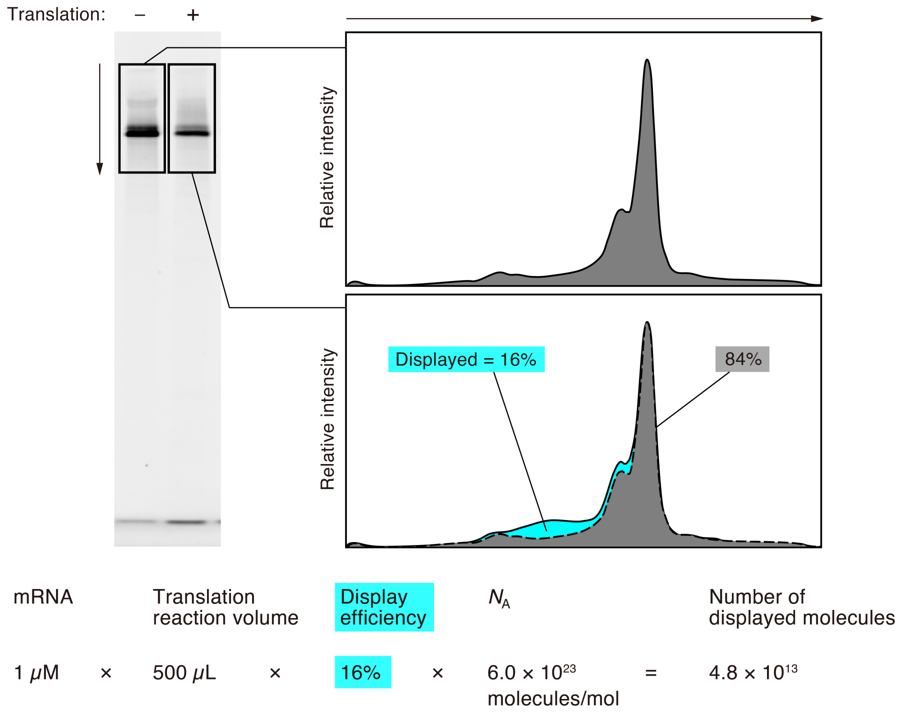


**Supplementary Figure 7. Calculation of the number of anticalin molecules displayed in the first selection round.**

mRNA/HEX-PuL complexes were analysed before and after the translation reaction using urea-SDS-PAGE followed by fluorescence imaging. The relative volume of the bands corresponding to the mRNA/PuL complexes, both with and without anticalin, was calculated (16%). The number of anticalin molecules displayed was calculated using the formula shown at the bottom (4.8 × $10^{13}$ molecules).

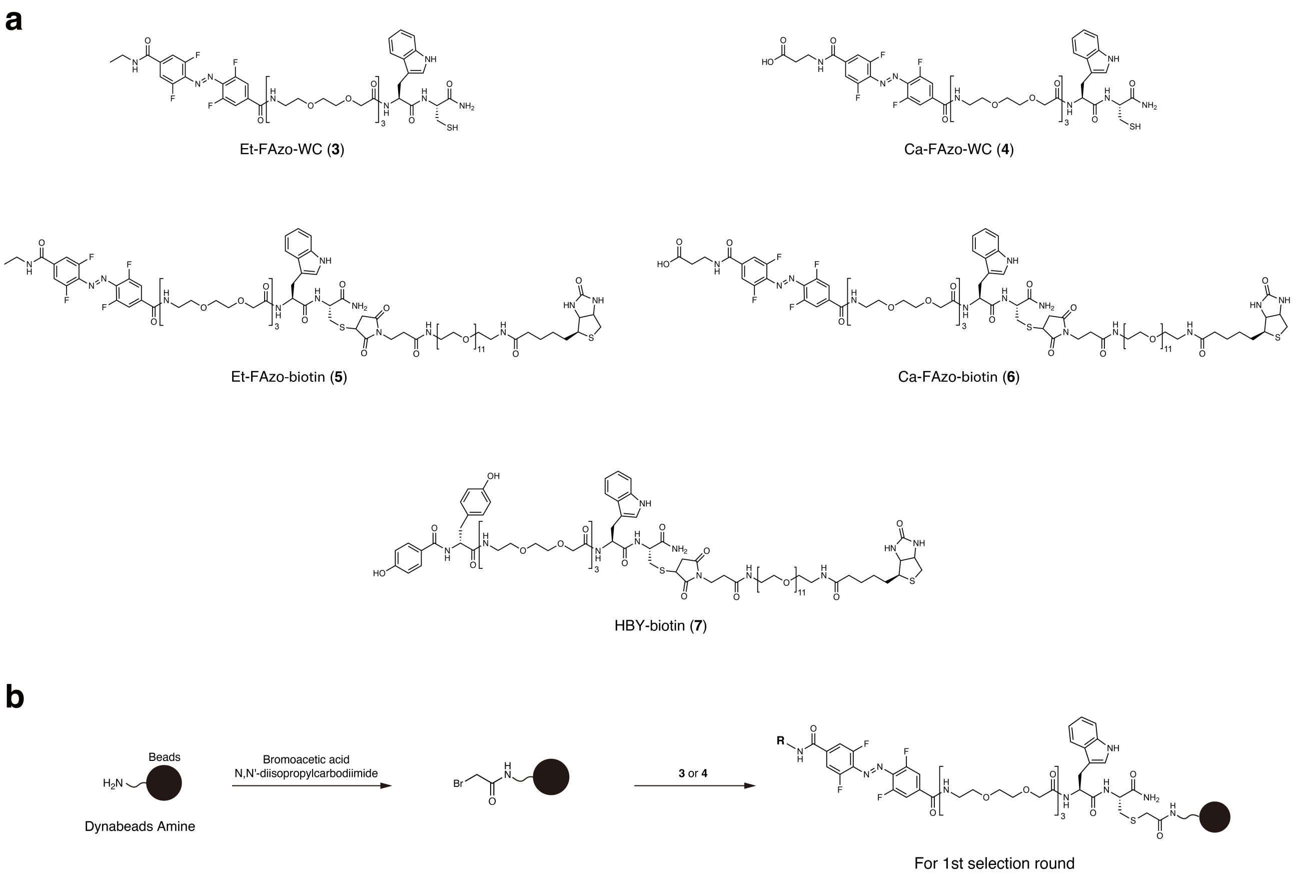


**Supplementary Figure 8. Materials used to prepare beads for *in vitro* selection.**
(**a**) Chemical structures of FAzo ligand derivatives and control compounds used for bead preparation: Et-FAzo-WC (**3**), Ca-FAzo-WC (**4**), HBY-biotin (**5**), Et-FAzo-biotin (**6**), and Ca-FAzo-biotin (**7**). (**b**) Synthetic route for preparing FAzo ligand-immobilized beads used in the first selection round.

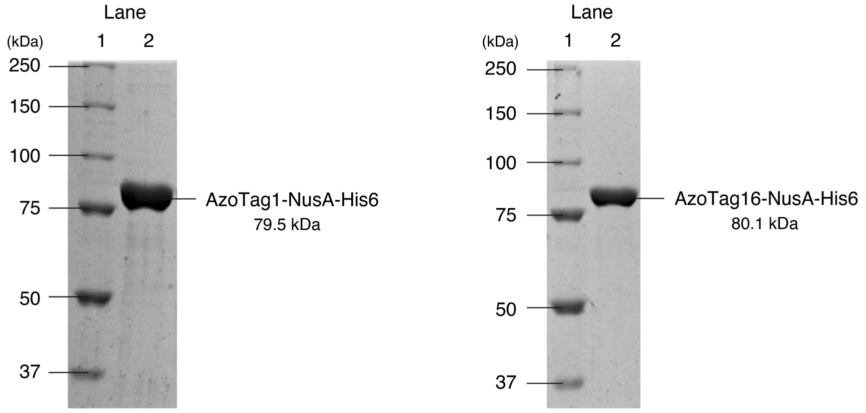


**Supplementary Figure 9. SDS-PAGE analysis of recombinant AzoTag1-NusA-His6 and AzoTag16-NusA-His6 proteins used in ITC experiments.**

Coomassie Brilliant Blue-stained gel images. Lane 1 contains molecular weight markers, while Lane 2 contains purified AzoTag1-NusA-His6 (left) and AzoTag16-NusA-His6 (right).

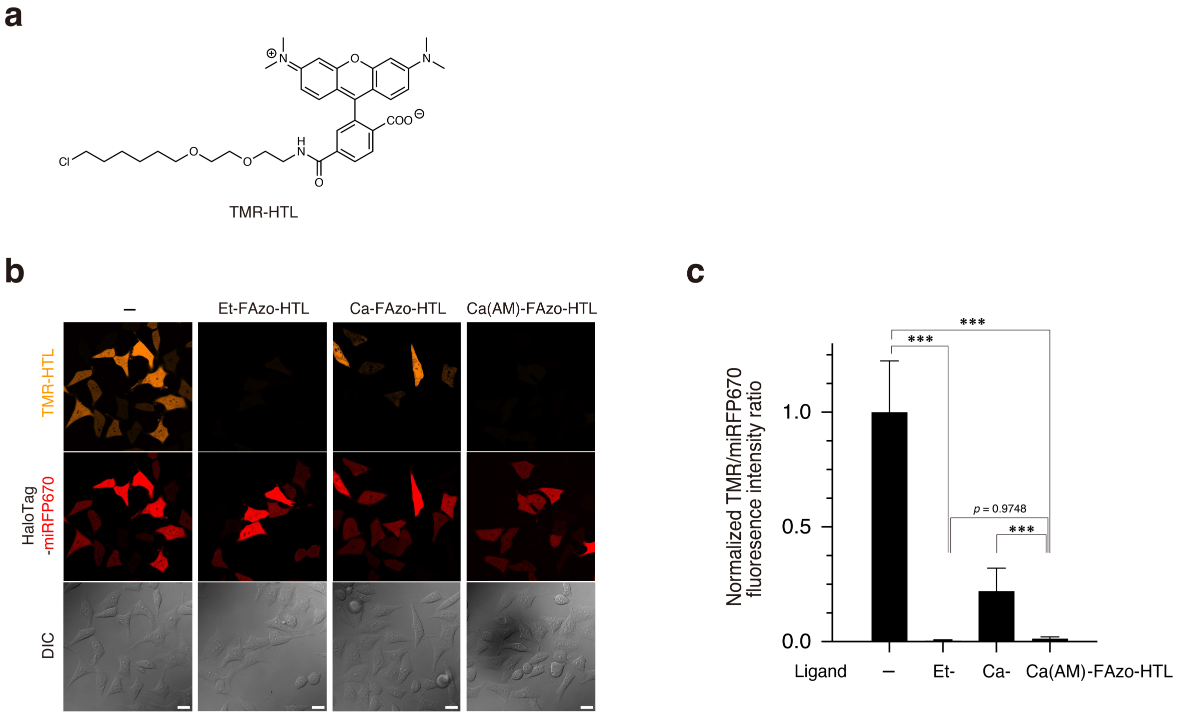


**Supplementary Figure 10. Evaluation of cell membrane permeability of Et-FAzo-HTL, Ca-FAzo-HTL, and Ca(AM)-FAzo-HTL using the CAPA assay.**

(**a**) Chemical structure of TMR-HTL (commercially available as HaloTag TMR ligand; Promega). (**b**) Confocal fluorescence images of HaloTag-miRFP670-expressing HeLa cells after treatment with FAzo ligands and TMR-HTL. The leftmost image shows cells without dimerizer treatment, while the three images on the right show cells treated with 10 $\mu$M Et-FAzo-HTL, 10 $\mu$M Ca-FAzo-HTL, and 5 $\mu$M Ca(AM)-FAzo-HTL. Scale bars, 20 $\mu$m. (**c**) Quantification of the normalized fluorescence intensity ratio between TMR and miRFP670 signals. Data are presented as the mean ± SD (n = 30 cells from three independent experiments per condition). ***$p < 0.001$, Student's $t$-test. For details of the CAPA assay, see Ref. S1.

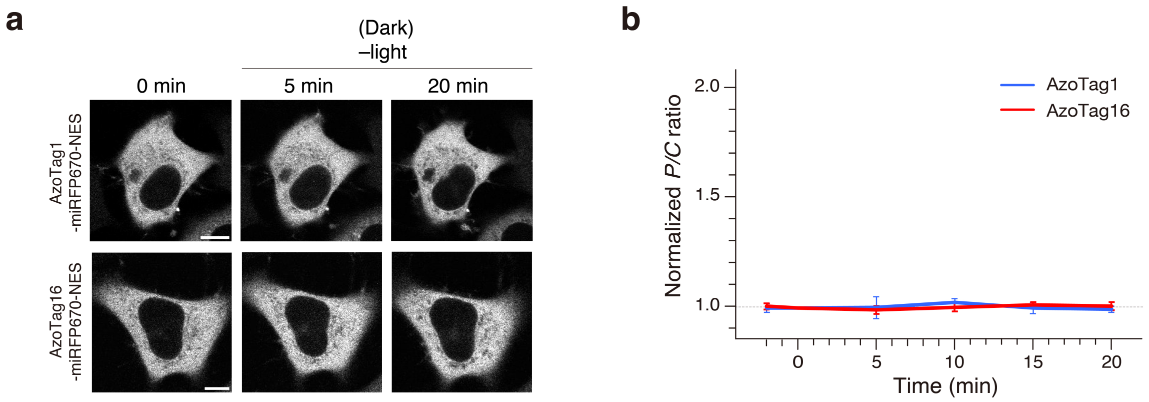


**Supplementary Figure 11. Localization changes of AzoTag1 and AzoTag16 in the dark.**
(**a**) Representative time-lapse confocal fluorescence images of HeLa cells coexpressing AzoTag1-miRFP670-NES with Et-FAzo-HaloTag$^{PM}$ (top) or AzoTag16-miRFP670-NES with Ca-FAzo-HaloTag$^{PM}$ (bottom) in the absence of light irradiation. Images were acquired at the indicated time points. Scale bars, 10 µm. (**b**) Quantification of protein translocation efficiencies. The normalized *P/C* ratios of AzoTag1-miRFP670-NES (blue) and AzoTag16-miRFP670-NES (red) are presented as the mean ± SD as a function of incubation time (n = 8 cells from three independent experiments per construct).

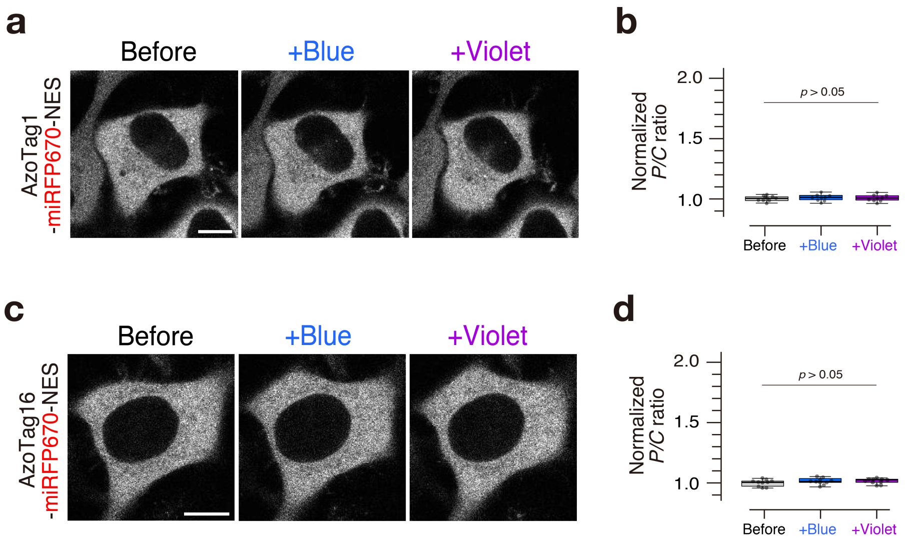


**Supplementary Figure 12. Light-induced localization changes of AzoTag1 and AzoTag16 without FAzo-HTL ligand treatment.**

(**a**,**c**) Representative confocal fluorescence images of HeLa cells coexpressing AzoTag1-miRFP670-NES with $\text{HaloTag}^{\text{PM}}$ (**a**) or AzoTag16-miRFP670-NES with $\text{HaloTag}^{\text{PM}}$ (**c**) in the absence of FAzo-HTL ligand treatment. Images were acquired before and 2 min after irradiation with blue light (483 nm, "+Blue") and violet light (400 nm, "+Violet"). Scale bars, 10 µm. (**b**,**d**) Quantification of protein translocation efficiencies. The normalized *P/C* ratios of AzoTag1-miRFP670-NES (**b**) and AzoTag16-miRFP670-NES (**d**) are presented as box plots (n = 10 cells from three independent experiments per condition). $p > 0.05$, Student's t-test.

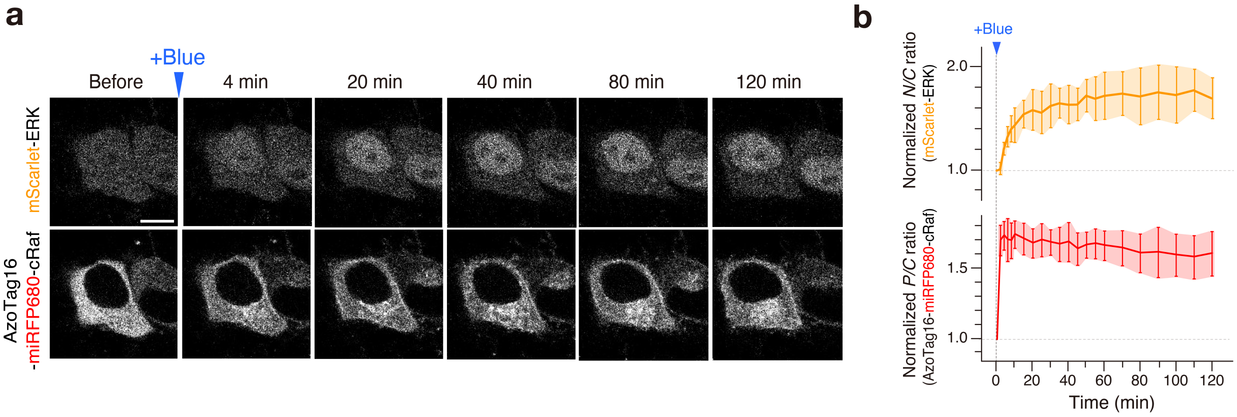


**Supplementary Figure 13. Sustained ERK activation induced by single blue-light irradiation.**
(**a**) Representative time-lapse confocal fluorescence images of HeLa cells coexpressing AzoTag16-miRFP680-cRaf, Ca-FAzo-HaloTag$^{PM}$, and mScarlet-ERK before and after a single irradiation with blue light (483 nm). Images were acquired at the indicated time points. Scale bar, 10 µm. For time-lapse movies, see **Supplementary Video 3**. (**b**) Time course of cRaf translocation and ERK activation. The normalized *P/C* ratios of AzoTag16-miRFP680-cRaf are plotted as a function of time to evaluate cRaf translocation (bottom). The normalized *N/C* ratios of mScarlet-ERK are plotted as a function of time to assess ERK activity (top). Arrowheads indicate light irradiation at 483 nm. Data are presented as the mean ± SD (n = 8 cells from three independent experiments).

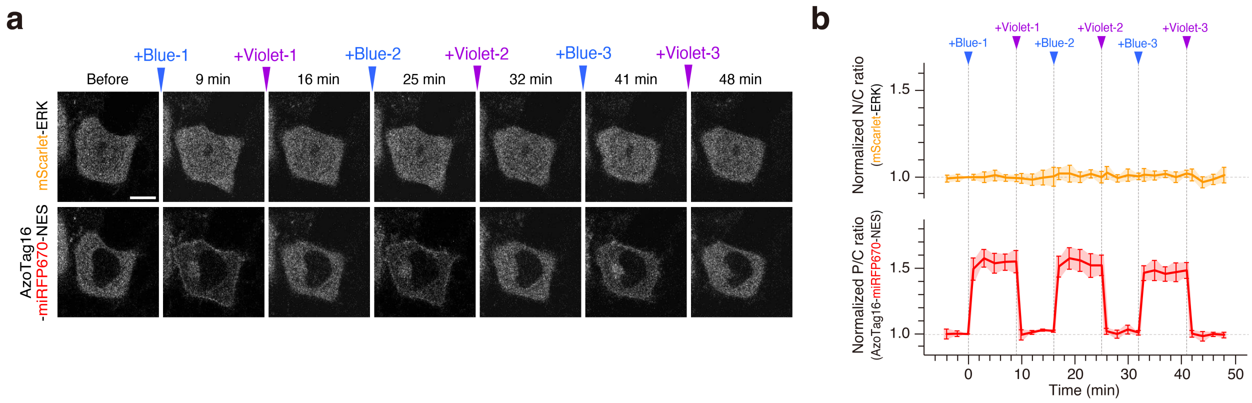


**Supplementary Figure 14. Light control of AzoTag16-miRFP670-NES without cRaf.**

(**a**) Representative time-lapse confocal fluorescence images of a HeLa cell coexpressing AzoTag16-miRFP670-NES lacking cRaf, Ca-FAzo-HaloTag$^{PM}$, and mScarlet-ERK before and after sequential irradiation with blue light (483 nm, "+Blue") and violet light (400 nm, "+Violet"). Images were acquired at the indicated time points. Scale bar, 10 µm. (**b**) Time course of cRaf translocation and ERK activation. The normalized *P/C* ratios of AzoTag16-miRFP680-cRaf are plotted as a function of time to evaluate cRaf translocation (bottom). The normalized *N/C* ratios of mScarlet-ERK are plotted as a function of time to assess ERK activity (top). Arrowheads indicate irradiation with 483 nm (blue) and 400 nm (violet) light. Data are presented as the mean ± SD (n = 8 cells from three independent experiments).

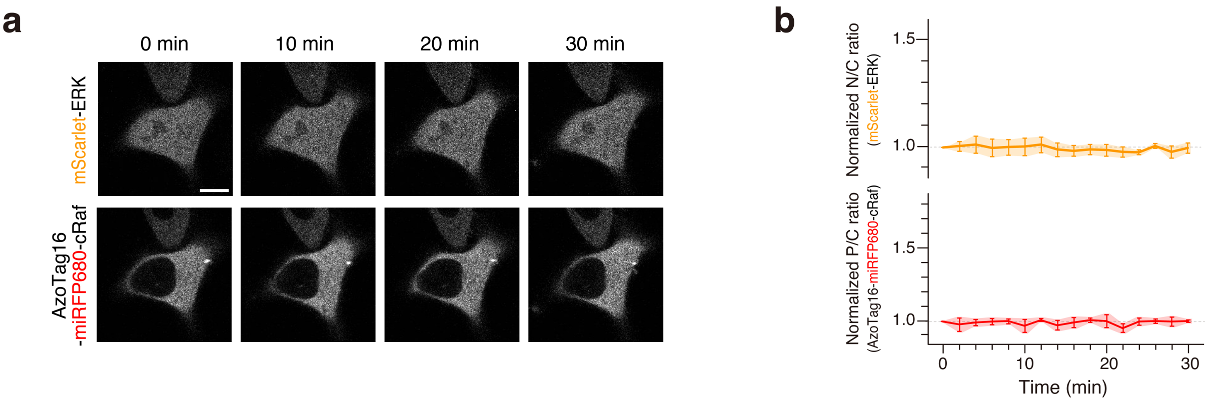


**Supplementary Figure 15. Localization and activity changes of AzoTag16-miRFP680-cRaf in the dark.** (**a**) Representative time-lapse confocal fluorescence images of a HeLa cell coexpressing AzoTag16-miRFP680-cRaf, Ca-FAzo-HaloTag$^{PM}$, and mScarlet-ERK in the absence of light irradiation. Images were acquired at the indicated time points. Scale bar, 10 µm. (**b**) Time course of cRaf translocation and ERK activation. The normalized *P/C* ratios of AzoTag16-miRFP680-cRaf are plotted across time to evaluate cRaf translocation (bottom). The normalized *N/C* ratios of mScarlet-ERK are plotted as a function of time to assess ERK activity (top). Data are presented as the mean ± SD (n = 8 cells from three independent experiments).

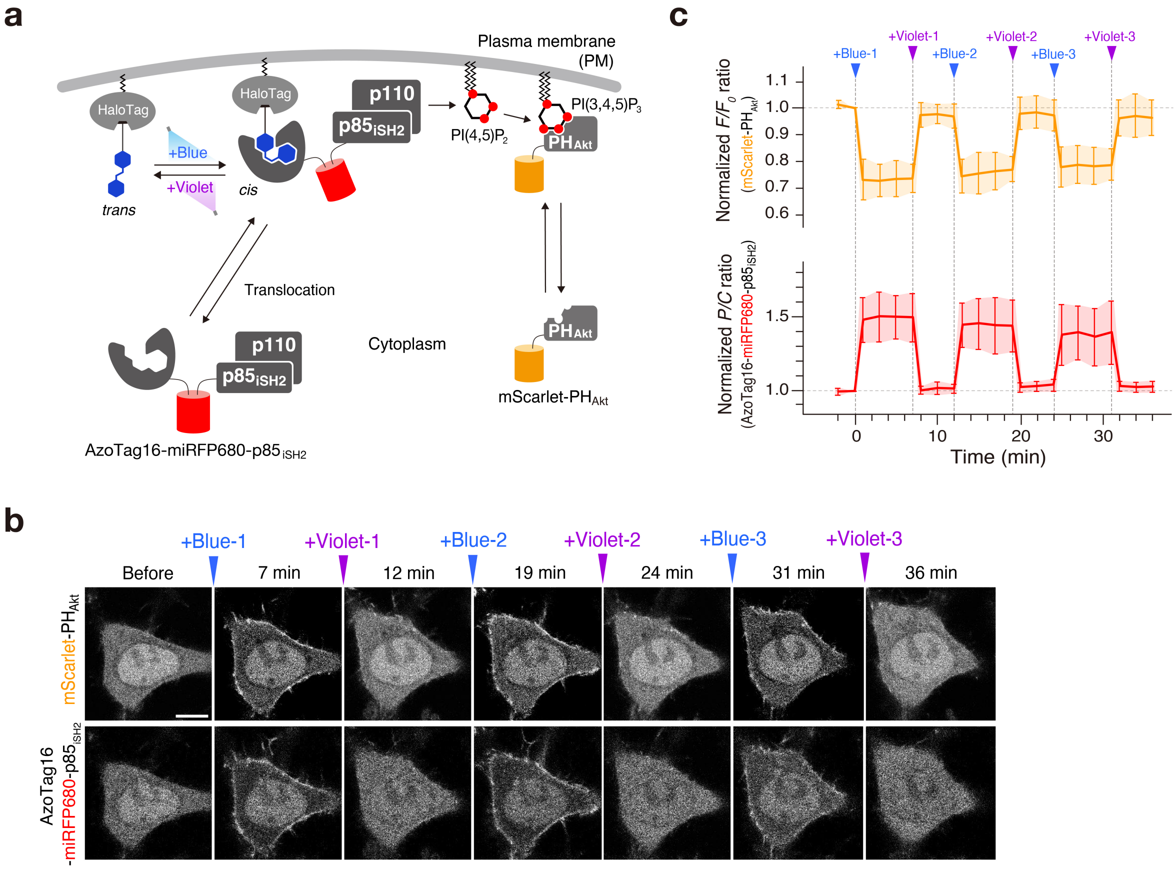


**Supplementary Figure 16. Reversible dual-light control of PI3K-mediated PI(3,4,5)$P_3$ production.**
(**a**) Schematic illustration of the experimental setup using AzoTag16-fused PI3K. (**b**) Representative time-lapse confocal fluorescence images of a HeLa cell coexpressing AzoTag16-miRFP680-$p85_{iSH2}$, Ca-FAzo-$HaloTag^{PM}$, and mScarlet-$PH_{AKT}$ before and after sequential irradiation with blue light (483 nm, "+Blue") and violet light (400 nm, "+Violet"). Images were acquired at the indicated time points. Scale bar, 10 µm. For time-lapse movies, see **Supplementary Video 5**. (**c**) Time course of PI3K translocation and PI(3,4,5)$P_3$ production. The normalized *P/C* ratios of AzoTag16-miRFP680-$p85_{iSH2}$ are plotted across time to evaluate PI3K translocation (bottom). The normalized *P/C* ratios of mScarlet-$PH_{AKT}$ are plotted as a function of time to assess PM PI(3,4,5)$P_3$ levels (top). Arrowheads indicate irradiation with 483 nm (blue) and 400 nm (violet) light. Data are presented as the mean ± SD (n = 10 cells from three independent experiments).

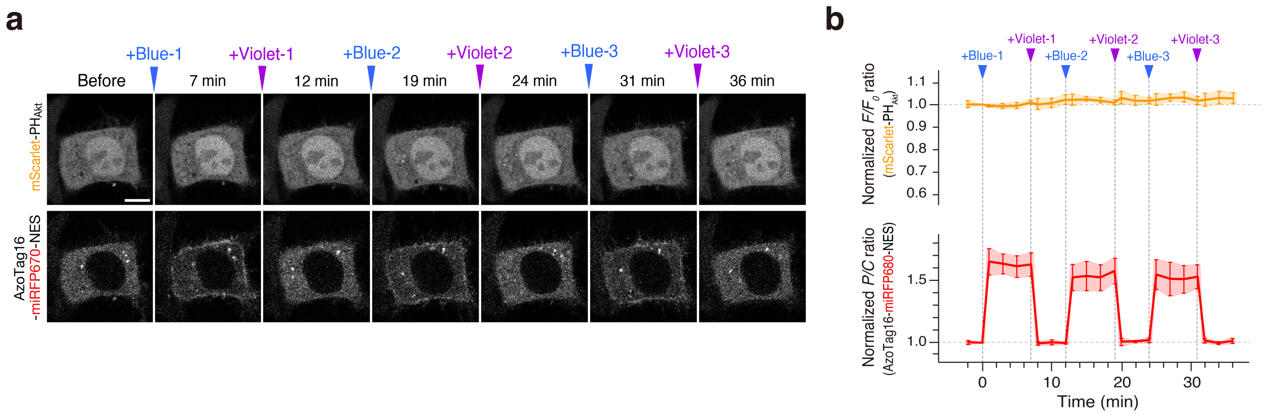


**Supplementary Figure 17. Light control of AzoTag16-miRFP670-NES lacking the iSH2 domain.**
(**a**) Representative time-lapse confocal fluorescence images of a HeLa cell coexpressing AzoTag16-miRFP670-NES lacking the iSH2 domain, Ca-FAzo-HaloTag$^{PM}$, and mScarlet-PH$_{AKT}$ before and after sequential irradiation with blue light (483 nm, "+Blue") and violet light (400 nm, "+Violet"). Images were acquired at the indicated time points. Scale bar, 10 µm. (**b**) Time course of PI3K translocation and PI(3,4,5)$P_3$ production. The normalized *P/C* ratios of AzoTag16-miRFP670-NES are plotted across time to evaluate PI3K translocation (bottom). The normalized *P/C* ratios of mScarlet-PH$_{AKT}$ are plotted as a function of time to assess PM PI(3,4,5)$P_3$ levels (top). Arrowheads indicate irradiation with 483 nm (blue) and 400 nm (violet) light. Data are presented as the mean ± SD (n = 8 cells from three independent experiments).

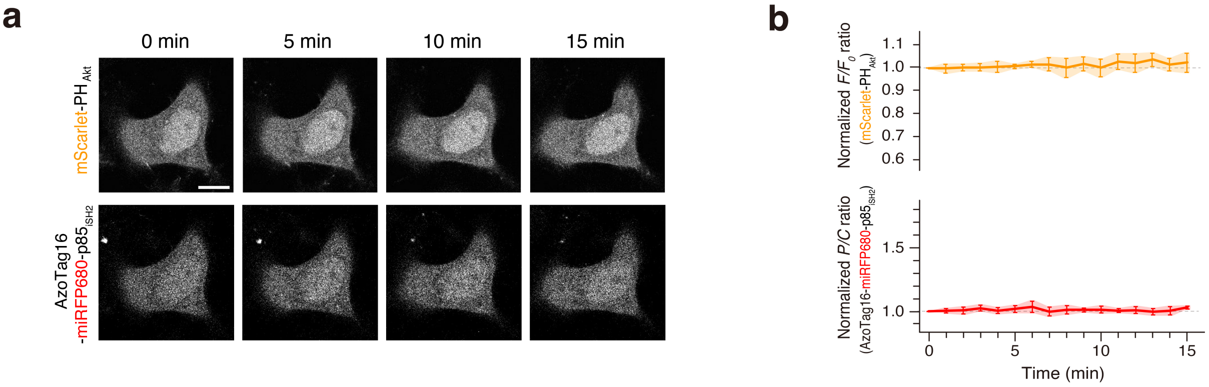


**Supplementary Figure 18. Localization and activity changes of AzoTag16-miRFP680-p85$_{iSH2}$ in the dark.** (**a**) Representative time-lapse confocal fluorescence images of a HeLa cell coexpressing AzoTag16-miRFP680-p85$_{iSH2}$, Ca-FAzo-HaloTag$^{PM}$, and mScarlet-PH$_{AKT}$ in the absence of irradiation. Images were acquired at the indicated time points. Scale bar, 10 µm. (**b**) Time course of PI3K translocation and PI(3,4,5)$P_3$ production. The normalized *P/C* ratios of AzoTag16-miRFP670-NES are plotted across time to evaluate PI3K translocation (bottom). The normalized *P/C* ratios of mScarlet-PH$_{AKT}$ are plotted across time to assess PM PI(3,4,5)$P_3$ levels (top). Data are presented as the mean ± SD (n = 8 cells from three independent experiments).

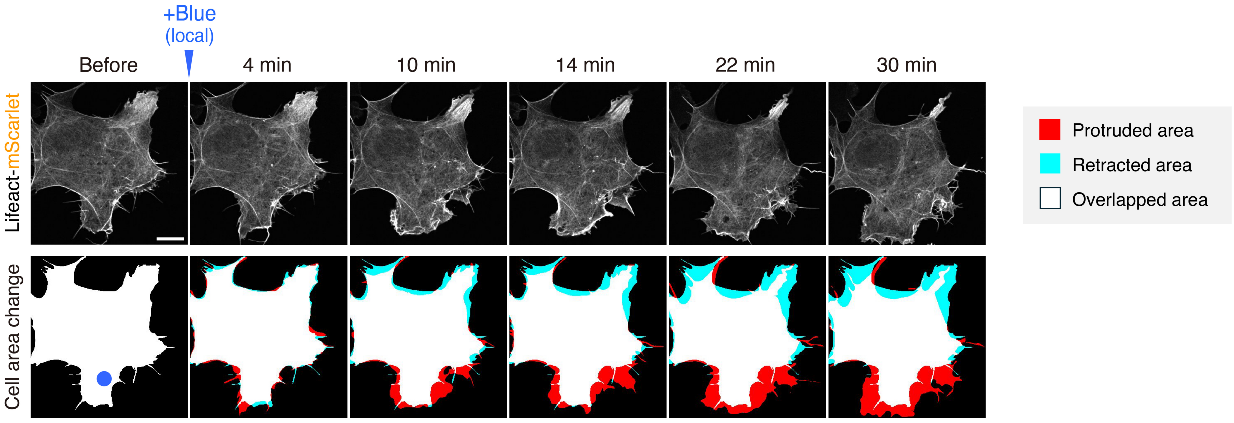


**Supplementary Figure 19. Sustained lamellipodium formation induced by single local light irradiation.** Representative time-lapse confocal fluorescence images of a Cos-7 cell coexpressing AzoTag16-miRFP680-$p85_{iSH2}$, Ca-FAzo-HaloTag$^{PM}$, and Lifeact-mScarlet before and after a single local light irradiation with a 488 nm laser. The blue circle indicates the region of local 488 nm laser irradiation. Scale bar, 20 µm. For time-lapse movies, see **Supplementary Video 6**.

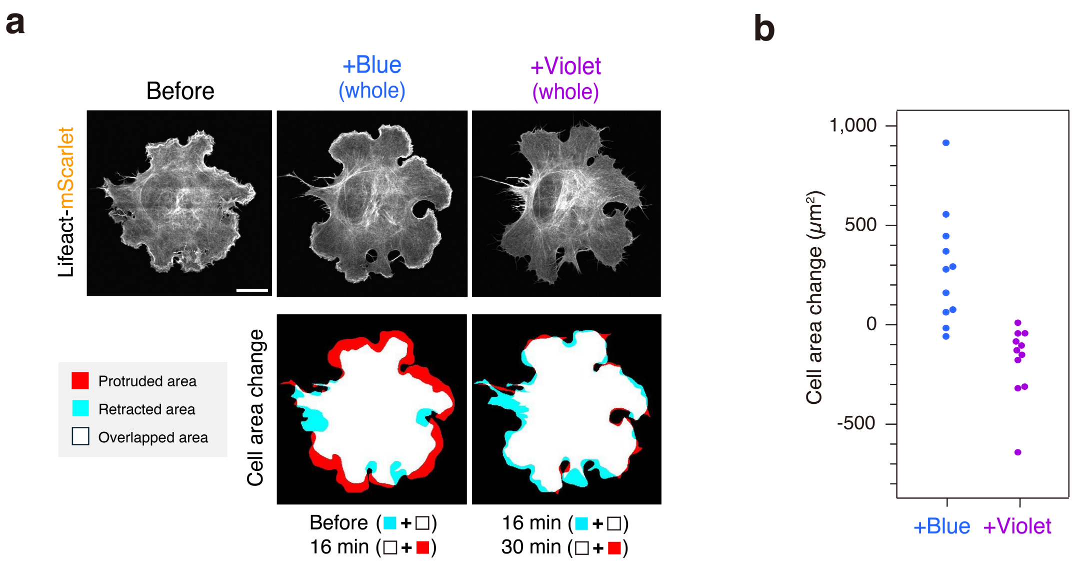


**Supplementary Figure 20. Global activation of AzoTag16-fused PI3K by whole-cell light irradiation.** (**a**) Representative time-lapse confocal fluorescence images of a Cos-7 cell coexpressing AzoTag16-miRFP680-p85$_{iSH2}$, Ca-FAzo-HaloTag$^{PM}$, and Lifeact-mScarlet before and after sequential whole-cell irradiation with a 488 nm laser ("+Blue") and a 405 nm laser ("+Violet"). The +Blue image was acquired 16 min after 488 nm laser irradiation, while the +Violet image was acquired 14 min after 405 nm laser irradiation. (**b**) Quantification of cell area changes. Cell area changes (μm²) were calculated by determining the difference in cell area between pre- and post-488 nm irradiation (blue) and between post-488 nm and post-405 nm irradiation (pink). Each dot represents a single-cell measurement (n = 11 cells).

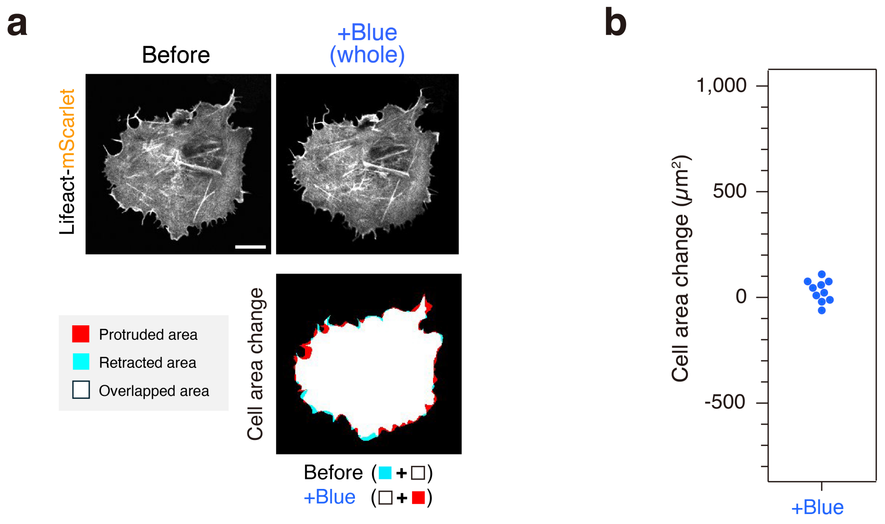


**Supplementary Figure 21. Light-mediated lamellipodia induction by AzoTag16-miRFP670-NES lacking the iSH2 domain.**

(**a**) Representative confocal fluorescence images of a Cos-7 cell coexpressing AzoTag16-miRFP670-NES lacking the iSH2 domain, Ca-FAzo-HaloTag$^{PM}$, and Lifeact-mScarlet before and 14 min after whole-cell irradiation with a 488 nm laser ("+Blue"). (**b**) Quantification of cell area changes. Cell area changes (μm²) were calculated by determining the difference in cell area between pre- and post-488 nm irradiation. Each dot represents a single-cell measurement (n = 10 cells).

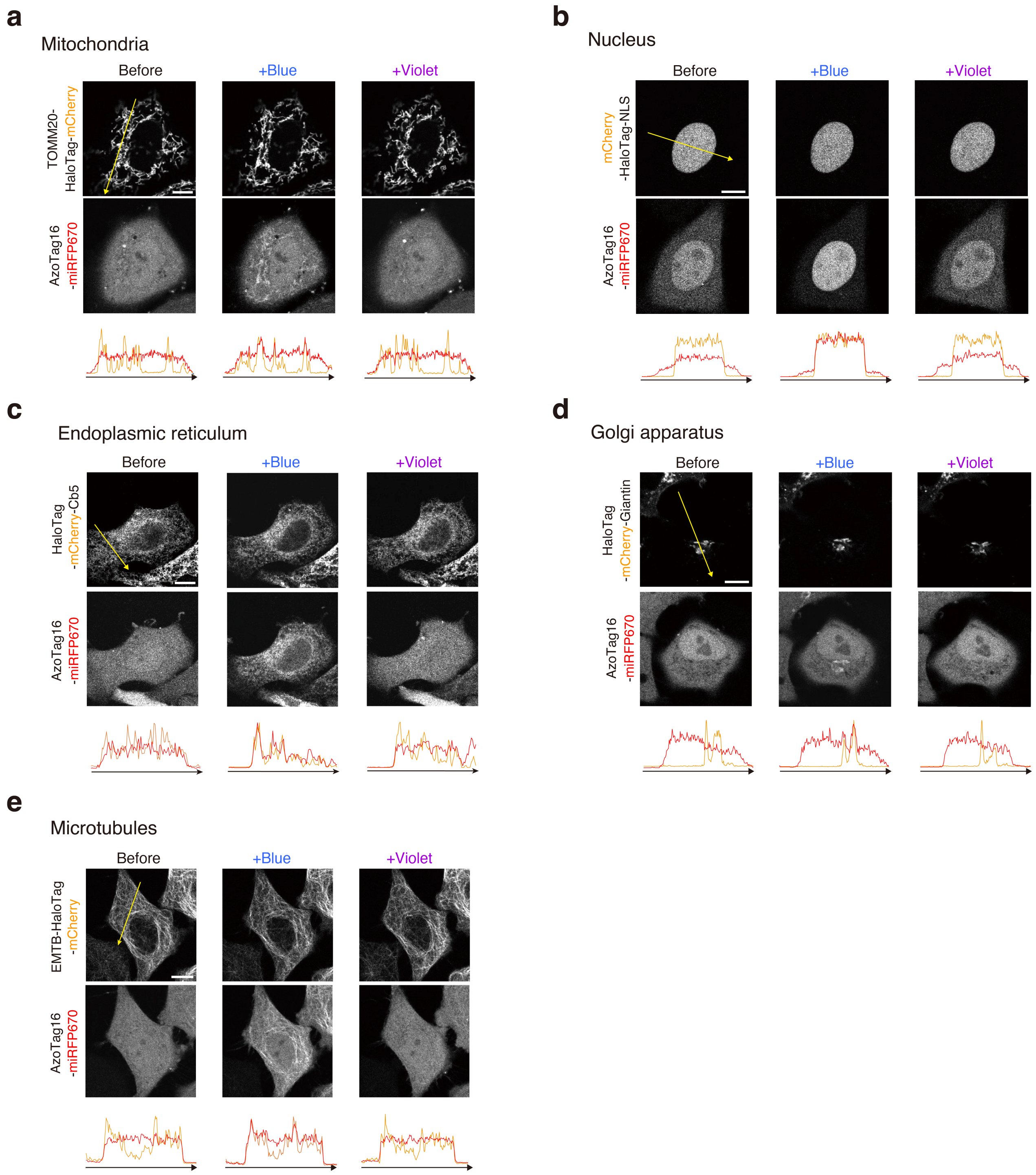


**Supplementary Figure 22. Application of the light-switchable AzoTag16/HaloTag dimerization system to various organelles.**
Representative confocal fluorescence images of HeLa cells coexpressing AzoTag16-miRFP670 with various Ca-FAzo–conjugated HaloTag fusions: TOMM20-HaloTag-mCherry for mitochondria (**a**), mCherry-HaloTag-NLS for the nucleus (**b**), HaloTag-mCherry-Cb5 for the endoplasmic reticulum (**c**), HaloTag-mCherry-Giantin for the Golgi apparatus (**d**), and EMTB-HaloTag-mCherry for microtubules (**e**). Images were acquired before and 2 min after irradiation with blue light (483 nm, "+Blue") and violet light (400 nm, "+Violet"). Bottom panels

show line scan intensity profiles of the organelle-targeted HaloTag (orange) and AzoTag16-miRFP670 (red) along the yellow dashed lines. Scale bars, 10 μm.

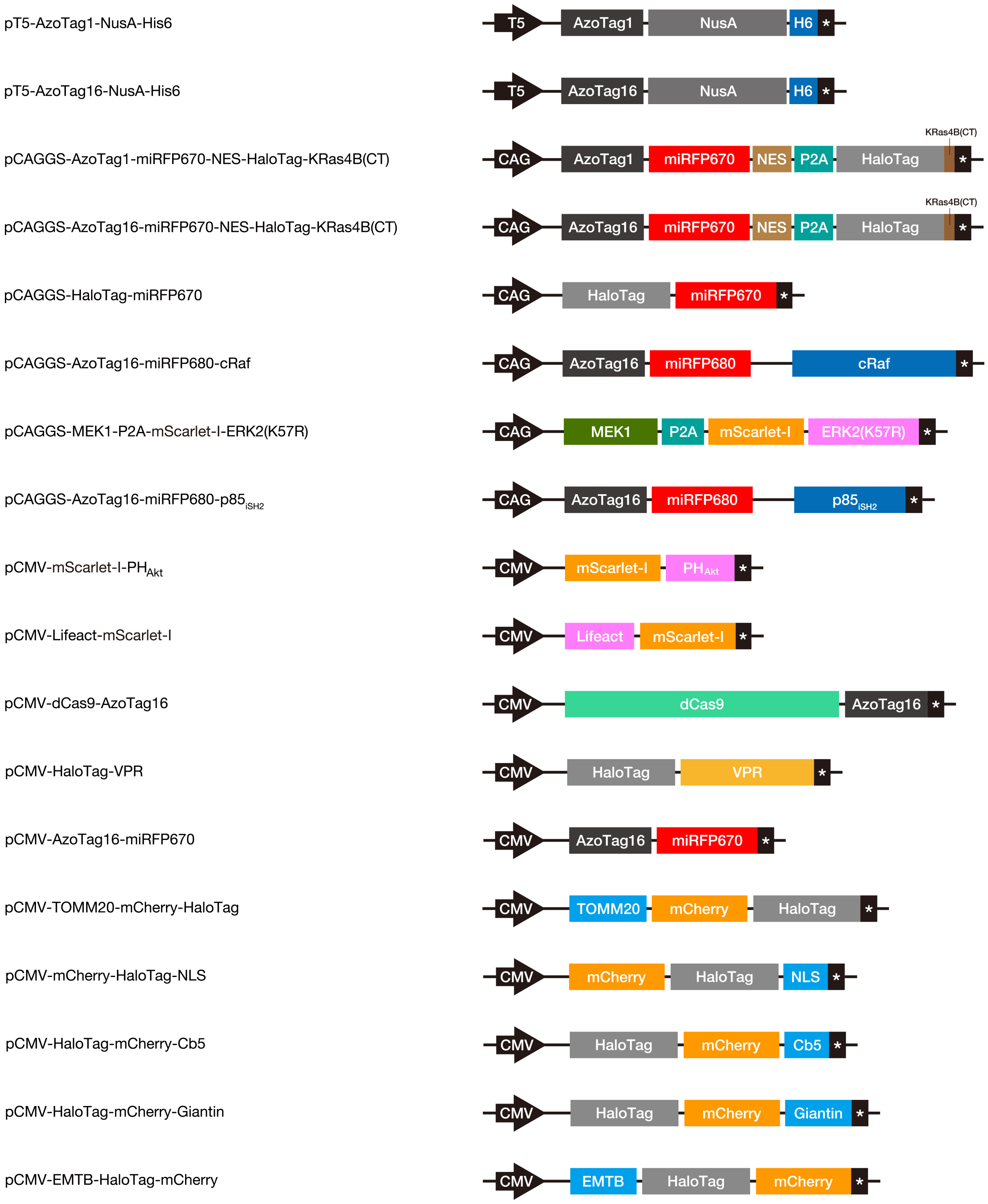


**Supplementary Figure 23. Schematic illustration of the domain structures of fusion proteins constructed in this study.**

DNA and amino acid sequences of the constructs are provided in the **Supplementary Sequences** section.

### Supplementary Tables

**Supplementary Table 1. Photochemical and photophysical properties of FAzo ligands.**

| Compound | π→π* $\lambda_{max}$ (*trans*) [nm] | ε (*trans*) [$M^{-1}$ $cm^{-1}$] | n→π* $\lambda_{max}$ (*trans*) [nm] | ε (*trans*) [$M^{-1}$ $cm^{-1}$] | n→π* $\lambda_{max}$ (*cis*) [nm] | ε (*cis*) [$M^{-1}$ $cm^{-1}$] | $PSS_{cis}$ [%] 360 | 380 | 400 | 420 | 438 | 488 | 516 |
|---|---|---|---|---|---|---|---|---|---|---|---|---|---|
| Et-FAzo (**1**) | 317 | 21,900 | 450 | 1,670 | 413 | 2,000 | 57 | 3 | 5 | 10 | 14 | 62 | 79 |
| Ca-FAzo (**2**) | 316 | 22,500 | 448 | 1,810 | 415 | 2,300 | 67 | 14 | 7 | 15 | 30 | 65 | 81 |

**Supplementary Table 2. Summary of next-generation sequencing analysis of anticalin clones collected in the 8th round of selection.**

| Targets | Frequency (%) | *cis/trans* ratio | Amino acids at the randomized positions 40 | 43 | 47 | 48 | 56 | 59 | 61 | 75 | 77 | 84 | 86 | 88 | 107 | 110 | 113 | 130 | 132 | 134 | 139 | 141 | 143 | Note |
|---|---|---|---|---|---|---|---|---|---|---|---|---|---|---|---|---|---|---|---|---|---|---|---|---|
| *cis*-Et-FAzo | 51.5 | 1.7 | Q | R | F | N | T | Q | I | P | Y | Q | N | D | Y | N | E | V | W | Y | W | S | Y | AzoTag1 |
| | 32.1 | 1.1 | Y | Y | Y | P | E | Y | D | T | Y | W | M | A | Y | W | I | G | Q | Y | Y | S | W | |
| | 5.6 | 1.1 | Y | Y | Y | P | E | Y | D | T | Y | W | M | A | Y | L | I | G | Q | Y | Y | S | W | |
| | 2.1 | 0.4 | T | W | I | Y | Y | A | R | W | W | F | E | Y | N | I | P | S | L | W | Y | Y | V | |
| | 2.0 | 0.3 | I | Y | D | F | F | W | R | G | F | Y | H | H | Y | Y | Q | T | Y | Y | M | Y | V | |
| *cis*-Ca-FAzo | 27.6 | 3.8 | I | Y | D | F | F | W | R | G | F | Y | H | H | Y | Y | Q | T | Y | Y | M | Y | V | AzoTag16 |
| | 23.1 | 3.8 | V | F | F | Y | A | W | K | Y | D | Y | N | F | Q | Y | T | N | F | Y | Y | Y | V | |
| | 15.6 | 1.6 | Y | H | Y | R | Y | Q | R | W | L | S | Y | Y | M | K | A | S | Y | H | I | Y | S | |
| | 10.3 | 1.9 | T | W | I | Y | Y | A | R | W | W | F | E | Y | N | I | P | S | L | W | Y | Y | V | |
| | 5.6 | 0.2 | Q | R | F | N | T | Q | I | P | Y | Q | N | D | Y | N | E | V | W | Y | W | S | Y | |

**Supplementary Table 3. Thermodynamic parameters of Et-FAzo/AzoTag1 and Ca-FAzo/AzoTag16 pairs.**

| Clones | Targets | $K_d$ (nM) | $\Delta H$ (kcal $mol^{-1}$) | $-T\Delta S$ (kcal $mol^{-1}$) | $\Delta G$ (kcal $mol^{-1}$) | $N$ |
|---|---|---|---|---|---|---|
| AzoTag1 | *cis*-Et-FAzo | 804.7 ± 22.6 | –4.85 ± 0.52 | –3.47 ± 0.53 | –8.31 ± 0.02 | 0.98 ± 0.03 |
| | *trans*-Et-FAzo | n.d. | | | | |
| AzoTag16 | *cis*-Ca-FAzo | 14.03 ± 0.02 | –19.31 ± 0.43 | 8.59 ± 0.44 | –10.71 ± 0.01 | 0.92 ± 0.09 |
| | *trans*-Ca-FAzo | n.d. | | | | |

**Supplementary Table 4. Oligonucleotides used for anticalin library preparation.**
**→ Excel file**

**Supplementary Table 5. Composition of the TRAP reaction solution.**
**→ Excel file**

**Supplementary Table 6. Next-generation sequencing results for anticalin clones collected in the eighth round of selection.**
**→ Excel file**

#### Supplementary Videos

**Supplementary Video 1. Time-lapse movie of Figure 3g. Scale bars, 10 µm.**

**Supplementary Video 2. Time-lapse movie of Figure 3h. Scale bars, 10 µm.**

**Supplementary Video 3. Time-lapse movie of Supplementary Figure 13.**

**Supplementary Video 4. Time-lapse movie of Figure 4b,c.**

**Supplementary Video 5. Time-lapse movie of Supplementary Figure 16b,c.**

**Supplementary Video 6. Time-lapse movie of Supplementary Figure 19.**

**Supplementary Video 7. Time-lapse movie of Figure 5b.**

#### Supplementary Methods: Chemical Synthesis

##### General materials and methods

All chemical reagents and solvents were purchased from commercial suppliers (Watanabe Chemical Industries, Tokyo Chemical Industry, FUJIFILM Wako Pure Chemical Corp., and Kanto Chemical) and used without further purification. Compounds **11**[S2] and **19**[S3] were synthesized as previously reported.

Thin-layer chromatography (TLC) was performed on silica gel 60 $F_{254}$ precoated aluminum sheets (Merck), and TLC plates were visualized by fluorescence quenching. Flash column chromatography was performed using silica gel 60 N (neutral, 40–50 μm) (Kanto Chemical). Reversed-phase HPLC was performed on a Hitachi LaChrom Elite system with UV detection at 220 nm using a YMC-Pack ODS-A column (10 × 250 mm or 20 × 250 mm) or YMC-Pack C4 column (10 × 250 mm or 20 × 250 mm).

$^{1}$H NMR spectra were recorded on a Bruker AVANCE III HD400SJ (400 MHz) spectrometer. $^{1}$H NMR chemical shifts were referenced to tetramethylsilane (0 ppm). MALDI-TOF mass spectra were acquired on a JEOL JMS-S3000 Spiral TOF mass spectrometer and a Bruker autoflex maX using α-cyano-4-hydroxycinnamic acid (CHCA) as a matrix. High-resolution mass spectra were measured on a Waters SYNAPT G2-Si mass spectrometer.

##### Reagent abbreviations

Boc-Adox-OH Boc-8-amino-3,6-dioxaoctanoic acid; DIPEA, *N*,*N*-diisopropylethylamine; DMF, *N*,*N*-dimethylformamide; EDC·HCl, 1-ethyl-3-(3-dimethylaminopropyl)carbodiimide hydrochloride; Fmoc-Adox-OH, Fmoc-8-amino-3,6-dioxaoctanoic acid; HBTU, 1-[bis(dimethylamino)methylene]-1*H*-benzotriazolium 3-oxide hexafluorophosphate; HOBt, 1-hydroxybenzotriazole; TFA, trifluoroacetic acid; THF, tetrahydrofuran; TIPS, triisopropylsilane.

#### Synthesis of Et-FAzo (1)

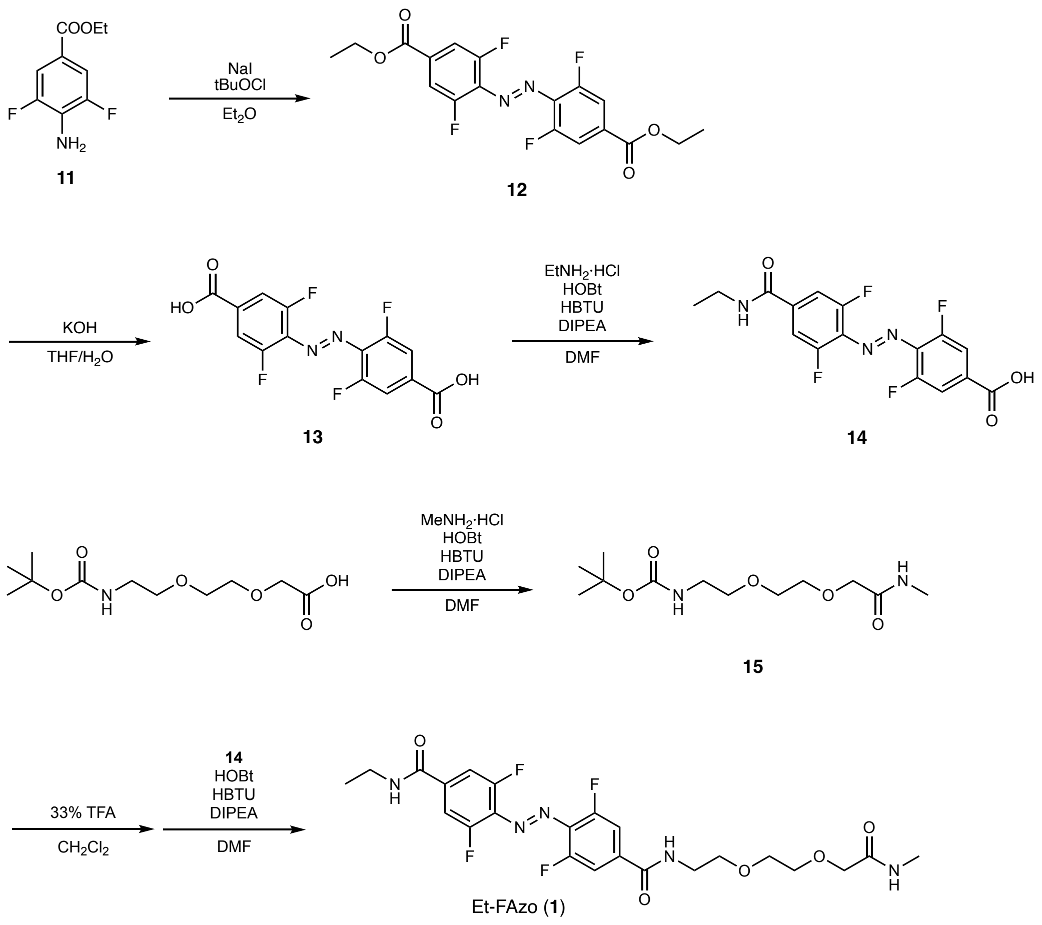


**Scheme S1.** Synthetic route of Et-FAzo (**1**)

Compound **12**

Compound **12** was synthesized using a modified procedure based on a previously reported method.[S4] Compound **11**[S2] (101 mg, 0.50 mmol) and sodium iodide (149 mg, 1.0 mmol) were dissolved in anhydrous $Et_2O$ (4.0 mL) and stirred under an argon atmosphere. To this solution, *tert*-butyl hypochlorite (113 μL, 1.0 mmol) was added, and the resulting mixture was stirred at room temperature for 24 h. The reaction mixture was then diluted with water (50 mL) and extracted twice with $CH_2Cl_2$ (50 mL each). The combined organic layers were dried over anhydrous $MgSO_4$, filtered, and concentrated under reduced pressure. The crude residue was purified by column chromatography (silica gel, hexane/EtOAc = 4:1), yielding compound **12** (64.5 mg, 65%) as a red solid.

$^1$H NMR (400 MHz, $CDCl_3$): δ 7.75 (4H, dd, $J$ = 1.48, 11.90 Hz), 4.44 (4H, q, $J$ = 7.13 Hz), 1.43 (6H, t, $J$ = 7.16 Hz).

Compound **13**

Compound **12** (51 mg, 128 μmol) was dissolved in THF (1.7 mL) and stirred. To this solution, 0.5 M KOH aqueous solution (0.8 mL) was added, and the resulting mixture was refluxed for 2 h. After cooling to room temperature, the reaction mixture was diluted with water (15 mL), and the aqueous layer was washed twice with EtOAc (20 mL each). The aqueous layer was then acidified to pH 2 with 1 M HCl solution, resulting in the formation of a precipitate. The precipitate was filtered, washed with water, and dried *in vacuo*, yielding compound **13** (33.1 mg, 76%) as a red solid.

$^1$H NMR (400 MHz, DMSO-$d_6$): δ 13.92 (2H, br s), 7.82 (4H, d, $J$ = 9.44 Hz).

Compound **14**

Compound **13** (30 mg, 87 μmol) was dissolved in anhydrous DMF (1.0 mL) and stirred under an argon atmosphere. To this solution, DIPEA (75.7 μL, 435 μmol), HOBt (13.2 mg, 96 μmol), HBTU (36.4 mg, 96 μmol), and ethylamine hydrochloride (7.8 mg, 96 μmol) were sequentially added. The resulting mixture was stirred at room temperature for 3 h and then concentrated under reduced pressure. The crude residue was purified by column chromatography (silica gel, $CHCl_3$/MeOH/AcOH = 100:10:1), yielding compound **14** (13.3 mg, 41%) as a red solid.

$^1$H NMR (400 MHz, $CD_3OD$): δ 7.75 (2H, d, $J$ = 9.20 Hz), 7.66 (2H, d, $J$ = 9.52 Hz), 3.43 (2H, q, $J$ = 7.25 Hz), 1.25 (3H, t, $J$ = 7.28 Hz).

Compound **15**

Boc-Adox-OH (150 mg, 337 μmol) was dissolved in anhydrous DMF (3.0 mL) and stirred under an argon atmosphere. To this solution, DIPEA (290 μL, 1.69 mmol), HOBt (51 mg, 370 μmol), HBTU (140 mg, 370 μmol), and methylamine hydrochloride (32 mg, 370 μmol) were sequentially added. The resulting mixture was stirred at room temperature for 5 h. The reaction mixture was then diluted with water (50 mL) and extracted with EtOAc (50 mL). The organic layer was washed twice with saturated $NaHCO_3$ aqueous solution (30 mL each), twice with 5% citric acid aqueous solution (30 mL each), and finally once with brine (30 mL). The organic layer was dried over anhydrous $Na_2SO_4$, filtered, and concentrated under reduced pressure. The crude residue was purified by column chromatography (silica gel, $CHCl_3$/MeOH = 10:1), yielding compound **15** (36.9 mg, 44%) as a colorless and transparent oil.

$^1$H NMR (400 MHz, $CDCl_3$): δ 4.00 (2H, s), 3.68–3.62 (4H, m), 3.56 (2H, t, $J$ = 5.32 Hz), 3.34 (2H, q, $J$ = 5.01 Hz), 2.86 (3H, d, $J$ = 8.36 Hz), 1.45 (9H, s).

Compound **1** (Et-FAzo)

Compound **15** (50 mg, 181 μmol) was dissolved in a mixture of $CH_2Cl_2$ (4.0 mL) and TFA (2.0 mL) and stirred at room temperature for 1 h. The reaction mixture was concentrated under reduced pressure, and the residue was dissolved in toluene (2.0 mL) before being concentrated again. This process was repeated to remove TFA, after which the residue was dissolved in anhydrous DMF (1.0 mL).

In a separate flask, compound **14** (55 mg, 150 μmol) was dissolved in anhydrous DMF (4.0 mL) and stirred under an argon atmosphere. To this solution, DIPEA (102 μL, 750 μmol), HOBt (24.9 mg, 181 μmol), HBTU (68.7 mg, 181 μmol), and the DMF solution of the deprotected compound **15** prepared above were sequentially added. The resulting mixture was stirred at room temperature for 3 h and then concentrated under reduced pressure. The crude residue was purified by reversed-phase HPLC using a semi-preparative ODS column with a linear gradient of MeCN containing 0.1% TFA and 0.1% aqueous TFA, yielding Et-FAzo (**1**) (13.9 mg, 21%) as a red solid consisting of a 3:1 mixture of *trans*- and *cis*-forms.

**Purification of *trans*- and *cis*-Et-FAzo**

Et-FAzo (**1**) (13.9 mg) was dissolved in a mixture of MeCN (2 mL) and $H_2O$ (2 mL) and irradiated with 400 nm light for 5 min. The *trans*- and *cis*-forms of Et-FAzo were then separated and purified by reversed-phase HPLC using a semi-preparative C4 column with a linear gradient from 40% to 45% MeCN/$H_2O$ containing 0.1% TFA over 30 min (**Supplementary Figure 1a**), yielding *trans*-Et-FAzo (6.2 mg, 45%) and *cis*-Et-FAzo (5.8 mg, 42%) as red solids.

*trans*-Et-FAzo

$^1H$ NMR (400 MHz, $CD_3OD$): δ 7.69–7.65 (4H, m), 3.98 (2H, s), 3.70 (6H, t, $J$ = 5.24 Hz), 3.62 (2H, q, $J$ = 4.77 Hz), 3.43 (2H, q, $J$ = 7.25 Hz), 2.76 (3H, s), 1.24 (3H, t, $J$ = 7.26 Hz).

HRMS (ESI): calculated for $[M+Na]^+$, 550.1690; found, 550.1694.

*cis*-Et-FAzo

$^1H$ NMR (400 MHz, $CD_3OD$): δ 7.50–7.45 (4H, m), 3.95 (2H, s), 3.66–3.62 (6H, m), 3.55–3.52 (2H, m), 3.35 (2H, q, $J$ = 7.29 Hz), 2.72 (3H, s), 1.18 (3H, t, $J$ = 7.20 Hz).

HRMS (ESI): calculated for $[M+Na]^+$, 550.1690; found, 550.1693.

**Synthesis of Ca-FAzo (2)**

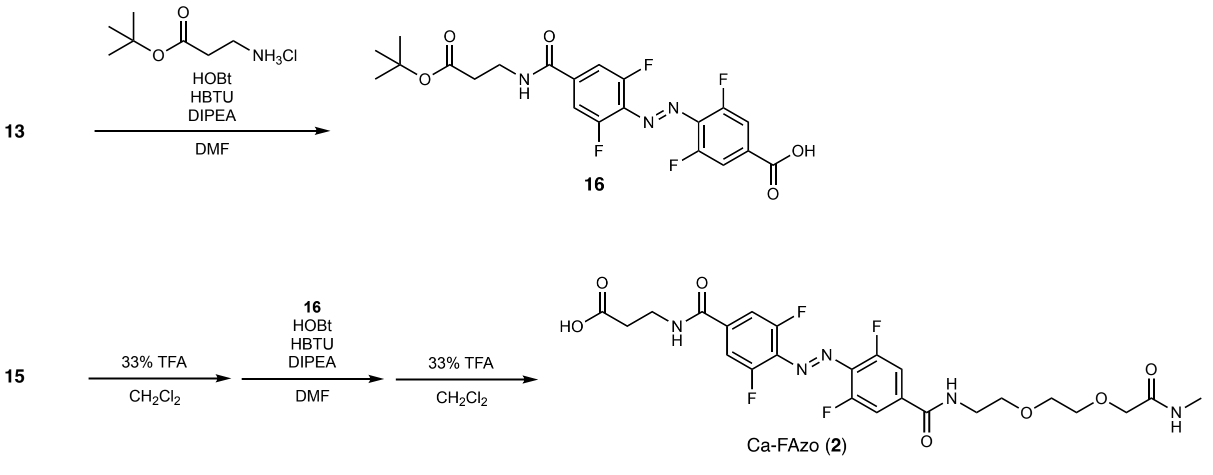


**Scheme S2.** Synthetic route of Ca-FAzo (**2**)

Compound **16**

Compound **13** (36 mg, 92.8 μmol) was dissolved in anhydrous DMF (1.0 mL) and stirred under an argon atmosphere. To this solution, DIPEA (78.7 μL, 464 μmol), HOBt (14.3 mg, 104 μmol), HBTU (39.5 mg, 104 μmol), and *β*-alanine *tert*-butyl ester hydrochloride (25 mg, 104 μmol) were sequentially added. The resulting mixture was stirred at room temperature for 3 h and then concentrated under reduced pressure. The crude residue was purified by column chromatography (silica gel, $CHCl_3$/MeOH/AcOH = 90:9:1), yielding compound **16** (24.5 mg, 56%) as a red solid.

$^1H$ NMR (400 MHz, $CD_3OD$): δ 7.75–7.63 (4H, m), 3.62 (2H, t, $J$ = 6.84), 2.59 (2H, t, $J$ = 6.90), 1.46 (9H, s).

Compound **2** (Ca-FAzo)

Compound **15** (30 mg, 102 μmol) was dissolved in a mixture of $CH_2Cl_2$ (4.0 mL) and TFA (2.0 mL) and stirred at room temperature for 1 h. The reaction mixture was concentrated under reduced pressure, and the residue was dissolved in toluene (2.0 mL) before being concentrated again. This process was repeated to remove TFA, after which the residue was dissolved in anhydrous DMF (1.0 mL).

In a separate flask, compound **16** (40 mg, 85 µmol) was dissolved in anhydrous DMF (4.0 mL) and stirred under an argon atmosphere. To this solution, DIPEA (58 µL, 425 µmol), HOBt (23.5 mg, 170 µmol), HBTU (64.6 mg, 170 µmol), and the DMF solution of the deprotected compound **15** prepared above were sequentially added. The resulting mixture was stirred at room temperature for 3 h and then concentrated under reduced pressure. The residue was dissolved in $CH_2Cl_2$ (4.0 mL) and TFA (2.0 mL) and stirred at room temperature for 2 h, followed by concentration under reduced pressure. The residue was dissolved in toluene (2.0 mL) and concentrated again under reduced pressure. This process was repeated to remove the TFA. The crude residue was purified by reversed-phase HPLC using a semi-preparative ODS column with a linear gradient of MeCN containing 0.1% TFA and 0.1% aqueous TFA, yielding Ca-FAzo (**2**) (24.2 mg, 50%) as a red solid (a 10:1 mixture of *trans*- and *cis*-forms).

**Purification of *trans*- and *cis*-Ca-FAzo**

Ca-FAzo (**2**) (24.2 mg) was dissolved in a mixture of MeCN (2 mL) and $H_2O$ (2 mL). The solution was irradiated with 400 nm light for 5 min. The *trans*- and *cis*-forms of Ca-FAzo were then separated and purified by reversed-phase HPLC using a semi-preparative ODS column with a linear gradient from 40% to 42% MeCN/$H_2O$ containing 0.1% TFA over 30 min (**Supplementary Figure 1b**), yielding *trans*-Ca-FAzo (10.8 mg, 45%) and *cis*-Ca-FAzo (11.4 mg, 47%) as red solids.

*trans*-Ca-FAzo

$^1$H NMR (400 MHz, $CD_3OD$): δ 7.71–7.64 (4H, m), 3.98 (2H, s), 3.70 (6H, t, *J* = 4.88 Hz), 3.66–3.60 (4H, m), 2.76 (3H, s), 2.64 (2H, t, *J* = 6.84 Hz).

HRMS (ESI): calculated for $[M+Na]^+$, 594.1588; found, 594.1577.

*cis*-Ca-FAzo

$^1$H NMR (400 MHz, $CD_3OD$): δ 7.49–7.45 (4H, m), 3.94 (2H, s), 3.65–3.62 (6H, m), 3.59–3.52 (4H, m), 2.72 (3H, s), 2.59 (2H, t, *J* = 6.86 Hz).

HRMS (ESI): calculated for $[M+Na]^+$, 594.1588; found, 594.1573.

#### Synthesis of compound 17

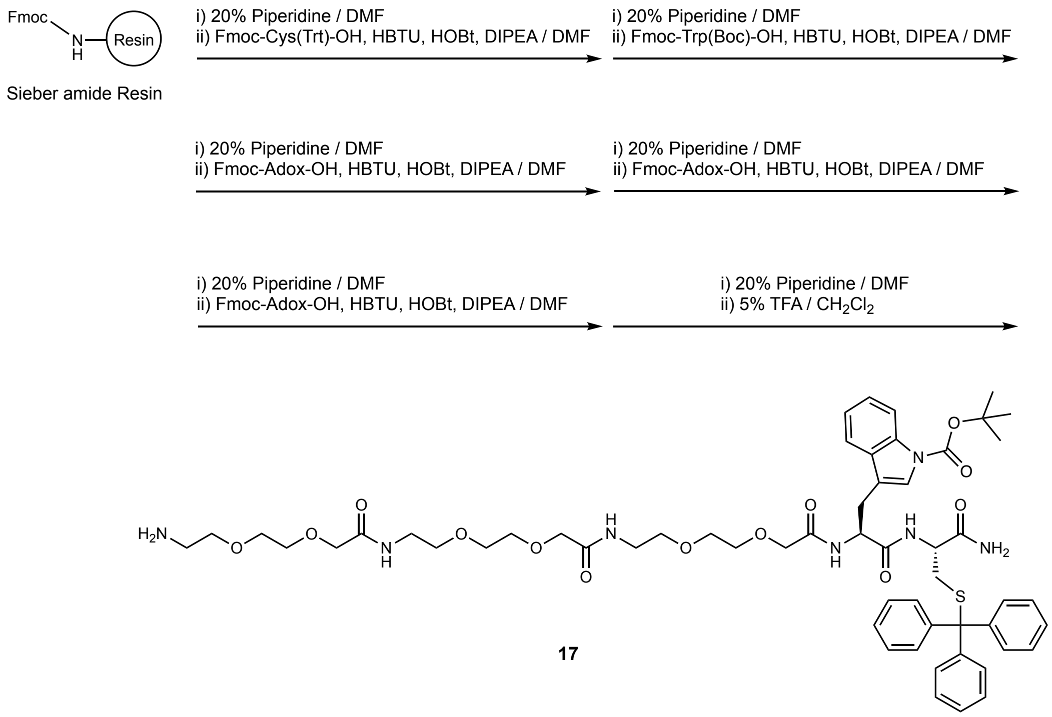


**Scheme S3.** Synthetic route of compound **17**

#### General methods for solid-phase synthesis

Solid-phase synthesis was performed manually using Sieber Amide resin according to standard Fmoc-based solid-phase peptide synthesis protocols. Fmoc deprotection was carried out with 20% piperidine in DMF at room temperature for 15 min. Amino acid coupling reactions were performed at room temperature with a mixture of Fmoc-protected amino acid (3.1 eq.), HBTU (3.0 eq.), HOBt (3.0 eq.), and DIPEA (6.0 eq.) in DMF. All standard Fmoc deprotection and coupling reactions were monitored using the Kaiser test.[S5] All washing procedures were completed with DMF.

Compound **17**

Compound **17** was synthesized on Sieber Amide resin (0.66 mmol/g) (61 mg, 40 μmol). Fmoc-Cys(Trt)-OH, Fmoc-Trp(Boc)-OH, and Fmoc-Adox-OH (×3) were sequentially coupled to the resin. After Fmoc deprotection, the protected compound **17** was cleaved from the resin by treatment with $CH_2Cl_2$ (2 mL) containing 5% TFA, followed by collection of the filtrate. This process was performed six times. The combined filtrates were mixed with toluene (20 mL) and concentrated under reduced pressure. The crude residue was purified by reversed-phase HPLC using a semi-preparative ODS column with a linear gradient of MeCN containing 0.1% TFA and 0.1% aqueous TFA, yielding compound **17** (24.8 mg, 52%) as a white solid.

$^1$H NMR (400 MHz, $CD_3OD$): δ 8.06 (1H, d, $J$ = 8.04 Hz), 7.62 (1H, d, $J$ = 7.60 Hz), 7.52 (1H, s), 7.38–7.20 (17H, m), 4.83–4.77 (1H, m), 4.26–4.20 (1H, m), 4.00–3.89 (6H, m), 3.69–3.33 (24H, m), 3.70–3.65 (7H, m), 3.15–3.08 (2H, m), 2.58–2.54 (2H, m), 1.66 (9H, s).

**Synthesis of Et-FAzo-WC (3)**

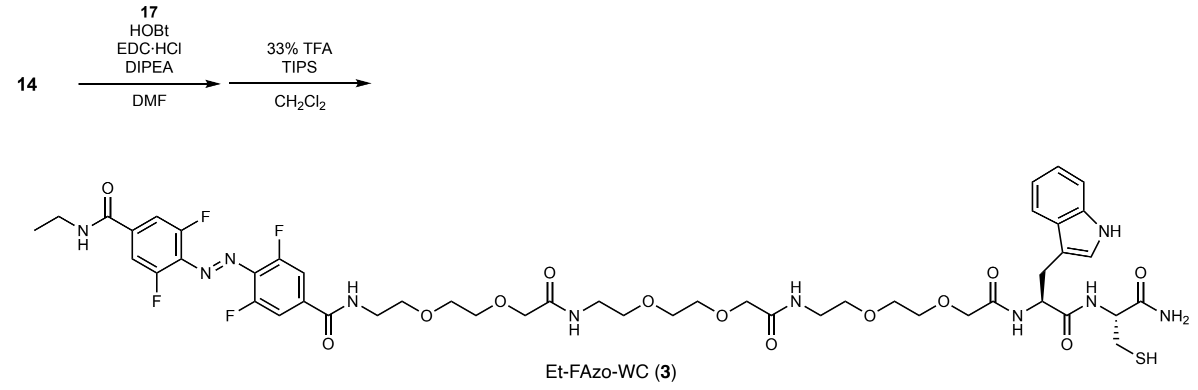


**Scheme S4.** Synthetic route of Et-FAzo-WC (**3**)

Compound **14** (12.1 mg, 10.1 μmol) was dissolved in anhydrous DMF (50 μL) and stirred under an argon atmosphere. To this solution, DIPEA (10 μL, 58 μmol), HOBt (2.5 mg, 16.3 μmol), EDC·HCl (3.1 mg, 16.3 μmol), and compound **17** (4.0 mg, 10.8 μmol) dissolved in anhydrous DMF (50 μL) were sequentially added. The resulting mixture was stirred at room temperature for 3 h and then concentrated under reduced pressure. The residue was dissolved in $CH_2Cl_2$ (1.0 mL) containing 30% TFA and 5.0% TIPS and stirred at room temperature for 2 h. The reaction mixture was subsequently concentrated under reduced pressure. The crude residue was purified by reversed-phase HPLC using a semi-preparative ODS column with a linear gradient of MeCN containing 0.1% TFA and 0.1% aqueous TFA, yielding Et-FAzo-WC (**3**) (4.2 mg, 38%) as a red solid.

$^1$H-NMR (400 MHz, $CD_3OD$): δ 7.69–7.64 (4H, m), 7.59 (1H, d, $J$ = 7.84 Hz), 7.32 (1H, d, $J$ = 8.20 Hz), 7.15 (1H, s), 7.08 (1H, t, $J$ = 7.46 Hz), 7.01 (1H, t, $J$ = 7.46 Hz), 4.74–4.70 (1H, m), 4.47–4.43 (1H, m), 4.00–3.87 (6H, m), 3.70–3.36 (26H, m), 2.84–2.72 (2H, m), 1.25 (3H, t, $J$ = 7.26 Hz).
MALDI-TOF-MS (CHCA): calculated for $[M+H]^+$, 1093.4076; found, 1093.2158.

**Synthesis of Ca-FAzo-WC (4)**

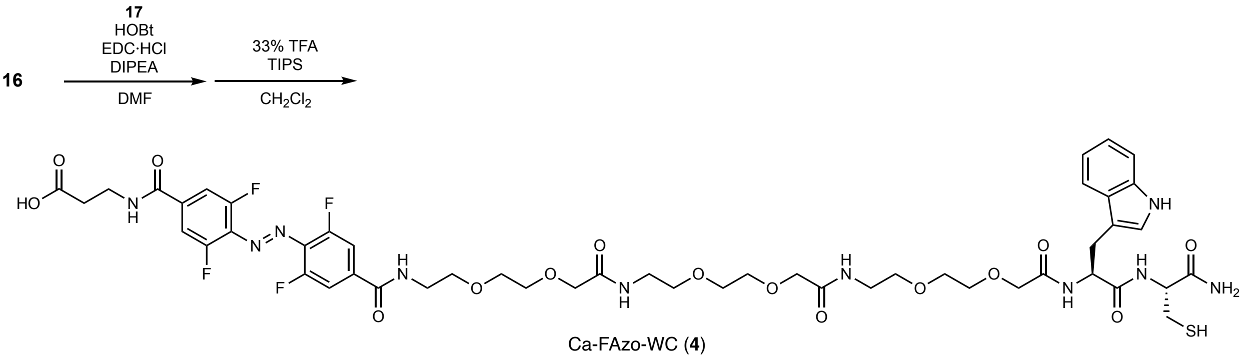


**Scheme S5.** Synthetic route of Ca-FAzo-WC (**4**)

Compound **16** (12.6 mg, 10.5 μmol) was dissolved in anhydrous DMF (50 μL) and stirred under an argon atmosphere. To this solution, DIPEA (9.1 μL, 52 μmol), HOBt (2.4 mg, 15.7 μmol), EDC·HCl (3.0 mg, 15.7 μmol), and compound **17** (6.2 mg, 13.2 μmol) dissolved in anhydrous DMF (50 μL)

were added sequentially. The resulting mixture was stirred at room temperature for 3 h and then concentrated under reduced pressure. The residue was dissolved in $CH_2Cl_2$ (1.0 mL) containing 30% TFA and 5.0% TIPS and then stirred at room temperature for 2 h. Afterward, the reaction mixture was concentrated under reduced pressure. The crude residue was purified by reversed-phase HPLC using a semi-preparative ODS column with a linear gradient of MeCN containing 0.1% TFA and 0.1% aqueous TFA, yielding Ca-FAzo-WC (**4**) (2.6 mg, 21%) as a red solid.
$^{1}$H NMR (400 MHz, $CD_3OD$): δ 7.69–7.63 (4H, m), 7.58 (1H, d, $J$ = 8.08 Hz), 7.32 (1H, d, $J$ = 7.96 Hz), 7.15 (1H, s), 7.08 (1H, t, $J$ = 7.64 Hz), 7.01 (1H, t, $J$ = 7.40 Hz), 4.75–4.70 (1H, m), 4.48–4.42 (1H, m), 4.00–3.86 (6H, m), 3.93–3.85 (1H, m), 3.70–3.40 (28H, m), 2.85–2.73 (2H, m), 2.66 (2H, t, $J$ = 7.40 Hz).
MALDI-TOF-MS (CHCA): calculated for $[M+H]^+$, 1159.3789; found, 1159.3287.

**Synthesis of Et-FAzo-biotin (5)**

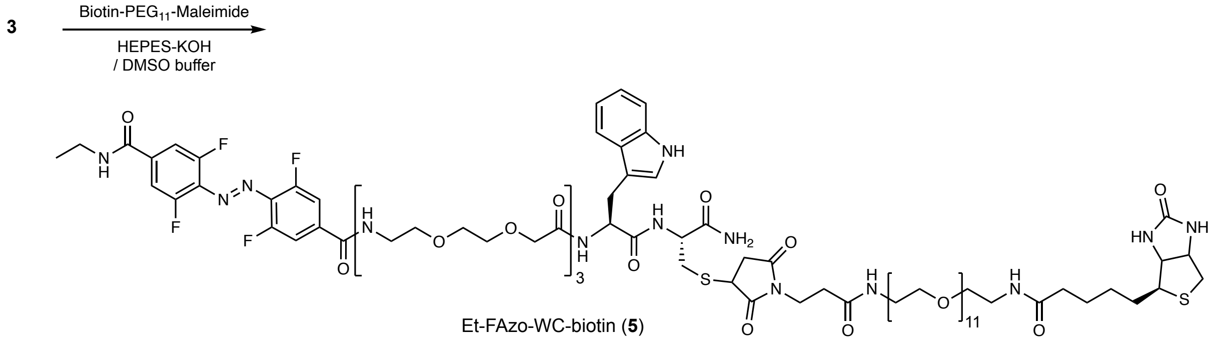


**Scheme S6.** Synthetic route of Et-FAzo-biotin (**5**)

Biotinylation of Et-FAzo-WC (**3**) was achieved by mixing 250 µM Et-FAzo-WC (**3**) with 250 µM Maleimide-PEG11-Biotin (Thermo Fisher Scientific) in 20 µL of 50 mM HEPES-KOH (pH 7.5) buffer containing 25% DMSO. The reaction mixture was incubated at 25°C for 2 h, and the reaction and the formation of the biotinylated target molecule, Et-FAzo-biotin (**5**), were confirmed by reversed-phase HPLC and MALDI-TOF mass spectrometry. The reaction solution was then diluted 50-fold with 50 mM HEPES-KOH buffer (pH 7.5) to achieve a final concentration of 5 µM Et-FAzo-biotin (**5**) and used directly in anticalin selection experiments without purification.
MALDI-TOF-MS (CHCA): calculated for $[M+H]^+$, 2014.862; found, 2014.013.

#### Synthesis of Ca-FAzo-WC-biotin (6)

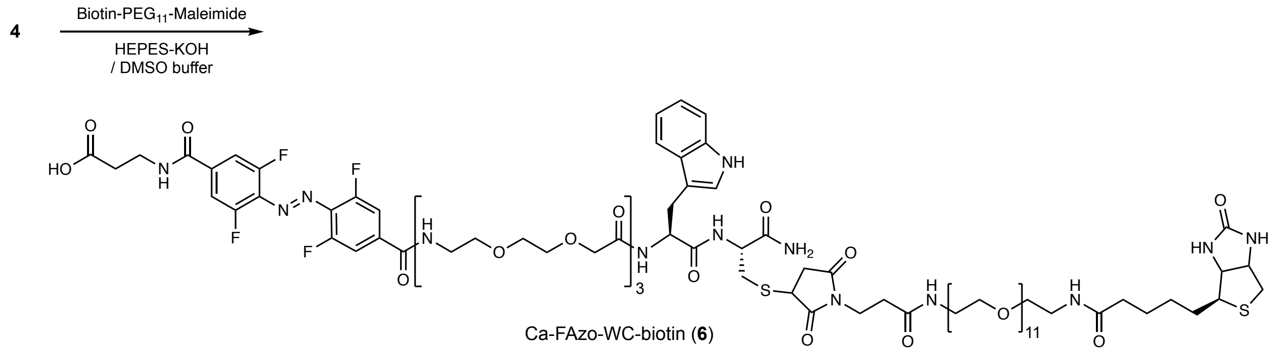


**Scheme S7.** Synthetic route of Ca-FAzo-biotin (**6**)

Ca-FAzo-biotin (**6**) was prepared from Ca-FAzo-WC (**4**) using the same procedure as described above. The reaction solution was also diluted 50-fold with 50 mM HEPES-KOH buffer (pH 7.5) and used directly in anticalin selection experiments without purification.

MALDI-TOF-MS (CHCA): calculated for $[M+H]^+$, 2058.851; found, 2058.978.

#### Synthesis of HBY-WC (18)

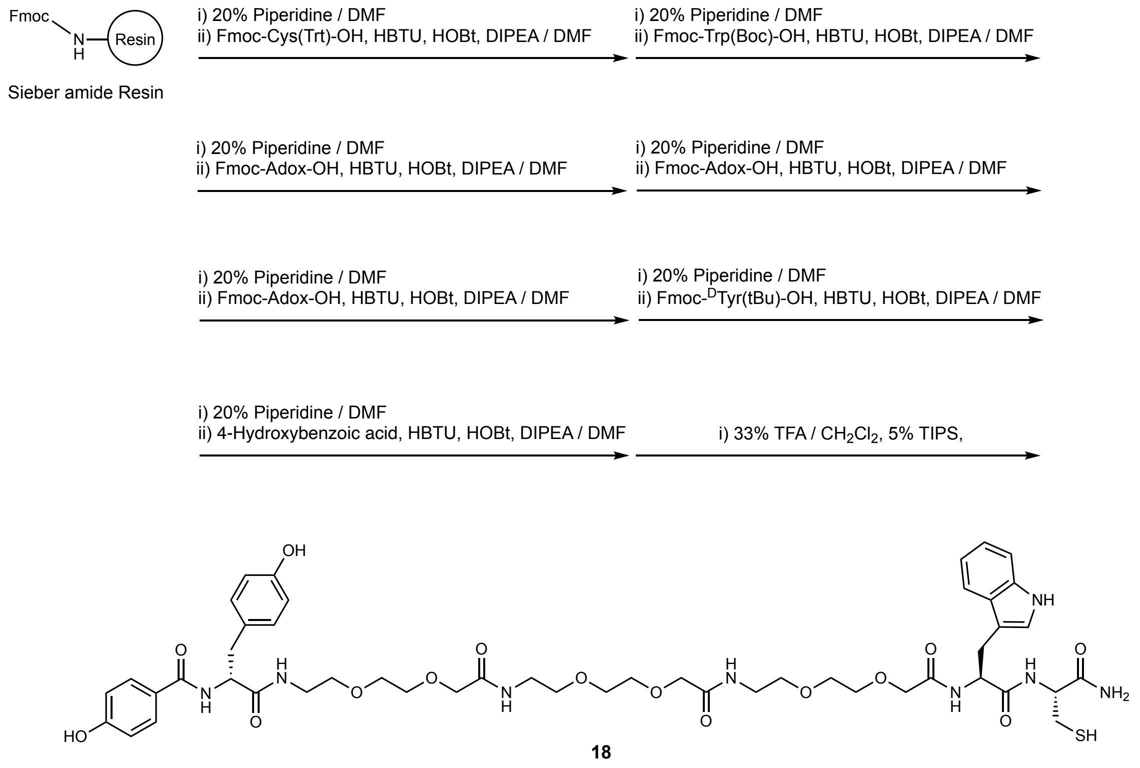


**Scheme S8.** Synthetic route of HBY-WC (**18**)

HBY-WC (**18**) was synthesized on Sieber Amide resin (0.66 mmol/g) (61 mg, 40 μmol). Fmoc-

Cys(Trt)-OH, Fmoc-Trp(Boc)-OH, Fmoc-Adox-OH (×3), and Fmoc-$^{D}$Tyr(tBu)-OH were sequentially coupled to the resin. Following Fmoc deprotection, 4-hydroxybenzoic acid was coupled to the N-terminus using a mixture of 4-hydroxybenzoic acid (8.3 mg, 60 μmol), HBTU (22 mg, 58 μmol), HOBt (9 mg, 59 μmol), and DIPEA (21 μL, 120 μmol) in DMF. Deprotection and cleavage from the resin were performed with $CH_2Cl_2$ (2 mL) containing 30% TFA and 5% TIPS (2 mL), followed by collection of the filtrate. The filtrate was concentrated under reduced pressure. The crude product was purified by reversed-phase HPLC using a semi-preparative ODS column with a linear gradient of MeCN containing 0.1% TFA and 0.1% aqueous TFA, yielding HBA-Y-WC (**18**) (4.44 mg, 22%) as a white solid.

$^{1}$H NMR (400 MHz, $CD_3OD$): δ 7.65 (2H, d, *J* = 8.80 Hz), 7.60 (1H, d, *J* = 7.60 Hz), 7.33 (1H, d, *J* = 8.04 Hz), 7.15 (1H, s), 7.10–7.00 (4H, m), 6.79 (2H, d, *J* = 8.80 Hz), 6.69 (2H, d, *J* = 8.56 Hz), 4.77–4.66 (2H, m), 4.48–4.43 (1H, m), 4.01–3.86 (6H, m), 3.62–3.38 (24H, m), 3.14–2.91 (2H, m), 2.84–2.71 (2H, m).

MALDI-TOF-MS (CHCA): calculated for $[M+Na]^+$, 1047.4104; found, 1047.3050.

**Synthesis of HBY-biotin (7)**

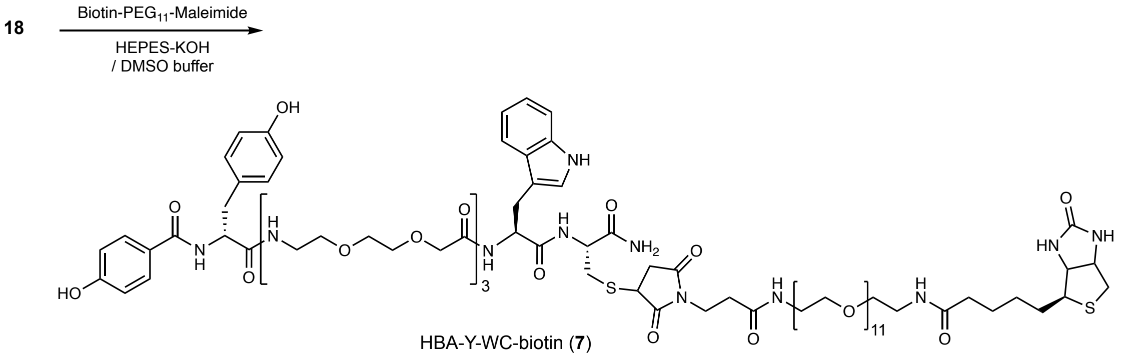


**Scheme S9.** Synthetic route of HBY-biotin (**7**)

HBA-Y-biotin (**7**) was prepared from HBA-Y-WC (**18**) following the same procedure as described in the **Synthesis of Et-FAzo-biotin (5)** section. The reaction solution was also diluted 50-fold with 50 mM HEPES-KOH buffer (pH 7.5) and used directly in anticalin selection experiments without purification.

MALDI-TOF-MS (CHCA): calculated for $[M+H]^+$, 1946.883; found, 1946.565.

#### Synthesis of Et-FAzo-HTL (8)

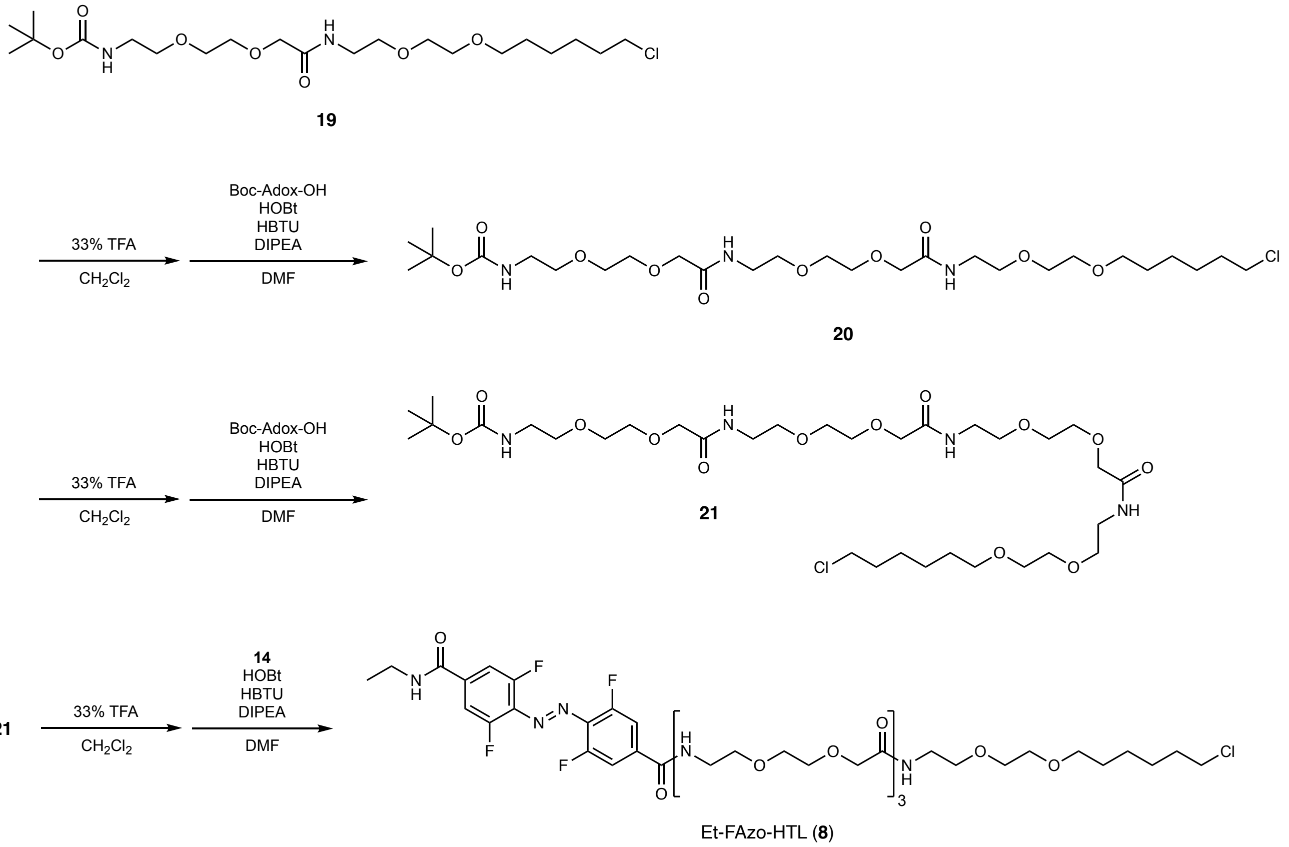

**Scheme S10.** Synthetic route of Et-FAzo-HTL (**8**)

Compound **20**

Compound **19**[S3] (100 mg, 214 μmol) was dissolved in a mixture of $CH_2Cl_2$ (6.0 mL) and TFA (3.0 mL) and stirred at room temperature for 1 h. The reaction mixture was then concentrated under reduced pressure. The residue was dissolved in toluene (3.0 mL) and concentrated again under reduced pressure. This process was repeated to remove TFA. The residue was then dissolved in anhydrous DMF (3.0 mL).

In another flask, Boc-Adox-OH (104 mg, 235 μmol) was dissolved in anhydrous DMF (2.0 mL) and stirred under an argon atmosphere. To this solution, DIPEA (186 μL, 1.07 mmol), HOBt (35.4 mg, 256 μmol), HBTU (97.3 mg, 256 μmol), and the DMF solution of the deprotected compound **19** prepared above were sequentially added. The resulting mixture was stirred at room temperature for 3 h and then concentrated under reduced pressure. The residue was dissolved in EtOAc (30 mL) and washed twice with saturated $NaHCO_3$ aqueous solution (30 mL each), twice with 5% citric acid aqueous solution (30 mL each), and finally once with brine (30 mL). The organic layer was dried over anhydrous $Na_2SO_4$, filtered, and concentrated under reduced pressure. The crude residue was purified by column chromatography (silica gel, $CHCl_3$/MeOH = 10:1), yielding compound **20** (123.6 mg, 83%) as a brown and transparent oil.

$^1$H NMR (400 MHz, $CDCl_3$): δ 4.01 (4H, s), 3.70–3.50 (24H, m), 3.45 (2H, t, $J$ = 5.40 Hz), 3.32 (2H, q, $J$ = 5.57 Hz), 1.81–1.75 (2H, m), 1.62–1.58 (4H, m), 1.44 (9H, s), 1.39–1.35 (2H, m).

Compound **21**

Compound **20** (100 mg, 163 μmol) was dissolved in a mixture of $CH_2Cl_2$ (4.0 mL) and TFA (2.0 mL) and stirred at room temperature for 1 h. The reaction mixture was then concentrated under reduced pressure. The residue was dissolved in toluene (3.0 mL) and concentrated again under reduced pressure. This process was repeated to remove TFA. The residue was then dissolved in anhydrous DMF (2.0 mL).

In another flask, Boc-Adox-OH (60.3 mg, 135 μmol) was dissolved in anhydrous DMF (3.0 mL) and stirred under an argon atmosphere. To this solution, DIPEA (235.2 μL, 1.35 mmol), HOBt (22.4 mg, 163 μmol), HBTU (61.9 mg, 163 μmol), and the DMF solution of the deprotected compound **20** prepared above were sequentially added. The resulting mixture was stirred at room temperature for 3 h and then concentrated under reduced pressure. The residue was dissolved in EtOAc (30 mL) and washed twice with saturated $NaHCO_3$ aqueous solution (30 mL each), twice with 5% citric acid aqueous solution (30 mL each), and finally once with brine (30 mL). The organic layer was dried over anhydrous $Na_2SO_4$, filtered, and concentrated under reduced pressure. The crude residue was purified by column chromatography (silica gel, $CHCl_3$/MeOH = 10:1), yielding compound **21** (71.8 mg, 61%) as a brown and transparent oil.

$^1$H NMR (400 MHz, $CDCl_3$): δ 4.00 (6H, s), 3.68–3.50 (32H, m), 3.45 (2H, t, $J$ = 6.66 Hz), 3.32 (2H, q, $J$ = 5.67 Hz), 1.81–1.75 (2H, m), 1.62–1.56 (4H, m), 1.44 (9H, s), 1.38–1.33 (2H, m).

Compound **8** (Et-FAzo-HTL)

Compound **21** (100 mg, 163 μmol) was dissolved in a mixture of $CH_2Cl_2$ (4.0 mL) and TFA (2.0 mL) and stirred at room temperature for 1 h. The reaction mixture was then concentrated under reduced pressure. The residue was dissolved in toluene (2.0 mL) and concentrated again under reduced pressure. This process was repeated to remove TFA. The residue was then dissolved in anhydrous DMF (1.0 mL).

In another flask, compound **14** (25 mg, 67 μmol) was dissolved in anhydrous DMF (1.0 mL) and stirred under an argon atmosphere. To this solution, DIPEA (75 μL, 435 μmol), HOBt (11.7 mg, 81 μmol), HBTU (30.7 mg, 81 μmol), and the DMF solution of the deprotected compound **21** prepared above were sequentially added. The resulting mixture was stirred at room temperature for 3 h and then concentrated under reduced pressure. The crude product was purified by reversed-phase HPLC using a semi-preparative ODS column with a linear gradient of MeCN containing 0.1% TFA and 0.1% aqueous TFA, yielding At-FAzo-HTL (**8**) (30.8 mg, 44%) as a red solid.

$^1$H NMR (400 MHz, $CD_3OD$): δ 7.71–7.65 (4H, m), 4.01–3.97 (6H, m), 3.73–3.34 (38H, m), 1.79–1.73 (2H, m), 1.61–1.57 (2H, m), 1.47–1.43 (2H, m), 1.39–1.35 (2H, m), 1.26 (3H, t, $J$ = 7.28 Hz).

HRMS (ESI): calculated for $[M+Na]^+$, 1032.4084; found, 1032.4073.

#### Synthesis of Ca-FAzo-HTL (9)

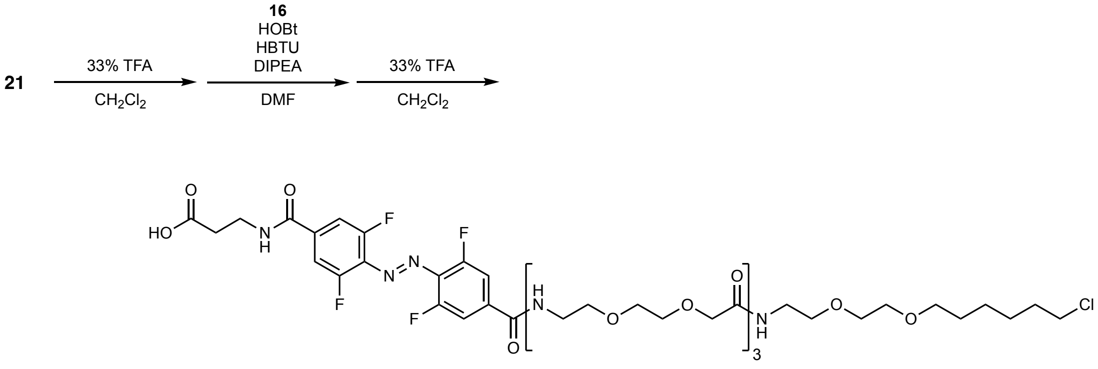


Ca-FAzo-HTL (**9**)

**Scheme S11.** Synthetic route of Ca-FAzo-HTL (**9**)

Compound **9** (Ca-FAzo-HTL)

Compound **21** (50 mg, 66 µmol) was dissolved in a mixture of $CH_2Cl_2$ (4.0 mL) and TFA (2.0 mL) and stirred at room temperature for 1 h. The reaction mixture was concentrated under reduced pressure. The residue was then dissolved in toluene (2.0 mL) and concentrated again under reduced pressure. This process was repeated to remove TFA. Finally, the residue was dissolved in anhydrous DMF (1.0 mL).

In another flask, compound **16** (28 mg, 60 µmol) was dissolved in anhydrous DMF (1.0 mL) and stirred under an argon atmosphere. To this solution, DIPEA (52 µL, 300 µmol), HOBt (9.7 mg, 72 µmol), HBTU (27.2 mg, 72 µmol), and the DMF solution of the deprotected compound **21** prepared above were sequentially added. The resulting mixture was stirred at room temperature for 3 h and then concentrated under reduced pressure. The residue was dissolved in $CH_2Cl_2$ (4.0 mL) and TFA (2.0 mL) and stirred at room temperature for 2 h. The reaction mixture was then concentrated under reduced pressure, and the residue was dissolved in toluene (2.0 mL) and concentrated again under reduced pressure. This process was repeated to remove TFA. The crude residue was purified by reversed-phase HPLC using a semi-preparative ODS column with a linear gradient of MeCN containing 0.1% TFA and 0.1% aqueous TFA, yielding Ca-FAzo-HTL (**9**) (15.7 mg, 25%) as a red solid.

$^1$H-NMR (400 MHz, $CD_3OD$): δ 7.70–7.64 (4H, m), 4.01–3.96 (6H, m), 3.71–3.40 (38H, m), 2.65 (2H, t, $J$ = 6.84 Hz), 1.79–1.72 (2H, m), 1.61–1.54 (2H, m), 1.49–1.43 (2H, m), 1.41–1.34 (2H, m).

HRMS (ESI): calculated for $[M+Na]^+$, 1076.3987; found, 1076.3971.

#### Synthesis of Ca(AM)-FAzo-HTL (10)

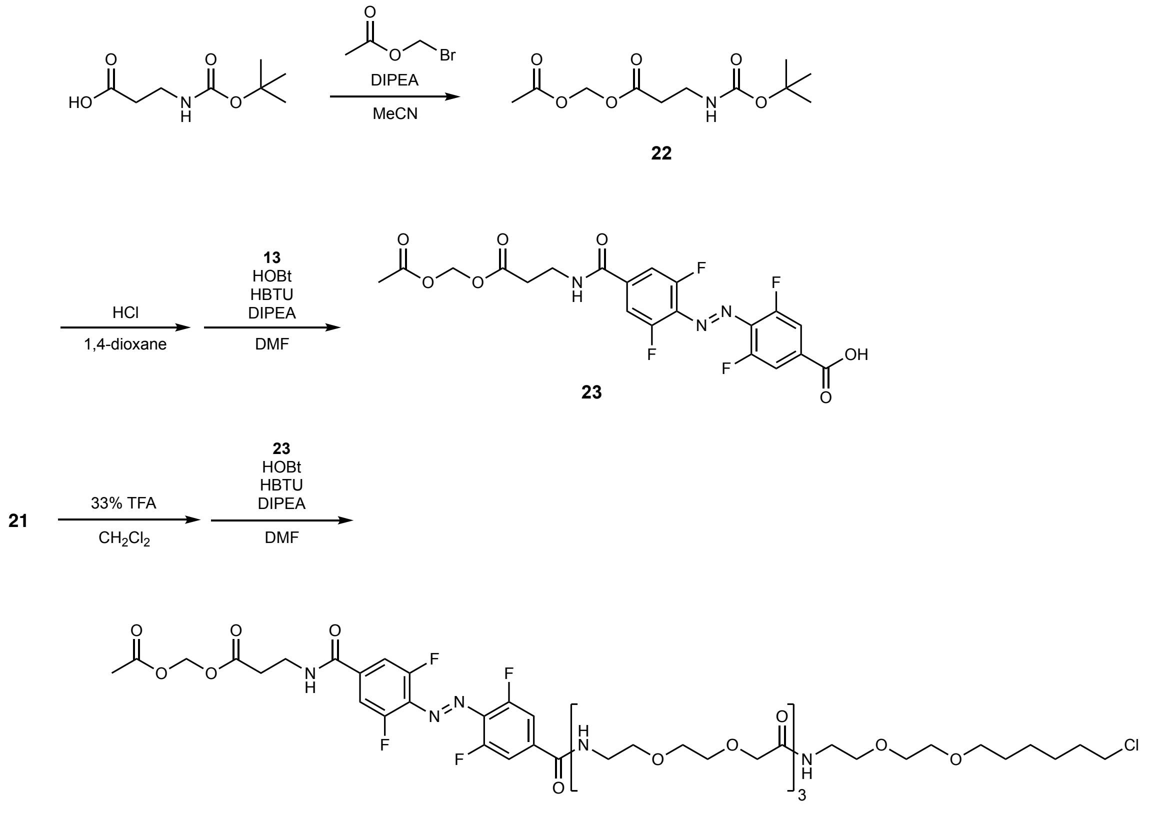

Ca(AM)-FAzo-HTL (**10**)

**Scheme S12.** Synthetic route of Ca(AM)-FAzo-HTL (**10**)

Compound **22**

*N*-(*tert*-butoxycarbonyl)-*β*-alanine (500 mg, 2.6 mmol) was dissolved in anhydrous MeCN (10 mL) and stirred under an argon atmosphere. DIPEA (1.15 mL, 6.6 mmol) and bromomethyl acetate (607 mg, 4.0 mmol) were then added. The resulting mixture was stirred at room temperature for 10 h and then concentrated under reduced pressure. The residue was dissolved in EtOAc (50 mL) and washed twice with 5% citric acid aqueous solution (50 mL each) and brine (30 mL). The organic layer was dried over anhydrous $Na_2SO_4$, filtered, and concentrated under reduced pressure. The crude residue was purified by column chromatography (silica gel, hexane/EtOAc = 1:1), yielding compound **22** (658.1 mg, 95%) as a colorless and transparent oil.

$^1$H NMR (400 MHz, $CDCl_3$): δ 5.76 (2H, s), 4.99 (1H, br s), 3.42 (2H, q, $J$ = 5.91 Hz), 2.59 (2H, t, $J$ = 6.04 Hz), 2.13 (3H, s), 1.44 (9H, s).

Compound **23**

Compound **22** (30 mg, 116 μmol) was dissolved in a mixture of 1,4-dioxane (1.3 mL) and 4 M HCl in 1,4-dioxane (1.0 mL) and stirred at room temperature for 3 h. The reaction mixture was then concentrated under reduced pressure. The residue was dissolved in 1,4-dioxane (2.0 mL) and concentrated again under reduced pressure. This process was repeated to remove HCl. The residue was then dissolved in anhydrous DMF (0.5 mL).

In another flask, compound **13** (30 mg, 87 µmol) was dissolved in anhydrous DMF (0.5 mL) and stirred under an argon atmosphere. DIPEA (73.7 µL, 435 µmol), HOBt (13.2 mg, 96 µmol), HBTU (36.4 mg, 96 µmol), and the DMF solution of the deprotected compound **22** prepared above were sequentially added. The resulting mixture was stirred at room temperature for 3 h and then concentrated under reduced pressure. The crude residue was purified by column chromatography (silica gel, $CHCl_3$/MeOH/AcOH = 100:10:1), yielding compound **23** (13.6 mg, 32%) as a red solid.
$^1$H NMR (400 MHz, DMSO-*d₆*): δ 14.05 (1H, s), 8.93 (1H, t, *J* = 5.36 Hz), 7.81–7.78 (4H, m), 5.69 (2H, s), 3.53 (2H, q, *J* = 6.20 Hz), 2.70 (2H, t, *J* = 6.66 Hz), 2.06 (3H, s).

Compound **10** [Ca(AM)-FAzo-HTL]
Compound **21** (100 mg, 131.9 µmol) was dissolved in a mixture of $CH_2Cl_2$ (5.0 mL) and TFA (2.5 mL) and stirred at room temperature for 1 h. The mixture was concentrated under reduced pressure. The residue was dissolved in toluene (2.0 mL) and concentrated again under reduced pressure. This process was repeated to remove TFA. The residue was then dissolved in anhydrous DMF (1.5 mL).

In another flask, compound **23** (70.3 mg, 145.1 µmol) was dissolved in anhydrous DMF (1.0 mL) and stirred under an argon atmosphere. DIPEA (115 µL, 659 µmol), HOBt (27.3 mg, 197.8 µmol), HBTU (75.1 mg, 197.8 µmol), and the DMF solution of the deprotected compound **21** prepared above were sequentially added. The resulting mixture was stirred at room temperature for 3 h and then concentrated under reduced pressure. The crude residue was purified by reversed-phase HPLC using a semi-preparative ODS column with a linear gradient of MeCN containing 0.1% TFA and 0.1% aqueous TFA, yielding Ca(AM)-FAzo-HTL (**10**) (53.7 mg, 35%) as a red solid.
$^1$H NMR (400 MHz, $CD_3OD$): δ 7.70–7.64 (4H, m), 5.75 (2H, s), 4.01–3.97 (6H, m), 3.71–3.39 (38H, m), 2.73 (2H, t, *J* = 6.72 Hz), 2.06 (3H, s), 1.79
–1.73 (2H, m), 1.61–1.57 (2H, m), 1.47–1.43 (2H, m), 1.39–1.35 (2H, m).
HRMS (ESI): calculated for $[M+Na]^+$, 1148.4194; found, 1148.4207.

#### Supplementary Methods: Preparation of ligand-immobilized beads

Et-FAzo– and Ca-FAzo–immobilized beads for the first selection round were prepared as follows. An aliquot of 100 µL (suspension volume) of Dynabeads M270 Amine (Invitrogen) was washed five times with DMF. A 100 µL DMF solution containing 1 M *N*,*N*'-diisopropylcarbodiimide and 1 M bromoacetic acid was added to the beads. The suspension was incubated at 25°C for 5 min to introduce bromoacetyl groups onto the amino groups on the surface of the beads, after which the beads were washed five times with DMF. For the preparation of Et-FAzo–immobilized beads, a 25 µL solution containing 100 mM HEPES-KOH (pH 8.0), 600 mM NaCl, 50% DMF, and 0.1 mM of Et-FAzo-WC (**3**) was added to the beads. The suspension was incubated at 37°C for 20 min to immobilize the Et-FAzo ligand. The resultant beads were washed once with DMF and then divided in half. A 50 µL solution containing 100 mM HEPES-KOH (pH 8.0), 600 mM NaCl, 50% DMF, and 1 mM PEG2000-thiol (NOF Corporation) was added to one half of the beads. The suspension was incubated at 37°C for 2 h to immobilize PEG2000 onto the Et-FAzo-modified beads. After washing the resultant Et-FAzo/PEG–immobilized beads (Et-FAzo-beads/+PEG) and the Et-FAzo–immobilized beads (the remaining half described above; Et-FAzo-beads/–PEG) with DMF, a 50 µL solution containing 100 mM HEPES-KOH (pH 8.0), 600 mM NaCl, 50% DMF, and 500 mM 1-thioglycerol was added, and the suspensions were incubated at 37°C for 1 min to modify the bromoacetyl groups. The resultant beads were washed once with DMF and once with 10 mM Tris-HCl (pH 8.0), and then resuspended in a 50 µL solution containing 10 mM AcONa and 1 mM DTT. For the preparation of Ca-FAzo–immobilized beads (Ca-FAzo-beads/+PEG and Ca-FAzo-beads/–PEG), Ca-FAzo-WC (**4**) was used instead of Et-FAzo-WC following the same procedure.

Control beads for negative selections were prepared as follows. HBY-immobilized M280 beads were prepared by adding 6 µL of 5 µM HBY-biotin (**5**) solution to 30 µL (suspension volume) of Dynabeads M280 Streptavidin (Invitrogen). HBY-immobilized M270 beads were prepared by adding 2 µL of 5 µM HBY-biotin solution to 2.4 µL (suspension volume) of Dynabeads M270 Streptavidin (Invitrogen). The solutions were mixed by pipetting for 1 min. The beads were washed twice with HBST buffer [50 mM HEPES-KOH (pH 7.5), 300 mM NaCl, 0.05% Tween 20], and the supernatant was removed just before use.

#### Supplementary Sequences

**pT5-AzoTag1-NusA-His6**

>Amino acid sequence
MTNRSGSQDSTSDLIPAPPLSKVPLQQNFQDNQFQGKWYQVGRAGNFNLREDKDPTKMQAIIYELKEDKSYNVTPVYFRKK
KCQYNIDTFVPGSQPGEFTLGNIKSYPGNTSELVRVVSTNYNQHAMVFVKWVYQNREWFSIYLYGRTKELTSELKENFIRF
SKSLGLPENHIVFPVPIDQCIDGTSGGGGENLYFQGSGELNKEILAVVEAVSNEKALPREKIFEALESALATATKKKYEQE
IDVRVQIDRKSGDFDTFRRWLVVDEVTQPTKEITLEAARYEDESLNLGDYVEDQIESVTFDRITTQTAKQVIVQKVREAER
AMVVDQFREHEGEIITGVVKKVNRDNISLDLGNNAEAVILREDMLPRENFRPGDRVRGVLYSVRPEARGAQLFVTRSKPEM
LIELFRIEVPEIGEEVIEIKAAARDPGSRAKIAVKTNDKRIDPVGACVGMRGARVQAVSTELGGERIDIVLWDDNPAQFVI
NAMAPADVASIVVDEDKHTMDIAVEAGNLAQAIGRNGQNVRLASQLSGWELNVMTVDDLQAKHQAEAHAAIDTFTKYLDID
EDFATVLVEEGFSTLEELAYVPMKELLEIEGLDEPTVEALRERAKNALATIAQAQEESLGDNKPADDLLNLEGVDRDLAFK
LAARGVCTLEDLAEQGIDDLADIEGLTDEKAGALIMAARNICWFGDEAGSGSGSGSLPETGGGSGHHHHHH*
>DNA sequence
ATGACAAATCGATCTGGTTCTCAAGATAGTACCAGCGATCTGATTCCGGCGCCGCCGCTGAGCAAGGTGCCGCTGCAGCAA
AACTTCCAAGACAACCAGTTTCAGGGTAAATGGTACCAGGTTGGTCGTGCGGGCAACTTCAACCTGCGTGAGGACAAGGAC
CCGACTAAAATGCAGGCGATCATTTATGAGCTGAAGGAAGACAAAAGCTACAACGTGACCCCGGTTTACTTCCGTAAGAAG
AAATGCCAGTATAACATTGACACCTTCGTGCCGGGTAGCCAACCGGGCGAATTTACCCTGGGTAACATTAAGAGCTACCCG
GGCAACACCAGCGAACTGGTTCGTGTGGTTAGCACCAACTATAACCAGCACGCGATGGTGTTCGTTAAGTGGGTTTACCAA
AACCGTGAGTGGTTCTCTATTTACCTGTATGGCCGTACCAAGGAACTGACCAGCGAGCTGAAAGAAAACTTCATCCGTTTT
AGCAAAAGCCTGGGTCTGCCGGAAAACCACATTGTGTTTCCGGTTCCGATCGACCAATGCATTGATGGCACTAGTGGCGGC
GGAGGAGAGAACCTGTACTTTCAGGGTTCAGGAGAGCTCAACAAAGAAATTTTGGCTGTAGTTGAAGCCGTATCCAATGAA
AAGGCGCTACCTCGCGAGAAGATTTTCGAAGCATTGGAAAGCGCGCTGGCGACAGCAACAAAGAAAAAATATGAACAAGAG
ATCGACGTCCGCGTACAGATCGATCGCAAAAGCGGTGATTTTGACACTTTCCGTCGCTGGTTAGTTGTTGATGAAGTCACC
CAGCCGACCAAGGAAATCACCCTTGAAGCCGCACGTTATGAAGATGAAAGCCTGAACCTGGGCGATTACGTTGAAGATCAG
ATTGAGTCTGTTACCTTTGACCGTATCACTACCCAGACGGCAAAACAGGTTATCGTGCAGAAAGTGCGTGAAGCCGAACGT
GCGATGGTGGTTGATCAGTTCCGTGAACACGAAGGTGAAATCATCACCGGCGTGGTGAAAAAAGTAAACCGCGACAACATC
TCTCTGGATCTGGGCAACAACGCTGAAGCCGTGATCCTGCGCGAAGATATGCTGCCGCGTGAAAACTTCCGCCCTGGCGAC
CGCGTTCGTGGCGTGCTCTATTCCGTTCGCCCGGAAGCGCGTGGCGCGCAACTGTTCGTCACTCGTTCCAAGCCGGAAATG
CTGATCGAACTGTTCCGTATTGAAGTGCCAGAAATCGGCGAAGAAGTGATTGAAATTAAAGCAGCGGCTCGCGATCCGGGT
TCTCGTGCGAAAATCGCGGTGAAAACCAACGATAAACGTATCGATCCGGTAGGTGCTTGCGTAGGTATGCGTGGCGCGCGT
GTTCAGGCGGTGTCTACTGAACTGGGTGGCGAGCGTATCGATATCGTCCTGTGGGATGATAACCCGGCGCAGTTCGTGATT
AACGCAATGGCACCGGCAGACGTTGCTTCTATCGTGGTGGATGAAGATAAACACACCATGGACATCGCCGTTGAAGCCGGT
AATCTGGCGCAGGCGATTGGCCGTAACGGTCAGAACGTGCGTCTGGCTTCGCAACTGAGCGGTTGGGAACTCAACGTGATG
ACCGTTGACGACCTGCAAGCTAAGCATCAGGCGGAAGCGCACGCAGCGATCGACACCTTCACCAAATATCTCGACATCGAC
GAAGACTTCGCGACTGTTCTGGTAGAAGAAGGCTTCTCGACGCTGGAAGAATTGGCCTATGTGCCGATGAAAGAGCTGTTG
GAAATCGAAGGCCTTGATGAGCCGACCGTTGAAGCACTGCGCGAGCGTGCTAAAAATGCACTGGCCACCATTGCACAGGCC
CAGGAAGAAAGCCTCGGTGATAACAAACCGGCTGACGATCTGCTGAACCTTGAAGGGGTAGATCGTGATTTGGCATTCAAA
CTGGCCGCCCGTGGCGTTTGTACGCTGGAAGATCTCGCCGAACAGGGCATTGATGATCTGGCTGATATCGAAGGGTTGACC
GACGAAAAAGCCGGAGCACTGATTATGGCTGCCCGTAATATTTGCTGGTTCGGTGACGAAGCGGGATCCGGTAGTGGATCT
GGATCACTGCCGGAAACCGGCGGTGGCAGCGGTCACCATCATCACCACCACTAA
>AzoTag1 NusA His6

**pT5-AzoTag16-NusA-His6**

>Amino acid sequence
MTNRSGSQDSTSDLIPAPPLSKVPLQQNFQDNQFQGKWYIVGYAGNDFLREDKDPFKMWARIYELKEDKSYNVTGVFFRKK
KCYYHIHTFVPGSQPGEFTLGNIKSYPGYTSQLVRVVSTNYNQHAMVFTKYVYQNREMFYIVLYGRTKELTSELKENFIRF
SKSLGLPENHIVFPVPIDQCIDGTSGGGGENLYFQGSGELNKEILAVVEAVSNEKALPREKIFEALESALATATKKKYEQE
IDVRVQIDRKSGDFDTFRRWLVVDEVTQPTKEITLEAARYEDESLNLGDYVEDQIESVTFDRITTQTAKQVIVQKVREAER
AMVVDQFREHEGEIITGVVKKVNRDNISLDLGNNAEAVILREDMLPRENFRPGDRVRGVLYSVRPEARGAQLFVTRSKPEM
LIELFRIEVPEIGEEVIEIKAAARDPGSRAKIAVKTNDKRIDPVGACVGMRGARVQAVSTELGGERIDIVLWDDNPAQFVI
NAMAPADVASIVVDEDKHTMDIAVEAGNLAQAIGRNGQNVRLASQLSGWELNVMTVDDLQAKHQAEAHAAIDTFTKYLDID
EDFATVLVEEGFSTLEELAYVPMKELLEIEGLDEPTVEALRERAKNALATIAQAQEESLGDNKPADDLLNLEGVDRDLAFK
LAARGVCTLEDLAEQGIDDLADIEGLTDEKAGALIMAARNICWFGDEAGSGSGSGSLPETGGGSGHHHHHH*
>DNA sequence
ATGACAAATCGATCTGGTTCTCAAGATAGTACCAGCGATCTGATTCCGGCGCCGCCGCTGAGCAAGGTGCCGCTGCAGCAA
AACTTCCAAGACAACCAGTTTCAGGGTAAATGGTACATCGTTGGTTACGCGGGCAACGACTTCCTGCGTGAGGACAAGGAC
CCGTTCAAAATGTGGGCGCGTATTTATGAGCTGAAGGAAGACAAAAGCTACAACGTGACCGGTGTTTTCTTCCGTAAGAAG
AAATGCTACTATCATATTCATACCTTCGTGCCGGGTAGCCAACCGGGCGAATTTACCCTGGGTAACATTAAGAGCTACCCG
GGCTACACCAGCCAGCTGGTTCGTGTGGTTAGCACCAACTATAACCAGCACGCGATGGTGTTCACTAAGTACGTTTACCAA
AACCGTGAGATGTTCTACATTGTTCTGTATGGCCGTACCAAGGAACTGACCAGCGAGCTGAAAGAAAACTTCATCCGTTTT
AGCAAAAGCCTGGGTCTGCCGGAAAACCACATTGTGTTTCCGGTTCCGATCGACCAATGCATTGATGGCACTAGTGGCGGC
GGAGGAGAGAACCTGTACTTTCAGGGTTCAGGAGAGCTCAACAAAGAAATTTTGGCTGTAGTTGAAGCCGTATCCAATGAA
AAGGCGCTACCTCGCGAGAAGATTTTCGAAGCATTGGAAAGCGCGCTGGCGACAGCAACAAAGAAAAAATATGAACAAGAG
ATCGACGTCCGCGTACAGATCGATCGCAAAAGCGGTGATTTTGACACTTTCCGTCGCTGGTTAGTTGTTGATGAAGTCACC
CAGCCGACCAAGGAAATCACCCTTGAAGCCGCACGTTATGAAGATGAAAGCCTGAACCTGGGCGATTACGTTGAAGATCAG
ATTGAGTCTGTTACCTTTGACCGTATCACTACCCAGACGGCAAAACAGGTTATCGTGCAGAAAGTGCGTGAAGCCGAACGT
GCGATGGTGGTTGATCAGTTCCGTGAACACGAAGGTGAAATCATCACCGGCGTGGTGAAAAAAGTAAACCGCGACAACATC

TCTCTGGATCTGGGCAACAACGCTGAAGCCGTGATCCTGCGCGAAGATATGCTGCCGCGTGAAAACTTCCGCCCTGGCGAC
CGCGTTCGTGGCGTGCTCTATTCCGTTCGCCCGGAAGCGCGTGGCGCGCAACTGTTCGTCACTCGTTCCAAGCCGGAAATG
CTGATCGAACTGTTCCGTATTGAAGTGCCAGAAATCGGCGAAGAAGTGATTGAAATTAAAGCAGCGGCTCGCGATCCGGGT
TCTCGTGCGAAAATCGCGGTGAAAACCAACGATAAACGTATCGATCCGGTAGGTGCTTGCGTAGGTATGCGTGGCGCGCGT
GTTCAGGCGGTGTCTACTGAACTGGGTGGCGAGCGTATCGATATCGTCCTGTGGGATGATAACCCGGCGCAGTTCGTGATT
AACGCAATGGCACCGGCAGACGTTGCTTCTATCGTGGTGGATGAAGATAAACACACCATGGACATCGCCGTTGAAGCCGGT
AATCTGGCGCAGGCGATTGGCCGTAACGGTCAGAACGTGCGTCTGGCTTCGCAACTGAGCGGTTGGGAACTCAACGTGATG
ACCGTTGACGACCTGCAAGCTAAGCATCAGGCGGAAGCGCACGCAGCGATCGACACCTTCACCAAATATCTCGACATCGAC
GAAGACTTCGCGACTGTTCTGGTAGAAGAAGGCTTCTCGACGCTGGAAGAATTGGCCTATGTGCCGATGAAAGAGCTGTTG
GAAATCGAAGGCCTTGATGAGCCGACCGTTGAAGCACTGCGCGAGCGTGCTAAAAATGCACTGGCCACCATTGCACAGGCC
CAGGAAGAAAGCCTCGGTGATAACAAACCGGCTGACGATCTGCTGAACCTTGAAGGGGTAGATCGTGATTTGGCATTCAAA
CTGGCCGCCCGTGGCGTTTGTACGCTGGAAGATCTCGCCGAACAGGGCATTGATGATCTGGCTGATATCGAAGGGTTGACC
GACGAAAAAGCCGGAGCACTGATTATGGCTGCCCGTAATATTTGCTGGTTCGGTGACGAAGCGGGATCCGGTAGTGGATCT
GGATCACTGCCGGAAACCGGCGGTGGCAGCGGTCACCATCATCACCACCACTAA
>AzoTag16 NusA His6

**pCAGGS-AzoTag1-miRFP670-NES-P2A-HaloTag-KRas4B(CT)**

>Amino acid sequence
MTNRSGSQDSTSDLIPAPPLSKVPLQQNFQDNQFQGKWYQVGRAGNFNLREDKDPTKMQAIIYELKEDKSYNVTPVYFRKK
KCQYNIDTFVPGSQPGEFTLGNIKSYPGNTSELVRVVSTNYNQHAMVFVKWVYQNREWFSIYLYGRTKELTSELKENFIRF
SKSLGLPENHIVFPVPIDQCIDGASGGSGGSPVATMVAGHASGSPAFGTASHSNCEHEEIHLAGSIQPHGALLVVSEHDHR
VIQASANAAEFLNLGSVLGVPLAEIDGDLLIKILPHLDPTAEGMPVAVRCRIGNPSTEYCGLMHRPPEGGLIIELERAGPS
IDLSGTLAPALERIRTAGSLRALCDDTVLLFQQCTGYDRVMVYRFDEQGHGLVFSECHVPGLESYFGNRYPSSTVPQMARQ
LYVRQRVRVLVDVTYQPVPLEPRLSPLTGRDLDMSGCFLRSMSPCHLQFLKDMGVRATLAVSLVVGGKLWGLVVCHHYLPR
FIRFELRAICKRLAERIATRITALESLYKGGSNELALKLAGLDINKTGGTGSGATNFSLLKQAGDVEENPGPQLMAEIGTG
FPFDPHYVEVLGERMHYVDVGPRDGTPVLFLHGNPTSSYVWRNIIPHVAPTHRCIAPDLIGMGKSDKPDLGYFFDDHVRFM
DAFIEALGLEEVVLVIHDWGSALGFHWAKRNPERVKGIAFMEFIRPIPTWDEWPEFARETFQAFRTTDVGRKLIIDQNVFI
EGTLPMGVVRPLTEVEMDHYREPFLNPVDREPLWRFPNELPIAGEPANIVALVEEYMDWLHQSPVPKLLFWGTPGVLIPPA
EAARLAKSLPNCKAVDIGPGLNLLQEDNPDLIGSEIARWLSTLEISGGSGASAGGGSLEISGGKKKKKKSKTKCVIM*
>DNA sequence
ATGACAAATCGATCTGGTTCTCAAGATAGTACCAGCGATCTGATTCCGGCGCCGCCGCTGAGCAAGGTGCCGCTGCAGCAA
AACTTCCAAGACAACCAGTTTCAGGGTAAATGGTACCAGGTTGGTCGTGCGGGCAACTTCAACCTGCGTGAGGACAAGGAC
CCGACTAAAATGCAGGCGATCATTTATGAGCTGAAGGAAGACAAAAGCTACAACGTGACCCCGGTTTACTTCCGTAAGAAG
AAATGCCAGTATAACATTGACACCTTCGTGCCGGGTAGCCAACCGGGCGAATTTACCCTGGGTAACATTAAGAGCTACCCG
GGCAACACCAGCGAACTGGTTCGTGTGGTTAGCACCAACTATAACCAGCACGCGATGGTGTTCGTTAAGTGGGTTTACCAA
AACCGTGAGTGGTTCTCTATTTACCTGTATGGCCGTACCAAGGAACTGACCAGCGAGCTGAAAGAAAACTTCATCCGTTTT
AGCAAAAGCCTGGGTCTGCCGGAAAACCACATTGTGTTTCCGGTTCCGATCGACCAATGCATTGATGGCGCTAGCGGTGGT
AGTGGTGGTTCACCGGTCGCCACCATGGTAGCAGGTCATGCCTCTGGCAGCCCCGCATTCGGGACCGCCTCTCATTCGAAT
TGCGAACATGAAGAGATCCACCTCGCCGGCTCGATCCAGCCGCATGGCGCGCTTCTGGTCGTCAGCGAACATGATCATCGC
GTCATCCAGGCCAGCGCCAACGCCGCGGAATTTCTGAATCTCGGAAGCGTACTCGGCGTTCCGCTCGCCGAGATCGACGGC
GATCTGTTGATCAAGATCCTGCCGCATCTCGATCCCACCGCCGAAGGCATGCCGGTCGCGGTGCGCTGCCGGATCGGCAAT
CCCTCTACGGAGTACTGCGGTCTGATGCATCGGCCTCCGGAAGGCGGGCTGATCATCGAACTCGAACGTGCCGGCCCGTCG
ATCGATCTGTCAGGCACGCTGGCGCCGGCGCTGGAGCGGATCCGCACGGCGGGTTCACTGCGCGCGCTGTGCGATGACACC
GTGCTGCTGTTTCAGCAGTGCACCGGCTACGACCGGGTGATGGTGTATCGTTTCGATGAGCAAGGCCACGGCCTGGTATTC
TCCGAGTGCCATGTGCCTGGGCTCGAATCCTATTTCGGCAACCGCTATCCGTCGTCGACTGTCCCGCAGATGGCGCGGCAG
CTGTACGTGCGGCAGCGCGTCCGCGTGCTGGTCGACGTCACCTATCAGCCGGTGCCGCTGGAGCCGCGGCTGTCGCCGCTG
ACCGGGCGCGATCTCGACATGTCGGGCTGCTTCCTGCGCTCGATGTCGCCGTGCCATCTGCAGTTCCTGAAGGACATGGGC
GTGCGCGCCACCCTGGCGGTGTCGCTGGTGGTCGGCGGCAAGCTGTGGGGCCTGGTTGTCTGTCACCATTATCTGCCGCGC
TTCATCCGTTTCGAGCTGCGGGCGATCTGCAAACGGCTCGCCGAAAGGATCGCGACGCGGATCACCGCGCTTGAGAGCCTG
TACAAGGGCGGCAGCAACGAGCTGGCCCTGAAGCTGGCCGGCCTGGACATCAACAAGACCGGCGGTACCGGAAGCGGAGCT
ACTAACTTCAGCCTGCTGAAGCAGGCTGGAGACGTGGAGGAGAACCCTGGACCTCAATTGATGGCAGAAATCGGTACTGGC
TTTCCATTCGACCCCCATTATGTGGAAGTCCTGGGCGAGCGCATGCACTACGTCGATGTTGGTCCGCGCGATGGCACCCCT
GTGCTGTTCCTGCACGGTAACCCGACCTCCTCCTACGTGTGGCGCAACATCATCCCGCATGTTGCACCGACCCATCGCTGC
ATTGCTCCAGACCTGATCGGTATGGGCAAATCCGACAAACCAGACCTGGGTTATTTCTTCGACGACCACGTCCGCTTCATG
GATGCCTTCATCGAAGCCCTGGGTCTGGAAGAGGTCGTCCTGGTCATTCACGACTGGGGCTCCGCTCTGGGTTTCCACTGG
GCCAAGCGCAATCCAGAGCGCGTCAAAGGTATTGCATTTATGGAGTTCATCCGCCCTATCCCGACCTGGGACGAATGGCCA
GAATTTGCCCGCGAGACCTTCCAGGCCTTCCGCACCACCGACGTCGGCCGCAAGCTGATCATCGATCAGAACGTTTTTATC
GAGGGTACGCTGCCGATGGGTGTCGTCCGCCCGCTGACTGAAGTCGAGATGGACCATTACCGCGAGCCGTTCCTGAATCCT
GTTGACCGCGAGCCACTGTGGCGCTTCCCAAACGAGCTGCCAATCGCCGGTGAGCCAGCGAACATCGTCGCGCTGGTCGAA
GAATACATGGACTGGCTGCACCAGTCCCCTGTCCCGAAGCTGCTGTTCTGGGGCACCCCAGGCGTTCTGATCCCACCGGCC
GAAGCCGCTCGCCTGGCCAAAAGCCTGCCTAACTGCAAGGCTGTGGACATCGGCCCGGGTCTGAATCTGCTGCAAGAAGAC
AACCCGGACCTGATCGGCAGCGAGATCGCGCGCTGGCTGTCGACGCTCGAGATTTCCGGCGGCTCCGGTGCCAGTGCTGGT
GGTGGCAGCCTCGAGATTTCCGGCGGGAAGAAAAAGAAGAAGAAGTCCAAGACAAAATGCGTGATTATGTAG
>AzoTag1 miRFP670 NES(nuclear export signal) P2A HaloTag KRas4B(CT)(amino acids 174-188 of KRas4B)

**pCAGGS-AzoTag16-miRFP670-NES-P2A-HaloTag-KRas4B(CT)**

>Amino acid sequence
MTNRSGSQDSTSDLIPAPPLSKVPLQQNFQDNQFQGKWYIVGYAGNDFLREDKDPFKMWARIYELKEDKSYNVTGVFFRKK
KCYYHIHTFVPGSQPGEFTLGNIKSYPGYTSQLVRVVSTNYNQHAMVFTKYVYQNREMFYIVLYGRTKELTSELKENFIRF
SKSLGLPENHIVFPVPIDQCIDGASGGSGGSPVATMVAGHASGSPAFGTASHSNCEHEEIHLAGSIQPHGALLVVSEHDHR
VIQASANAAEFLNLGSVLGVPLAEIDGDLLIKILPHLDPTAEGMPVAVRCRIGNPSTEYCGLMHRPPEGGLIIELERAGPS
IDLSGTLAPALERIRTAGSLRALCDDTVLLFQQCTGYDRVMVYRFDEQGHGLVFSECHVPGLESYFGNRYPSSTVPQMARQ
LYVRQRVRVLVDVTYQPVPLEPRLSPLTGRDLDMSGCFLRSMSPCHLQFLKDMGVRATLAVSLVVGGKLWGLVVCHHYLPR
FIRFELRAICKRLAERIATRITALESLYKGGSNELALKLAGLDINKTGGTGSGATNFSLLKQAGDVEENPGPQLMAEIGTG
FPFDPHYVEVLGERMHYVDVGPRDGTPVLFLHGNPTSSYVWRNIIPHVAPTHRCIAPDLIGMGKSDKPDLGYFFDDHVRFM
DAFIEALGLEEVVLVIHDWGSALGFHWAKRNPERVKGIAFMEFIRPIPTWDEWPEFARETFQAFRTTDVGRKLIIDQNVFI
EGTLPMGVVRPLTEVEMDHYREPFLNPVDREPLWRFPNELPIAGEPANIVALVEEYMDWLHQSPVPKLLFWGTPGVLIPPA
EAARLAKSLPNCKAVDIGPGLNLLQEDNPDLIGSEIARWLSTLEISGGSGASAGGGSLEISGGKKKKKKSKTKCVIM*
>DNA sequence
ATGACAAATCGATCTGGTTCTCAAGATAGTACCAGCGATCTGATTCCGGCGCCGCCGCTGAGCAAGGTGCCGCTGCAGCAA
AACTTCCAAGACAACCAGTTTCAGGGTAAATGGTACATCGTTGGTTACGCGGGCAACGACTTCCTGCGTGAGGACAAGGAC
CCGTTCAAAATGTGGGCGCGTATTTATGAGCTGAAGGAAGACAAAAGCTACAACGTGACCGGTGTTTTCTTCCGTAAGAAG
AAATGCTACTATCATATTCATACCTTCGTGCCGGGTAGCCAACCGGGCGAATTTACCCTGGGTAACATTAAGAGCTACCCG
GGCTACACCAGCCAGCTGGTTCGTGTGGTTAGCACCAACTATAACCAGCACGCGATGGTGTTCACTAAGTACGTTTACCAA
AACCGTGAGATGTTCTACATTGTTCTGTATGGCCGTACCAAGGAACTGACCAGCGAGCTGAAAGAAAACTTCATCCGTTTT
AGCAAAAGCCTGGGTCTGCCGGAAAACCACATTGTGTTTCCGGTTCCGATCGACCAATGCATTGATGGCGCTAGCGGTGGT
AGTGGTGGTTCACCGGTCGCCACCATGGTAGCAGGTCATGCCTCTGGCAGCCCCGCATTCGGGACCGCCTCTCATTCGAAT
TGCGAACATGAAGAGATCCACCTCGCCGGCTCGATCCAGCCGCATGGCGCGCTTCTGGTCGTCAGCGAACATGATCATCGC
GTCATCCAGGCCAGCGCCAACGCCGCGGAATTTCTGAATCTCGGAAGCGTACTCGGCGTTCCGCTCGCCGAGATCGACGGC
GATCTGTTGATCAAGATCCTGCCGCATCTCGATCCCACCGCCGAAGGCATGCCGGTCGCGGTGCGCTGCCGGATCGGCAAT
CCCTCTACGGAGTACTGCGGTCTGATGCATCGGCCTCCGGAAGGCGGGCTGATCATCGAACTCGAACGTGCCGGCCCGTCG
ATCGATCTGTCAGGCACGCTGGCGCCGGCGCTGGAGCGGATCCGCACGGCGGGTTCACTGCGCGCGCTGTGCGATGACACC
GTGCTGCTGTTTCAGCAGTGCACCGGCTACGACCGGGTGATGGTGTATCGTTTCGATGAGCAAGGCCACGGCCTGGTATTC
TCCGAGTGCCATGTGCCTGGGCTCGAATCCTATTTCGGCAACCGCTATCCGTCGTCGACTGTCCCGCAGATGGCGCGGCAG
CTGTACGTGCGGCAGCGCGTCCGCGTGCTGGTCGACGTCACCTATCAGCCGGTGCCGCTGGAGCCGCGGCTGTCGCCGCTG
ACCGGGCGCGATCTCGACATGTCGGGCTGCTTCCTGCGCTCGATGTCGCCGTGCCATCTGCAGTTCCTGAAGGACATGGGC
GTGCGCGCCACCCTGGCGGTGTCGCTGGTGGTCGGCGGCAAGCTGTGGGGCCTGGTTGTCTGTCACCATTATCTGCCGCGC
TTCATCCGTTTCGAGCTGCGGGCGATCTGCAAACGGCTCGCCGAAAGGATCGCGACGCGGATCACCGCGCTTGAGAGCCTG
TACAAGGGCGGCAGCAACGAGCTGGCCCTGAAGCTGGCCGGCCTGGACATCAACAAGACCGGCGGTACCGGAAGCGGAGCT
ACTAACTTCAGCCTGCTGAAGCAGGCTGGAGACGTGGAGGAGAACCCTGGACCTCAATTGATGGCAGAAATCGGTACTGGC
TTTCCATTCGACCCCCATTATGTGGAAGTCCTGGGCGAGCGCATGCACTACGTCGATGTTGGTCCGCGCGATGGCACCCCT
GTGCTGTTCCTGCACGGTAACCCGACCTCCTCCTACGTGTGGCGCAACATCATCCCGCATGTTGCACCGACCCATCGCTGC
ATTGCTCCAGACCTGATCGGTATGGGCAAATCCGACAAACCAGACCTGGGTTATTTCTTCGACGACCACGTCCGCTTCATG
GATGCCTTCATCGAAGCCCTGGGTCTGGAAGAGGTCGTCCTGGTCATTCACGACTGGGGCTCCGCTCTGGGTTTCCACTGG
GCCAAGCGCAATCCAGAGCGCGTCAAAGGTATTGCATTTATGGAGTTCATCCGCCCTATCCCGACCTGGGACGAATGGCCA
GAATTTGCCCGCGAGACCTTCCAGGCCTTCCGCACCACCGACGTCGGCCGCAAGCTGATCATCGATCAGAACGTTTTTATC
GAGGGTACGCTGCCGATGGGTGTCGTCCGCCCGCTGACTGAAGTCGAGATGGACCATTACCGCGAGCCGTTCCTGAATCCT
GTTGACCGCGAGCCACTGTGGCGCTTCCCAAACGAGCTGCCAATCGCCGGTGAGCCAGCGAACATCGTCGCGCTGGTCGAA
GAATACATGGACTGGCTGCACCAGTCCCCTGTCCCGAAGCTGCTGTTCTGGGGCACCCCAGGCGTTCTGATCCCACCGGCC
GAAGCCGCTCGCCTGGCCAAAAGCCTGCCTAACTGCAAGGCTGTGGACATCGGCCCGGGTCTGAATCTGCTGCAAGAAGAC
AACCCGGACCTGATCGGCAGCGAGATCGCGCGCTGGCTGTCGACGCTCGAGATTTCCGGCGGCTCCGGTGCCAGTGCTGGT
GGTGGCAGCCTCGAGATTTCCGGCGGGAAGAAAAAGAAGAAGAAGTCCAAGACAAAATGCGTGATTATGTAG
>AzoTag16 miRFP670 NES P2A HaloTag KRas4B(CT)

**pCAGGS-HaloTag-miRFP670**
>Amino acid sequence
MAEIGTGFPFDPHYVEVLGERMHYVDVGPRDGTPVLFLHGNPTSSYVWRNIIPHVAPTHRCIAPDLIGMGKSDKPDLGYFF
DDHVRFMDAFIEALGLEEVVLVIHDWGSALGFHWAKRNPERVKGIAFMEFIRPIPTWDEWPEFARETFQAFRTTDVGRKLI
IDQNVFIEGTLPMGVVRPLTEVEMDHYREPFLNPVDREPLWRFPNELPIAGEPANIVALVEEYMDWLHQSPVPKLLFWGTP
GVLIPPAEAARLAKSLPNCKAVDIGPGLNLLQEDNPDLIGSEIARWLSTLEISGAAASAPPVATMVAGHASGSPAFGTASH
SNCEHEEIHLAGSIQPHGALLVVSEHDHRVIQASANAAEFLNLGSVLGVPLAEIDGDLLIKILPHLDPTAEGMPVAVRCRI
GNPSTEYCGLMHRPPEGGLIIELERAGPSIDLSGTLAPALERIRTAGSLRALCDDTVLLFQQCTGYDRVMVYRFDEQGHGL
VFSECHVPGLESYFGNRYPSSTVPQMARQLYVRQRVRVLVDVTYQPVPLEPRLSPLTGRDLDMSGCFLRSMSPCHLQFLKD
MGVRATLAVSLVVGGKLWGLVVCHHYLPRFIRFELRAICKRLAERIATRITALESSGSASREYPGYSRPPPRWSSSFCSL*
>DNA sequence
ATGGCAGAAATCGGTACTGGCTTTCCATTCGACCCCCATTATGTGGAAGTCCTGGGCGAGCGCATGCACTACGTCGATGTT
GGTCCGCGCGATGGCACCCCTGTGCTGTTCCTGCACGGTAACCCGACCTCCTCCTACGTGTGGCGCAACATCATCCCGCAT
GTTGCACCGACCCATCGCTGCATTGCTCCAGACCTGATCGGTATGGGCAAATCCGACAAACCAGACCTGGGTTATTTCTTC
GACGACCACGTCCGCTTCATGGATGCCTTCATCGAAGCCCTGGGTCTGGAAGAGGTCGTCCTGGTCATTCACGACTGGGGC
TCCGCTCTGGGTTTCCACTGGGCCAAGCGCAATCCAGAGCGCGTCAAAGGTATTGCATTTATGGAGTTCATCCGCCCTATC
CCGACCTGGGACGAATGGCCAGAATTTGCCCGCGAGACCTTCCAGGCCTTCCGCACCACCGACGTCGGCCGCAAGCTGATC
ATCGATCAGAACGTTTTTATCGAGGGTACGCTGCCGATGGGTGTCGTCCGCCCGCTGACTGAAGTCGAGATGGACCATTAC
CGCGAGCCGTTCCTGAATCCTGTTGACCGCGAGCCACTGTGGCGCTTCCCAAACGAGCTGCCAATCGCCGGTGAGCCAGCG
AACATCGTCGCGCTGGTCGAAGAATACATGGACTGGCTGCACCAGTCCCCTGTCCCGAAGCTGCTGTTCTGGGGCACCCCA

GGCGTTCTGATCCCACCGGCCGAAGCCGCTCGCCTGGCCAAAAGCCTGCCTAACTGCAAGGCTGTGGACATCGGCCCAGGT
CTGAATCTGCTGCAAGAAGACAACCCGGACCTGATCGGCAGCGAGATCGCGCGCTGGCTGTCCACGCTGGAGATTTCCGGC
GCGGCCGCTTCGGCTCCACCGGTCGCCACCATGGTAGCAGGTCATGCCTCTGGCAGCCCCGCATTCGGGACCGCCTCTCAT
TCGAATTGCGAACATGAAGAGATCCACCTCGCCGGCTCGATCCAGCCGCATGGCGCGCTTCTGGTCGTCAGCGAACATGAT
CATCGCGTCATCCAGGCCAGCGCCAACGCCGCGGAATTTCTGAATCTCGGAAGCGTACTCGGCGTTCCGCTCGCCGAGATC
GACGGCGATCTGTTGATCAAGATCCTGCCGCATCTCGATCCCACCGCCGAAGGCATGCCGGTCGCGGTGCGCTGCCGGATC
GGCAATCCCTCTACGGAGTACTGCGGTCTGATGCATCGGCCTCCGGAAGGCGGGCTGATCATCGAACTCGAACGTGCCGGC
CCGTCGATCGATCTGTCAGGCACGCTGGCGCCGGCGCTGGAGCGGATCCGCACGGCGGGTTCACTGCGCGCGCTGTGCGAT
GACACCGTGCTGCTGTTTCAGCAGTGCACCGGCTACGACCGGGTGATGGTGTATCGTTTCGATGAGCAAGGCCACGGCCTG
GTATTCTCCGAGTGCCATGTGCCTGGGCTCGAATCCTATTTCGGCAACCGCTATCCGTCGTCGACTGTCCCGCAGATGGCG
CGGCAGCTGTACGTGCGGCAGCGCGTCCGCGTGCTGGTCGACGTCACCTATCAGCCGGTGCCGCTGGAGCCGCGGCTGTCG
CCGCTGACCGGGCGCGATCTCGACATGTCGGGCTGCTTCCTGCGCTCGATGTCGCCGTGCCATCTGCAGTTCCTGAAGGAC
ATGGGCGTGCGCGCCACCCTGGCGGTGTCGCTGGTGGTCGGCGGCAAGCTGTGGGGCCTGGTTGTCTGTCACCATTATCTG
CCGCGCTTCATCCGTTTCGAGCTGCGGGCGATCTGCAAACGGCTCGCCGAAAGGATCGCGACGCGGATCACCGCGCTTGAG
AGCTCTGGGTCCGCATCTCGAGAGTACCCGGGATACTCGCGGCCGCCACCGCGGTGGAGCTCCAGCTTTTGTTCCCTTTAG
>HaloTag miRFP670

**pCAGGS-AzoTag16-miRFP680-cRaf**

>Amino acid sequence

MTNRSGSQDSTSDLIPAPPLSKVPLQQNFQDNQFQGKWYIVGYAGNDFLREDKDPFKMWARIYELKEDKSYNVTGVFFRKK
KCYYHIHTFVPGSQPGEFTLGNIKSYPGYTSQLVRVVSTNYNQHAMVFTKYVYQNREMFYIVLYGRTKELTSELKENFIRF
SKSLGLPENHIVFPVPIDQCIDGASLIKGAMAEGSVARQPDLLTCDDEPIHIPGAIQPHGLLLALAADMTIVAGSDNLPEL
TGLAIGALIGRSAADVFDSETHNRLTIALAEPGAAVGAPITVGFTMRKDAGFIGSWHRHDQLIFLELEPPQRDVAEPQAFF
RRTNSAIRRLQAAETLESACAAAAQEVRKITGFDRVMIYRFASDFSGEVIAEDRCAEVESKLGLHYPASTVPAQARRLYTI
NPVRIIPDINYRPVPVTPDLNPVTGRPIDLSFAILRSVSPCHLEFMRNIGMHGTMSISILRGERLWGLIVCHHRTPYYVDL
DGRQACKRVAERLATQIGVMEELEMAAAGAASNPGGGPEMVRGQAFDVGPRYILYKSGLRSRQGSGAGSGAGSGAGSGAGS
GAPRSLEMEHIQGAWKTISNGFGFKDAVFDGSSCISPTIVQQFGYQRRASDDGKLTDPSKTSNTIRVFLPNKQRTVVNVRN
GMSLHDCLMKALKVRGLQPECCAVFRLLHEHKGKKARLDWNTDAASLIGEELQVDFLDHVPLTTHNFARKTFLKLAFCDIC
QKFLLNGFRCQTCGYKFHEHCSTKVPTMCVDWSNIRQLLLFPNSTIGDSGVPALPSLTMRRMRESVSRMPVSSQHRYSTPH
AFTFNTSSPSSEGSLSQRQRSTSTPNVHMVSTTLPVDSRMIEDAIRSHSESASPSALSSSPNNLSPTGWSQPKTPVPAQRE
RAPVSGTQEKNKIRPRGQRDSSYYWEIEASEVMLSTRIGSGSFGTVYKGKWHGDVAVKILKVVDPTPEQFQAFRNEVAVLR
KTRHVNILLFMGYMTKDNLAIVTQWCEGSSLYKHLHVQETKFQMFQLIDIARQTAQGMDYLHAKNIIHRDMKSNNIFLHEG
LTVKIGDFGLATVKSRWSGSQQVEQPTGSVLWMAPEVIRMQDNNPFSFQSDVYSYGIVLYELMTGELPYSHINNRDQIIFM
VGRGYASPDLSKLYKNCPKAMKRLVADCVKKVKEERPLFPQILSSIELLQHSLPKINRSASEPSLHRAAHTEDINACTLTT
SPRLPVF*

>DNA sequence

ATGACAAATCGATCTGGTTCTCAAGATAGTACCAGCGATCTGATTCCGGCGCCGCCGCTGAGCAAGGTGCCGCTGCAGCAA
AACTTCCAAGACAACCAGTTTCAGGGTAAATGGTACATCGTTGGTTACGCGGGCAACGACTTCCTGCGTGAGGACAAGGAC
CCGTTCAAAATGTGGGCGCGTATTTATGAGCTGAAGGAAGACAAAAGCTACAACGTGACCGGTGTTTTCTTCCGTAAGAAG
AAATGCTACTATCATATTCATACCTTCGTGCCGGGTAGCCAACCGGGCGAATTTACCCTGGGTAACATTAAGAGCTACCCG
GGCTACACCAGCCAGCTGGTTCGTGTGGTTAGCACCAACTATAACCAGCACGCGATGGTGTTCACTAAGTACGTTTACCAA
AACCGTGAGATGTTCTACATTGTTCTGTATGGCCGTACCAAGGAACTGACCAGCGAGCTGAAAGAAAACTTCATCCGTTTT
AGCAAAAGCCTGGGTCTGCCGGAAAACCACATTGTGTTTCCGGTTCCGATCGACCAATGCATTGATGGCGCTAGCTTAATT
AAGGGCGCAATGGCGGAAGGCTCCGTCGCCAGGCAGCCTGACCTCTTGACCTGCGACGATGAGCCGATCCATATCCCCGGT
GCCATCCAACCGCATGGACTGCTGCTCGCCCTCGCCGCCGACATGACGATCGTTGCCGGCAGCGACAACCTTCCCGAACTC
ACCGGACTGGCGATCGGCGCCCTGATCGGCCGCTCTGCGGCCGATGTCTTCGACTCGGAGACGCACAACCGTCTGACGATC
GCCTTGGCCGAGCCCGGGGCGGCCGTCGGAGCACCGATCACTGTCGGCTTCACGATGCGAAAGGACGCAGGCTTCATCGGC
TCCTGGCATCGCCATGATCAGCTCATCTTCCTCGAGCTCGAGCCTCCCCAGCGGGACGTCGCCGAGCCGCAGGCGTTCTTC
CGCCGCACCAACAGCGCCATCCGCCGCCTGCAGGCCGCCGAAACCTTGGAAAGCGCCTGCGCCGCCGCGGCGCAAGAGGTG
CGGAAGATTACCGGCTTCGATCGGGTGATGATCTATCGCTTCGCCTCCGACTTCAGCGGCGAAGTGATCGCAGAGGATCGG
TGCGCCGAGGTCGAGTCAAAACTAGGCCTGCACTATCCTGCCTCAACCGTGCCGGCGCAGGCCCGTCGGCTCTATACCATC
AACCCGGTACGGATCATTCCCGATATCAATTATCGGCCGGTGCCGGTCACCCCAGACCTCAATCCGGTCACCGGGCGGCCG
ATTGATCTTAGCTTCGCCATCCTGCGCAGCGTCTCGCCCTGCCATCTGGAGTTCATGCGCAACATAGGCATGCACGGCACG
ATGTCGATCTCGATTTTGCGCGGCGAGCGACTGTGGGGATTGATCGTTTGCCATCACCGAACGCCGTACTACGTCGATCTC
GATGGCCGCCAAGCCTGCAAGAGAGTGGCCGAGAGGCTGGCCACCCAGATCGGCGTGATGGAAGAGCTCGAGATGGCAGCG
GCAGGAGCTGCGTCTAACCCCGGCGGGGGTCCGGAGATGGTGCGGGGCCAGGCGTTCGACGTAGGCCCTCGATACATCCTG
TACAAGTCCGGACTCAGATCTCGACAAGGTAGTGGTGCTGGCTCTGGTGCTGGTAGTGGCGCTGGTTCCGGTGCTGGCTCT
GGCGCGCCTCGAAGCCTCGAGATGGAGCACATACAGGGAGCTTGGAAGACGATCAGCAATGGTTTTGGATTCAAAGATGCC
GTGTTTGATGGCTCCAGCTGCATCTCTCCTACAATAGTTCAGCAGTTTGGCTATCAGCGCCGGGCATCAGATGATGGCAAA
CTCACAGATCCTTCTAAGACAAGCAACACTATCCGTGTTTTCTTGCCGAACAAGCAAAGAACAGTGGTCAATGTGCGAAAT
GGAATGAGCTTGCATGACTGCCTTATGAAAGCACTCAAGGTGAGGGGCCTGCAACCAGAGTGCTGTGCAGTGTTCAGACTT
CTCCACGAACACAAAGGTAAAAAAGCACGCTTAGATTGGAATACTGATGCTGCGTCTTTGATTGGAGAAGAACTTCAAGTA
GATTTCCTGGATCATGTTCCCCTCACAACACACAACTTTGCTCGGAAGACGTTCCTGAAGCTTGCCTTCTGTGACATCTGT
CAGAAATTCCTGCTCAATGGATTTCGATGTCAGACTTGTGGCTACAAATTTCATGAGCACTGTAGCACCAAAGTACCTACT
ATGTGTGTGGACTGGAGTAACATCAGACAACTCTTATTGTTTCCAAATTCCACTATTGGTGATAGTGGAGTCCCAGCACTA
CCTTCTTTGACTATGCGTCGTATGCGAGAGTCTGTTTCCAGGATGCCTGTTAGTTCTCAGCACAGATATTCTACACCTCAC
GCCTTCACCTTTAACACCTCCAGTCCCTCATCTGAAGGTTCCCTCTCCCAGAGGCAGAGGTCGACATCCACACCTAATGTC
CACATGGTCAGCACCACGCTGCCTGTGGACAGCAGGATGATTGAGGATGCAATTCGAAGTCACAGCGAATCAGCCTCACCT

TCAGCCCTGTCCAGTAGCCCCAACAATCTGAGCCCAACAGGCTGGTCACAGCCGAAAACCCCCGTGCCAGCACAAAGAGAG
CGGGCACCAGTATCTGGGACCCAGGAGAAAAACAAAATTAGGCCTCGTGGACAGAGAGATTCAAGCTATTATTGGGAAATA
GAAGCCAGTGAAGTGATGCTGTCCACTCGGATTGGGTCAGGCTCTTTTGGAACTGTTTATAAGGGTAAATGGCACGGAGAT
GTTGCAGTAAAGATCCTAAAGGTTGTCGACCCAACCCCAGAGCAATTCCAGGCCTTCAGGAATGAGGTGGCTGTTCTGCGC
AAAACACGGCATGTGAACATTCTGCTTTTCATGGGGTACATGACAAAGGACAACCTGGCAATTGTGACCCAGTGGTGCGAG
GGCAGCAGCCTCTACAAACACCTGCATGTCCAGGAGACCAAGTTTCAGATGTTCCAGCTAATTGACATTGCCCGGCAGACG
GCTCAGGGAATGGACTATTTGCATGCAAAGAACATCATCCATAGAGACATGAAATCCAACAATATATTTCTCCATGAAGGC
TTAACAGTGAAAATTGGAGATTTTGGTTTGGCAACAGTAAAGTCACGCTGGAGTGGTTCTCAGCAGGTTGAACAACCTACT
GGCTCTGTCCTCTGGATGGCCCCAGAGGTGATCCGAATGCAGGATAACAACCCATTCAGTTTCCAGTCGGATGTCTACTCC
TATGGCATCGTATTGTATGAACTGATGACGGGGGAGCTTCCTTATTCTCACATCAACAACCGAGATCAGATCATCTTCATG
GTGGGCCGAGGATATGCCTCCCCAGATCTTAGTAAGCTATATAAGAACTGCCCCAAAGCAATGAAGAGGCTGGTAGCTGAC
TGTGTGAAGAAAGTAAAGGAAGAGAGGCCTCTTTTTCCCCAGATCCTGTCTTCCATTGAGCTGCTCCAACACTCTCTACCG
AAGATCAACCGGAGCGCTTCCGAGCCATCCTTGCATCGGGCAGCCCACACTGAGGATATCAATGCTTGCACGCTGACCACG
TCCCCGAGGCTGCCTGTCTTCTAG
>AzoTag16 miRFP680 cRaf

**pCAGGS-MEK1-P2A-mScarlet-ERK2(K57R)**
>Amino acid sequence
MPKKKPTPIQLNPNPEGTAVNGTPTAETNLEALQKKLEELELDEQQRKRLEAFLTQKQKVGELKDDDFEKVSELGAGNGGV
VFKVSHKPTSLIMARKLIHLEIKPAIRNQIIRELQVLHECNSPYIVGFYGAFYSDGEISICMEHMDGGSLDQVLKKAGKIP
EKILGKVSIAVIKGLTYLREKHKIMHRDVKPSNILVNSRGEIKLCDFGVSGQLIDSMANSFVGTRSYMSPERLQGTHYSVQ
SDIWSMGLSLVEMAIGRYPIPPPDAKELELIFGCSVERDPASSELAPRPRPPGRPISSYGPDSRPPMAIFELLDYIVNEPP
PKLPSGVFGAEFQDFVNKCLVKNPAERADLKQLMVHSFIKQSELEEVDFAGWLCSTMGLKQPSTPTHAAGVGGRGTSGSGA
TNFSLLKQAGDVEENPGPQLILPVTMVSKGEAVIKEFMRFKVHMEGSMNGHEFEIEGEGEGRPYEGTQTAKLKVTKGGPLP
FSWDILSPQFMYGSRAFIKHPADIPDYYKQSFPEGFKWERVMNFEDGGAVTVTQDTSLEDGTLIYKVKLRGTNFPPDGPVM
QKKTMGWEASTERLYPEDGVLKGDIKMALRLKDGGRYLADFKTTYKAKKPVQMPGAYNVDRKLDITSHNEDYTVVEQYERS
EGRHSTGGMDELYKSTVPRKLDITSHNEDYTIVEQYDRAEGRHSTGGMDELYLEMAAAGAASNPGGGPEMVRGQAFDVGPR
YINLAYIGEGAYGMVCSAHDNVNKVRVAIRKISPFEHQTYCQRTLREIKILLRFKHENIIGINDIIRAPTIEQMKDVYIVQ
DLMETDLYKLLKTQHLSNDHICYFLYQILRGLKYIHSANVLHRDLKPSNLLLNTTCDLKICDFGLARVADPDHDHTGFLTE
YVATRWYRAPEIMLNSKGYTKSIDIWSVGCILAEMLSNRPIFPGKHYLDQLNHILGILGSPSQEDLNCIINLKARNYLLSL
PHKNKVPWNRLFPNADPKALDLLDKMLTFNPHKRIEVEAALAHPYLEQYYDPSDEPVAEAPFKFEMELDDLPKETLKELIF
EETARFQPGY*
>DNA sequence
ATGCCTAAAAAGAAGCCTACGCCCATACAGCTGAATCCCAACCCCGAAGGGACTGCTGTGAACGGGACCCCTACAGCCGAG
ACAAACCTTGAAGCTCTGCAGAAAAAGTTGGAAGAGCTTGAGCTGGATGAGCAGCAGAGGAAGCGTCTGGAGGCTTTTCTC
ACCCAGAAGCAGAAAGTTGGGGAACTGAAGGATGACGACTTTGAAAAAGTTTCAGAGCTTGGAGCAGGCAACGGAGGAGTG
GTGTTTAAGGTGTCCCACAAGCCAACCAGCTTGATTATGGCCAGGAAGTTGATTCATCTGGAGATTAAGCCTGCAATCCGA
AACCAGATTATCCGAGAGTTGCAGGTTCTGCATGAATGTAACTCCCCATACATTGTGGGGTTCTATGGGGCCTTCTACAGT
GATGGAGAGATCAGCATTTGCATGGAACACATGGATGGAGGCTCCCTTGATCAGGTTCTGAAGAAAGCTGGCAAAATCCCA
GAAAAGATTTTGGGAAAAGTCAGCATTGCAGTGATAAAAGGTCTAACCTACCTGAGAGAAAAGCATAAGATAATGCACAGA
GATGTGAAACCTTCTAACATCCTGGTCAACTCTAGAGGAGAGATAAAACTCTGCGACTTTGGGGTCAGCGGGCAACTCATA
GACTCCATGGCAAATTCCTTTGTTGGGACAAGATCCTATATGTCACCGGAGCGACTACAGGGCACTCATTATTCTGTGCAA
TCAGACATCTGGAGCATGGGGCTGTCGCTGGTGGAAATGGCCATTGGAAGGTATCCCATTCCACCCCCTGATGCCAAAGAG
CTGGAACTTATCTTTGGGTGTTCTGTAGAAAGGGATCCAGCGTCTTCTGAACTGGCACCTCGCCCCCGGCCACCCGGACGT
CCAATAAGCTCATACGGTCCTGATAGTCGACCACCCATGGCTATTTTTGAACTTCTGGATTATATCGTGAACGAGCCGCCT
CCAAAATTGCCCAGTGGAGTATTTGGAGCTGAGTTCCAGGACTTTGTGAATAAATGTCTTGTGAAGAATCCGGCAGAGAGA
GCAGACCTTAAACAGCTAATGGTTCACAGCTTCATTAAGCAGTCAGAGTTGGAGGAAGTGGATTTTGCTGGATGGCTCTGT
TCCACTATGGGCCTTAAGCAGCCCAGTACCCCAACCCATGCCGCCGGAGTGGGCGGCCGCGGCACTAGTGGAAGCGGAGCT
ACTAACTTCAGCCTGCTGAAGCAGGCTGGAGACGTGGAGGAGAACCCTGGACCTCAATTAATTCTACCGGTCACCATGGTG
AGCAAGGGCGAGGCAGTGATCAAGGAGTTCATGCGGTTCAAGGTGCACATGGAGGGCTCCATGAACGGCCACGAGTTCGAG
ATCGAGGGCGAGGGCGAGGGCCGCCCCTACGAGGGCACCCAGACCGCCAAGCTGAAGGTGACCAAGGGTGGCCCCCTGCCC
TTCTCCTGGGACATCCTGTCCCCTCAGTTCATGTACGGCTCCAGGGCCTTCATCAAGCACCCCGCCGACATCCCCGACTAC
TATAAGCAGTCCTTCCCCGAGGGCTTCAAGTGGGAGCGCGTGATGAACTTCGAGGACGGCGGCGCCGTGACCGTGACCCAG
GACACCTCCCTGGAGGACGGCACCCTGATCTACAAGGTGAAGCTCCGCGGCACCAACTTCCCTCCTGACGGCCCCGTAATG
CAGAAGAAGACAATGGGCTGGGAAGCATCCACCGAGCGGTTGTACCCCGAGGACGGCGTGCTGAAGGGCGACATTAAGATG
GCCCTGCGCCTGAAGGACGGCGGCCGCTACCTGGCGGACTTCAAGACCACCTACAAGGCCAAGAAGCCCGTGCAGATGCCC
GGCGCCTACAACGTCGACCGCAAGTTGGACATCACCTCCCACAACGAGGACTACACCGTGGTGGAACAGTACGAACGCTCC
GAGGGCCGCCACTCCACCGGCGGCATGGACGAGCTGTACAAGTCGACGGTACCGCGGAAGTTGGACATCACCTCCCACAAC
GAGGACTACACCATCGTGGAACAGTACGACCGCGCCGAGGGCCGCCACTCCACCGGCGGCATGGACGAGCTGTACCTCGAG
ATGGCAGCGGCAGGAGCTGCGTCTAACCCCGGCGGGGGTCCGGAGATGGTGCGGGGCCAGGCGTTCGACGTAGGCCCTCGA
TACATCAATCTGGCTTATATCGGCGAGGGAGCGTACGGCATGGTGTGTTCTGCCCATGACAATGTTAACAAAGTTCGAGTT
GCTATCAGGAAAATCAGCCCATTTGAGCATCAGACATACTGCCAGCGAACATTGCGGGAGATCAAAATCTTGCTACGTTTT
AAACATGAAAACATCATTGGGATAAACGACATTATTCGCGCTCCAACCATTGAGCAGATGAAAGATGTGTACATTGTGCAG
GACCTCATGGAGACAGACCTCTATAAGCTCCTGAAGACTCAGCATCTTAGCAATGACCATATCTGCTATTTCTTGTACCAG
ATTCTGAGAGGATTAAAGTACATCCATTCAGCCAATGTTCTACATCGTGATCTTAAGCCTTCAAATTTGCTGCTTAACACT
ACCTGTGATCTCAAGATCTGTGATTTTGGATTGGCTCGTGTTGCAGACCCAGATCATGATCACACTGGCTTTCTCACAGAA
TATGTAGCCACTCGCTGGTACAGAGCTCCTGAGATCATGCTGAATTCCAAGGGCTATACCAAATCAATTGACATCTGGTCT
GTTGGCTGCATTCTTGCTGAGATGCTTTCTAATAGACCCATATTTCCTGGGAAACATTATCTTGACCAGCTTAATCACATA

CTTGGTATTCTTGGATCTCCATCTCAAGAGGACCTAAACTGTATAATCAATTTAAAAGCTAGGAATTACTTGCTTTCCCTT
CCTCACAAAAATAAGGTGCCATGGAACAGACTTTTCCCCAATGCAGATCCCAAAGCTCTAGACTTACTGGACAAGATGCTG
ACTTTCAACCCCCATAAAAGAATTGAAGTAGAGGCAGCTTTGGCTCATCCTTATCTGGAGCAGTATTATGACCCAAGTGAT
GAGCCTGTAGCTGAAGCTCCCTTTAAATTTGAAATGGAGCTTGATGATTTGCCCAAGGAGACTCTTAAGGAGCTAATTTTT
GAAGAAACCGCTAGATTCCAGCCAGGGTACTAA
>MEK1 P2A mScarlet-I ERK2(K57R)

**pCAGGS-AzoTag16-miRFP680-p85$_{iSH2}$**

>Amino acid sequence
MTNRSGSQDSTSDLIPAPPLSKVPLQQNFQDNQFQGKWYIVGYAGNDFLREDKDPFKMWARIYELKEDKSYNVTGVFFRKK
KCYYHIHTFVPGSQPGEFTLGNIKSYPGYTSQLVRVVSTNYNQHAMVFTKYVYQNREMFYIVLYGRTKELTSELKENFIRF
SKSLGLPENHIVFPVPIDQCIDGASLIKGAMAEGSVARQPDLLTCDDEPIHIPGAIQPHGLLLALAADMTIVAGSDNLPEL
TGLAIGALIGRSAADVFDSETHNRLTIALAEPGAAVGAPITVGFTMRKDAGFIGSWHRHDQLIFLELEPPQRDVAEPQAFF
RRTNSAIRRLQAAETLESACAAAAQEVRKITGFDRVMIYRFASDFSGEVIAEDRCAEVESKLGLHYPASTVPAQARRLYTI
NPVRIIPDINYRPVPVTPDLNPVTGRPIDLSFAILRSVSPCHLEFMRNIGMHGTMSISILRGERLWGLIVCHHRTPYYVDL
DGRQACKRVAERLATQIGVMEELEMAAAGAASNPGGGPEMVRGQAFDVGPRYILYKSGLRSRQGSGAGSGAGSGAGSGAGS
GAPRASKYQQDQVVKEDSVEAVGAQLKVYHQQYQDKSREYDQLYEEYTRTSQELQMKRTAIEAFNETIKIFEEQGQTQEKC
SKEYLERFRREGNEKEMQRILLNSERLKSRIAEIHESRTKLEQDLRAQASDNREIDKRMNSLKPDLMQLRKIRDQYLVWLT
QKGARQRKINEWLGIKNETEDQYSLMEDEDALPHHEERT*
>DNA sequence
ATGACAAATCGATCTGGTTCTCAAGATAGTACCAGCGATCTGATTCCGGCGCCGCCGCTGAGCAAGGTGCCGCTGCAGCAA
AACTTCCAAGACAACCAGTTTCAGGGTAAATGGTACATCGTTGGTTACGCGGGCAACGACTTCCTGCGTGAGGACAAGGAC
CCGTTCAAAATGTGGGCGCGTATTTATGAGCTGAAGGAAGACAAAAGCTACAACGTGACCGGTGTTTTCTTCCGTAAGAAG
AAATGCTACTATCATATTCATACCTTCGTGCCGGGTAGCCAACCGGGCGAATTTACCCTGGGTAACATTAAGAGCTACCCG
GGCTACACCAGCCAGCTGGTTCGTGTGGTTAGCACCAACTATAACCAGCACGCGATGGTGTTCACTAAGTACGTTTACCAA
AACCGTGAGATGTTCTACATTGTTCTGTATGGCCGTACCAAGGAACTGACCAGCGAGCTGAAAGAAAACTTCATCCGTTTT
AGCAAAAGCCTGGGTCTGCCGGAAAACCACATTGTGTTTCCGGTTCCGATCGACCAATGCATTGATGGCGCTAGCTTAATT
AAGGGCGCAATGGCGGAAGGCTCCGTCGCCAGGCAGCCTGACCTCTTGACCTGCGACGATGAGCCGATCCATATCCCCGGT
GCCATCCAACCGCATGGACTGCTGCTCGCCCTCGCCGCCGACATGACGATCGTTGCCGGCAGCGACAACCTTCCCGAACTC
ACCGGACTGGCGATCGGCGCCCTGATCGGCCGCTCTGCGGCCGATGTCTTCGACTCGGAGACGCACAACCGTCTGACGATC
GCCTTGGCCGAGCCCGGGGCGGCCGTCGGAGCACCGATCACTGTCGGCTTCACGATGCGAAAGGACGCAGGCTTCATCGGC
TCCTGGCATCGCCATGATCAGCTCATCTTCCTCGAGCTCGAGCCTCCCCAGCGGGACGTCGCCGAGCCGCAGGCGTTCTTC
CGCCGCACCAACAGCGCCATCCGCCGCCTGCAGGCCGCCGAAACCTTGGAAAGCGCCTGCGCCGCCGCGGCGCAAGAGGTG
CGGAAGATTACCGGCTTCGATCGGGTGATGATCTATCGCTTCGCCTCCGACTTCAGCGGCGAAGTGATCGCAGAGGATCGG
TGCGCCGAGGTCGAGTCAAAACTAGGCCTGCACTATCCTGCCTCAACCGTGCCGGCGCAGGCCCGTCGGCTCTATACCATC
AACCCGGTACGGATCATTCCCGATATCAATTATCGGCCGGTGCCGGTCACCCCAGACCTCAATCCGGTCACCGGGCGGCCG
ATTGATCTTAGCTTCGCCATCCTGCGCAGCGTCTCGCCCTGCCATCTGGAGTTCATGCGCAACATAGGCATGCACGGCACG
ATGTCGATCTCGATTTTGCGCGGCGAGCGACTGTGGGGATTGATCGTTTGCCATCACCGAACGCCGTACTACGTCGATCTC
GATGGCCGCCAAGCCTGCAAGAGAGTGGCCGAGAGGCTGGCCACCCAGATCGGCGTGATGGAAGAGCTCGAGATGGCAGCG
GCAGGAGCTGCGTCTAACCCCGGCGGGGGTCCGGAGATGGTGCGGGGCCAGGCGTTCGACGTAGGCCCTCGATACATCCTG
TACAAGTCCGGACTCAGATCTCGACAAGGTAGTGGTGCTGGCTCTGGTGCTGGTAGTGGCGCTGGTTCCGGTGCTGGCTCT
GGCGCGCCTCGAGCATCCAAGTACCAACAAGACCAGGTGGTGAAGGAGGACAGCGTAGAGGCTGTGGGCGCCCAGCTCAAG
GTCTACCACCAGCAGTACCAGGACAAGAGCCGCGAATATGACCAGCTGTATGAAGAATACACACGGACCTCCCAGGAGCTG
CAGATGAAGCGCACAGCCATAGAGGCCTTCAACGAGACCATCAAGATCTTCGAAGAGCAGGGCCAGACACAGGAGAAGTGC
AGCAAGGAGTATTTGGAGCGCTTCCGGCGAGAGGGAAATGAGAAGGAGATGCAGAGGATCCTGCTGAACTCCGAGCGACTC
AAGTCTCGCATCGCGGAGATACACGAAAGCCGCACGAAGTTGGAGCAGGATCTGCGGGCGCAGGCCTCCGACAACCGTGAG
ATCGACAAGCGCATGAACAGCCTCAAACCTGACCTCATGCAGCTGCGCAAGATCAGGGACCAGTACCTCGTGTGGCTCACC
CAGAAAGGTGCCCGACAGAGGAAGATCAACGAATGGCTGGGAATCAAGAACGAGACTGAGGACCAGTATTCACTGATGGAG
GATGAGGACGCCCTCCCCCACCACGAGGAGCGCACGTGA
>AzoTag16 miRFP680 p85$_{iSH2}$(the inter-SH2 domain of the p85 subunit of PI3K)

**pCMV-mScarlet-PH$_{Akt}$**

>Amino acid sequence
MVSKGEAVIKEFMRFKVHMEGSMNGHEFEIEGEGEGRPYEGTQTAKLKVTKGGPLPFSWDILSPQFMYGSRAFIKHPADIP
DYYKQSFPEGFKWERVMNFEDGGAVTVTQDTSLEDGTLIYKVKLRGTNFPPDGPVMQKKTMGWEASTERLYPEDGVLKGDI
KMALRLKDGGRYLADFKTTYKAKKPVQMPGAYNVDRKLDITSHNEDYTVVEQYERSEGRHSTGGMDELYKDGTAGPGSMSD
VAIVKEGWLHKRGEYIKTWRPRYFLLKNDGTFIGYKERPQDVDQREAPLNNFSVAQCQLMKTERPRPNTFIIRCLQWTTVI
ERTFHVETPEEREEWTTAIQTVADGLKKQEEEEMDFRSGSPSDNSGAEEMEVSLAKPKHRVTMN*
>DNA sequence
ATGGTGAGCAAGGGCGAGGCAGTGATCAAGGAGTTCATGCGGTTCAAGGTGCACATGGAGGGCTCCATGAACGGCCACGAG
TTCGAGATCGAGGGCGAGGGCGAGGGCCGCCCCTACGAGGGCACCCAGACCGCCAAGCTGAAGGTGACCAAGGGTGGCCCC
CTGCCCTTCTCCTGGGACATCCTGTCCCCTCAGTTCATGTACGGCTCCAGGGCCTTCATCAAGCACCCCGCCGACATCCCC
GACTACTATAAGCAGTCCTTCCCCGAGGGCTTCAAGTGGGAGCGCGTGATGAACTTCGAGGACGGCGGCGCCGTGACCGTG
ACCCAGGACACCTCCCTGGAGGACGGCACCCTGATCTACAAGGTGAAGCTCCGCGGCACCAACTTCCCTCCTGACGGCCCC
GTAATGCAGAAGAAGACAATGGGCTGGGAAGCATCCACCGAGCGGTTGTACCCCGAGGACGGCGTGCTGAAGGGCGACATT
AAGATGGCCCTGCGCCTGAAGGACGGCGGCCGCTACCTGGCGGACTTCAAGACCACCTACAAGGCCAAGAAGCCCGTGCAG
ATGCCCGGCGCCTACAACGTCGACCGCAAGTTGGACATCACCTCCCACAACGAGGACTACACCGTGGTGGAACAGTACGAA

CGCTCCGAGGGCCGCCACTCCACCGGCGGCATGGACGAGCTGTACAAGDGTAGPGSATGAGCGACGTGGCTATTGTGAAGG
AGGGTTGGCTGCACAAACGAGGGGAGTACATCAAGACCTGGCGGCCACGCTACTTCCTCCTCAAGAATGATGGCACCTTCA
TTGGCTACAAGGAGCGGCCGCAGGATGTGGACCAACGTGAGGCTCCCCTCAACAACTTCTCTGTGGCGCAGTGCCAGCTGA
TGAAGACGGAGCGGCCCCGGCCCAACACCTTCATCATCCGCTGCCTGCAGTGGACCACTGTCATCGAACGCACCTTCCATG
TGGAGACTCCTGAGGAGCGGGAGGAGTGGACAACCGCCATCCAGACTGTGGCTGACGGCCTCAAGAAGCAGGAGGAGGAGG
AGATGGACTTCCGGTCGGGCTCACCCAGTGACAACTCAGGGGCTGAAGAGATGGAGGTGTCCCTGGCCAAGCCCAAGCACC
GCGTGACCATGAACTAA
>mScarlet-I $PH_{Akt}$ (the PH domain of Akt)

**pCMV-Lifeact-mScarlet**
>Amino acid sequence
MGVADLIKKFESISKEEGDPPVTMVSKGEAVIKEFMRFKVHMEGSMNGHEFEIEGEGEGRPYEGTQTAKLKVTKGGPLPFS
WDILSPQFMYGSRAFIKHPADIPDYYKQSFPEGFKWERVMNFEDGGAVTVTQDTSLEDGTLIYKVKLRGTNFPPDGPVMQK
KTMGWEASTERLYPEDGVLKGDIKMALRLKDGGRYLADFKTTYKAKKPVQMPGAYNVDRKLDITSHNEDYTVVEQYERSEG
RHSTGGMDELYK*
>DNA sequence
ATGGGCGTGGCCGACTTGATCAAGAAGTTCGAGTCCATCTCCAAGGAGGAGGGGGATCCACCGGTCACCATGGTGAGCAAG
GGCGAGGCAGTGATCAAGGAGTTCATGCGGTTCAAGGTGCACATGGAGGGCTCCATGAACGGCCACGAGTTCGAGATCGAG
GGCGAGGGCGAGGGCCGCCCCTACGAGGGCACCCAGACCGCCAAGCTGAAGGTGACCAAGGGTGGCCCCCTGCCCTTCTCC
TGGGACATCCTGTCCCCTCAGTTCATGTACGGCTCCAGGGCCTTCATCAAGCACCCCGCCGACATCCCCGACTACTATAAG
CAGTCCTTCCCCGAGGGCTTCAAGTGGGAGCGCGTGATGAACTTCGAGGACGGCGGCGCCGTGACCGTGACCCAGGACACC
TCCCTGGAGGACGGCACCCTGATCTACAAGGTGAAGCTCCGCGGCACCAACTTCCCTCCTGACGGCCCCGTAATGCAGAAG
AAGACAATGGGCTGGGAAGCATCCACCGAGCGGTTGTACCCCGAGGACGGCGTGCTGAAGGGCGACATTAAGATGGCCCTG
CGCCTGAAGGACGGCGGCCGCTACCTGGCGGACTTCAAGACCACCTACAAGGCCAAGAAGCCCGTGCAGATGCCCGGCGCC
TACAACGTCGACCGCAAGTTGGACATCACCTCCCACAACGAGGACTACACCGTGGTGGAACAGTACGAACGCTCCGAGGGC
CGCCACTCCACCGGCGGCATGGACGAGCTGTACAAGTAG
>Lifeact mScarlet-I

**pCMV-dCas9-AzoTag16**
>Amino acid sequence
MSPKKKRKVEASDKKYSIGLAIGTNSVGWAVITDEYKVPSKKFKVLGNTDRHSIKKNLIGALLFDSGETAEATRLKRTARR
RYTRRKNRICYLQEIFSNEMAKVDDSFFHRLEESFLVEEDKKHERHPIFGNIVDEVAYHEKYPTIYHLRKKLVDSTDKADL
RLIYLALAHMIKFRGHFLIEGDLNPDNSDVDKLFIQLVQTYNQLFEENPINASGVDAKAILSARLSKSRRLENLIAQLPGE
KKNGLFGNLIALSLGLTPNFKSNFDLAEDAKLQLSKDTYDDDLDNLLAQIGDQYADLFLAAKNLSDAILLSDILRVNTEIT
KAPLSASMIKRYDEHHQDLTLLKALVRQQLPEKYKEIFFDQSKNGYAGYIDGGASQEEFYKFIKPILEKMDGTEELLVKLN
REDLLRKQRTFDNGSIPHQIHLGELHAILRRQEDFYPFLKDNREKIEKILTFRIPYYVGPLARGNSRFAWMTRKSEETITP
WNFEEVVDKGASAQSFIERMTNFDKNLPNEKVLPKHSLLYEYFTVYNELTKVKYVTEGMRKPAFLSGEQKKAIVDLLFKTN
RKVTVKQLKEDYFKKIECFDSVEISGVEDRFNASLGTYHDLLKIIKDKDFLDNEENEDILEDIVLTLTLFEDREMIEERLK
TYAHLFDDKVMKQLKRRRYTGWGRLSRKLINGIRDKQSGKTILDFLKSDGFANRNFMQLIHDDSLTFKEDIQKAQVSGQGD
SLHEHIANLAGSPAIKKGILQTVKVVDELVKVMGRHKPENIVIEMARENQTTQKGQKNSRERMKRIEEGIKELGSQILKEH
PVENTQLQNEKLYLYYLQNGRDMYVDQELDINRLSDYDVDAIVPQSFLKDDSIDNKVLTRSDKNRGKSDNVPSEEVVKKMK
NYWRQLLNAKLITQRKFDNLTKAERGGLSELDKAGFIKRQLVETRQITKHVAQILDSRMNTKYDENDKLIREVKVITLKSK
LVSDFRKDFQFYKVREINNYHHAHDAYLNAVVGTALIKKYPKLESEFVYGDYKVYDVRKMIAKSEQEIGKATAKYFFYSNI
MNFFKTEITLANGEIRKRPLIETNGETGEIVWDKGRDFATVRKVLSMPQVNIVKKTEVQTGGFSKESILPKRNSDKLIARK
KDWDPKKYGGFDSPTVAYSVLVVAKVEKGKSKKLKSVKELLGITIMERSSFEKNPIDFLEAKGYKEVKKDLIIKLPKYSLF
ELENGRKRMLASAGELQKGNELALPSKYVNFLYLASHYEKLKGSPEDNEQKQLFVEQHKHYLDEIIEQISEFSKRVILADA
NLDKVLSAYNKHRDKPIREQAENIIHLFTLTNLGAPAAFKYFDTTIDRKRYTSTKEVLDATLIHQSITGLYETRIDLSQLG
GDSAGGGGSGGGGSGGGGSGPKKKRKVAAAGSMTNRSGSQDSTSDLIPAPPLSKVPLQQNFQDNQFQGKWYIVGYAGNDFL
REDKDPFKMWARIYELKEDKSYNVTGVFFRKKKCYYHIHTFVPGSQPGEFTLGNIKSYPGYTSQLVRVVSTNYNQHAMVFT
KYVYQNREMFYIVLYGRTKELTSELKENFIRFSKSLGLPENHIVFPVPIDQCIDG*
>DNA sequence
ATGAGCCCCAAGAAGAAGAGAAAGGTGGAGGCCAGCGACAAGAAGTACAGCATCGGCCTGGCCATCGGCACCAACTCTGTG
GGCTGGGCCGTGATCACCGACGAGTACAAGGTGCCCAGCAAGAAATTCAAGGTGCTGGGCAACACCGACCGGCACAGCATC
AAGAAGAACCTGATCGGAGCCCTGCTGTTCGACAGCGGCGAAACAGCCGAGGCCACCCGGCTGAAGAGAACCGCCAGAAGA
AGATACACCAGACGGAAGAACCGGATCTGCTATCTGCAAGAGATCTTCAGCAACGAGATGGCCAAGGTGGACGACAGCTTC
TTCCACAGACTGGAAGAGTCCTTCCTGGTGGAAGAGGATAAGAAGCACGAGCGGCACCCCATCTTCGGCAACATCGTGGAC
GAGGTGGCCTACCACGAGAAGTACCCCACCATCTACCACCTGAGAAAGAAACTGGTGGACAGCACCGACAAGGCCGACCTG
CGGCTGATCTATCTGGCCCTGGCCCACATGATCAAGTTCCGGGGCCACTTCCTGATCGAGGGCGACCTGAACCCCGACAAC
AGCGACGTGGACAAGCTGTTCATCCAGCTGGTGCAGACCTACAACCAGCTGTTCGAGGAAAACCCCATCAACGCCAGCGGC
GTGGACGCCAAGGCCATCCTGTCTGCCAGACTGAGCAAGAGCAGACGGCTGGAAAATCTGATCGCCCAGCTGCCCGGCGAG
AAGAAGAATGGCCTGTTCGGCAACCTGATTGCCCTGAGCCTGGGCCTGACCCCCAACTTCAAGAGCAACTTCGACCTGGCC
GAGGATGCCAAACTGCAGCTGAGCAAGGACACCTACGACGACGACCTGGACAACCTGCTGGCCCAGATCGGCGACCAGTAC
GCCGACCTGTTTCTGGCCGCCAAGAACCTGTCCGACGCCATCCTGCTGAGCGACATCCTGAGAGTGAACACCGAGATCACC
AAGGCCCCCCTGAGCGCCTCTATGATCAAGAGATACGACGAGCACCACCAGGACCTGACCCTGCTGAAAGCTCTCGTGCGG
CAGCAGCTGCCTGAGAAGTACAAAGAGATTTTCTTCGACCAGAGCAAGAACGGCTACGCCGGCTACATTGACGGCGGAGCC
AGCCAGGAAGAGTTCTACAAGTTCATCAAGCCCATCCTGGAAAAGATGGACGGCACCGAGGAACTGCTCGTGAAGCTGAAC
AGAGAGGACCTGCTGCGGAAGCAGCGGACCTTCGACAACGGCAGCATCCCCCACCAGATCCACCTGGGAGAGCTGCACGCC

ATTCTGCGGCGGCAGGAAGATTTTTACCCATTCCTGAAGGACAACCGGGAAAAGATCGAGAAGATCCTGACCTTCCGCATC
CCCTACTACGTGGGCCCTCTGGCCAGGGGAAACAGCAGATTCGCCTGGATGACCAGAAAGAGCGAGGAAACCATCACCCCC
TGGAACTTCGAGGAAGTGGTGGACAAGGGCGCTTCCGCCCAGAGCTTCATCGAGCGGATGACCAACTTCGATAAGAACCTG
CCCAACGAGAAGGTGCTGCCCAAGCACAGCCTGCTGTACGAGTACTTCACCGTGTATAACGAGCTGACCAAAGTGAAATAC
GTGACCGAGGGAATGAGAAAGCCCGCCTTCCTGAGCGGCGAGCAGAAAAAGGCCATCGTGGACCTGCTGTTCAAGACCAAC
CGGAAAGTGACCGTGAAGCAGCTGAAAGAGGACTACTTCAAGAAAATCGAGTGCTTCGACTCCGTGGAAATCTCCGGCGTG
GAAGATCGGTTCAACGCCTCCCTGGGCACATACCACGATCTGCTGAAAATTATCAAGGACAAGGACTTCCTGGACAATGAG
GAAAACGAGGACATTCTGGAAGATATCGTGCTGACCCTGACACTGTTTGAGGACAGAGAGATGATCGAGGAACGGCTGAAA
ACCTATGCCCACCTGTTCGACGACAAAGTGATGAAGCAGCTGAAGCGGCGGAGATACACCGGCTGGGGCAGGCTGAGCCGG
AAGCTGATCAACGGCATCCGGGACAAGCAGTCCGGCAAGACAATCCTGGATTTCCTGAAGTCCGACGGCTTCGCCAACAGA
AACTTCATGCAGCTGATCCACGACGACAGCCTGACCTTTAAAGAGGACATCCAGAAAGCCCAGGTGTCCGGCCAGGGCGAT
AGCCTGCACGAGCACATTGCCAATCTGGCCGGCAGCCCCGCCATTAAGAAGGGCATCCTGCAGACAGTGAAGGTGGTGGAC
GAGCTCGTGAAAGTGATGGGCCGGCACAAGCCCGAGAACATCGTGATCGAAATGGCCAGAGAGAACCAGACCACCCAGAAG
GGACAGAAGAACAGCCGCGAGAGAATGAAGCGGATCGAAGAGGGCATCAAAGAGCTGGGCAGCCAGATCCTGAAAGAACAC
CCCGTGGAAAACACCCAGCTGCAGAACGAGAAGCTGTACCTGTACTACCTGCAGAATGGGCGGGATATGTACGTGGACCAG
GAACTGGACATCAACCGGCTGTCCGACTACGATGTGGACGCTATCGTGCCTCAGAGCTTTCTGAAGGACGACTCCATCGAC
AACAAGGTGCTGACCAGAAGCGACAAGAACCGGGGCAAGAGCGACAACGTGCCCTCCGAAGAGGTCGTGAAGAAGATGAAG
AACTACTGGCGGCAGCTGCTGAACGCCAAGCTGATTACCCAGAGAAAGTTCGACAATCTGACCAAGGCCGAGAGAGGCGGC
CTGAGCGAACTGGATAAGGCCGGCTTCATCAAGAGACAGCTGGTGGAAACCCGGCAGATCACAAAGCACGTGGCACAGATC
CTGGACTCCCGGATGAACACTAAGTACGACGAGAATGACAAGCTGATCCGGGAAGTGAAAGTGATCACCCTGAAGTCCAAG
CTGGTGTCCGATTTCCGGAAGGATTTCCAGTTTTACAAAGTGCGCGAGATCAACAACTACCACCACGCCCACGACGCCTAC
CTGAACGCCGTCGTGGGAACCGCCCTGATCAAAAAGTACCCTAAGCTGGAAAGCGAGTTCGTGTACGGCGACTACAAGGTG
TACGACGTGCGGAAGATGATCGCCAAGAGCGAGCAGGAAATCGGCAAGGCTACCGCCAAGTACTTCTTCTACAGCAACATC
ATGAACTTTTTCAAGACCGAGATTACCCTGGCCAACGGCGAGATCCGGAAGCGGCCTCTGATCGAGACAAACGGCGAAACC
GGGGAGATCGTGTGGGATAAGGGCCGGGATTTTGCCACCGTGCGGAAAGTGCTGAGCATGCCCCAAGTGAATATCGTGAAA
AAGACCGAGGTGCAGACAGGCGGCTTCAGCAAAGAGTCTATCCTGCCCAAGAGGAACAGCGATAAGCTGATCGCCAGAAAG
AAGGACTGGGACCCTAAGAAGTACGGCGGCTTCGACAGCCCCACCGTGGCCTATTCTGTGCTGGTGGTGGCCAAAGTGGAA
AAGGGCAAGTCCAAGAAACTGAAGAGTGTGAAAGAGCTGCTGGGGATCACCATCATGGAAAGAAGCAGCTTCGAGAAGAAT
CCCATCGACTTTCTGGAAGCCAAGGGCTACAAAGAAGTGAAAAAGGACCTGATCATCAAGCTGCCTAAGTACTCCCTGTTC
GAGCTGGAAAACGGCCGGAAGAGAATGCTGGCCTCTGCCGGCGAACTGCAGAAGGGAAACGAACTGGCCCTGCCCTCCAAA
TATGTGAACTTCCTGTACCTGGCCAGCCACTATGAGAAGCTGAAGGGCTCCCCCGAGGATAATGAGCAGAAACAGCTGTTT
GTGGAACAGCACAAGCACTACCTGGACGAGATCATCGAGCAGATCAGCGAGTTCTCCAAGAGAGTGATCCTGGCCGACGCT
AATCTGGACAAAGTGCTGTCCGCCTACAACAAGCACCGGGATAAGCCCATCAGAGAGCAGGCCGAGAATATCATCCACCTG
TTTACCCTGACCAATCTGGGAGCCCCTGCCGCCTTCAAGTACTTTGACACCACCATCGACCGGAAGAGGTACACCAGCACC
AAAGAGGTGCTGGACGCCACCCTGATCCACCAGAGCATCACCGGCCTGTACGAGACACGGATCGACCTGTCTCAGCTGGGA
GGCGACAGCGCTGGAGGAGGTGGAAGCGGAGGAGGAGGAAGCGGAGGAGGAGGTAGCGGACCTAAGAAAAAGAGGAAGGTG
GCGGCCGCTGGATCCATGACAAATCGATCTGGTTCTCAAGATAGTACCAGCGATCTGATTCCGGCGCCGCCGCTGAGCAAG
GTGCCGCTGCAGCAAAACTTCCAAGACAACCAGTTTCAGGGTAAATGGTACATCGTTGGTTACGCGGGCAACGACTTCCTG
CGTGAGGACAAGGACCCGTTCAAAATGTGGGCGCGTATTTATGAGCTGAAGGAAGACAAAAGCTACAACGTGACCGGTGTT
TTCTTCCGTAAGAAGAAATGCTACTATCATATTCATACCTTCGTGCCGGGTAGCCAACCGGGCGAATTTACCCTGGGTAAC
ATTAAGAGCTACCCGGGCTACACCAGCCAGCTGGTTCGTGTGGTTAGCACCAACTATAACCAGCACGCGATGGTGTTCACT
AAGTACGTTTACCAAAACCGTGAGATGTTCTACATTGTTCTGTATGGCCGTACCAAGGAACTGACCAGCGAGCTGAAAGAA
AACTTCATCCGTTTTAGCAAAAGCCTGGGTCTGCCGGAAAACCACATTGTGTTTCCGGTTCCGATCGACCAATGCATTGAT
GGCTAA

>dCas9 AzoTag16 NLS(nuclear localization sequence)

**pCMV-HaloTag-VPR**

>Amino acid sequence

MAEIGTGFPFDPHYVEVLGERMHYVDVGPRDGTPVLFLHGNPTSSYVWRNIIPHVAPTHRCIAPDLIGMGKSDKPDLGYFF
DDHVRFMDAFIEALGLEEVVLVIHDWGSALGFHWAKRNPERVKGIAFMEFIRPIPTWDEWPEFARETFQAFRTTDVGRKLI
IDQNVFIEGTLPMGVVRPLTEVEMDHYREPFLNPVDREPLWRFPNELPIAGEPANIVALVEEYMDWLHQSPVPKLLFWGTP
GVLIPPAEAARLAKSLPNCKAVDIGPGLNLLQEDNPDLIGSEIARWLSTLEISGPGGGGGSGGGGSGGGGSGPKKKRKVAA
AGSEASGSGRADALDDFDLDMLGSDALDDFDLDMLGSDALDDFDLDMLGSDALDDFDLDMLINSRSSGSPKKKRKVGSQYL
PDTDDRHRIEEKRKRTYETFKSIMKKSPFSGPTDPRPPPRRIAVPSRSSASVPKPAPQPYPFTSSLSTINYDEFPTMVFPS
GQISQASALAPAPPQVLPQAPAPAPAPAMVSALAQAPAPVPVLAPGPPQAVAPPAPKPTQAGEGTLSEALLQLQFDDEDLG
ALLGNSTDPAVFTDLASVDNSEFQQLLNQGIPVAPHTTEPMLMEYPEAITRLVTGAQRPPDPAPAPLGAPGLPNGLLSGDE
DFSSIADMDFSALLGSGSGSRDSREGMFLPKPEAGSAISDVFEGREVCQPKRIRPFHPPGSPWANRPLPASLAPTPTGPVH
EPVGSLTPAPVPQPLDPAPAVTPEASHLLEDPDEETSQAVKALREMADTVIPQKEEAAICGQMDLSHPPPRGHLDELTTTL
ESMTEDLNLDSPLTPELNEILDTFLNDECLLHAMHISTGLSIFDTSLF*

>DNA sequence

ATGGCAGAAATCGGTACTGGCTTTCCATTCGACCCCCATTATGTGGAAGTCCTGGGCGAGCGCATGCACTACGTCGATGTT
GGTCCGCGCGATGGCACCCCTGTGCTGTTCCTGCACGGTAACCCGACCTCCTCCTACGTGTGGCGCAACATCATCCCGCAT
GTTGCACCGACCCATCGCTGCATTGCTCCAGACCTGATCGGTATGGGCAAATCCGACAAACCAGACCTGGGTTATTTCTTC
GACGACCACGTCCGCTTCATGGATGCCTTCATCGAAGCCCTGGGTCTGGAAGAGGTCGTCCTGGTCATTCACGACTGGGGC
TCCGCTCTGGGTTTCCACTGGGCCAAGCGCAATCCAGAGCGCGTCAAAGGTATTGCATTTATGGAGTTCATCCGCCCTATC
CCGACCTGGGACGAATGGCCAGAATTTGCCCGCGAGACCTTCCAGGCCTTCCGCACCACCGACGTCGGCCGCAAGCTGATC
ATCGATCAGAACGTTTTTATCGAGGGTACGCTGCCGATGGGTGTCGTCCGCCCGCTGACTGAAGTCGAGATGGACCATTAC
CGCGAGCCGTTCCTGAATCCTGTTGACCGCGAGCCACTGTGGCGCTTCCCAAACGAGCTGCCAATCGCCGGTGAGCCAGCG

```
AACATCGTCGCGCTGGTCGAAGAATACATGGACTGGCTGCACCAGTCCCCTGTCCCGAAGCTGCTGTTCTGGGGCACCCCA
GGCGTTCTGATCCCACCGGCCGAAGCCGCTCGCCTGGCCAAAAGCCTGCCTAACTGCAAGGCTGTGGACATCGGCCCGGGT
CTGAATCTGCTGCAAGAAGACAACCCGGACCTGATCGGCAGCGAGATCGCGCGCTGGCTGTCCACGCTGGAGATTTCCGGC
CCCGGGGGAGGAGGTGGTAGCGGAGGAGGAGGAAGCGGTGGTGGCGGTAGCGGACCTAAGAAAAAGAGGAAGGTGGCGGCC
GCTGGATCCGAGGCCAGCGGTTCCGGACGGGCTGACGCATTGGACGATTTTGATCTGGATATGCTGGGAAGTGACGCCCTC
GATGATTTTGACCTTGACATGCTTGGTTCGGATGCCCTTGATGACTTTGACCTCGACATGCTCGGCAGTGACGCCCTTGAT
GATTTCGACCTGGACATGCTGATTAACTCTAGAAGTTCCGGATCTCCGAAAAAGAAACGCAAAGTTGGTAGCCAGTACCTG
CCCGACACCGACGACCGGCACCGGATCGAGGAAAAGCGGAAGCGGACCTACGAGACATTCAAGAGCATCATGAAGAAGTCC
CCCTTCAGCGGCCCCACCGACCCTAGACCTCCACCTAGAAGAATCGCCGTGCCCAGCAGATCCAGCGCCAGCGTGCCAAAA
CCTGCCCCCCAGCCTTACCCCTTCACCAGCAGCCTGAGCACCATCAACTACGACGAGTTCCCTACCATGGTGTTCCCCAGC
GGCCAGATCTCTCAGGCCTCTGCTCTGGCTCCAGCCCCTCCTCAGGTGCTGCCTCAGGCTCCTGCTCCTGCACCAGCTCCA
GCCATGGTGTCTGCACTGGCTCAGGCACCAGCACCCGTGCCTGTGCTGGCTCCTGGACCTCCACAGGCTGTGGCTCCACCA
GCCCCTAAACCTACACAGGCCGGCGAGGGCACACTGTCTGAAGCTCTGCTGCAGCTGCAGTTCGACGACGAGGATCTGGGA
GCCCTGCTGGGAAACAGCACCGATCCTGCCGTGTTCACCGACCTGGCCAGCGTGGACAACAGCGAGTTCCAGCAGCTGCTG
AACCAGGGCATCCCTGTGGCCCCTCACACCACCGAGCCCATGCTGATGGAATACCCCGAGGCCATCACCCGGCTCGTGACA
GGCGCTCAGAGGCCTCCTGATCCAGCTCCTGCCCCTCTGGGAGCACCAGGCCTGCCTAATGGACTGCTGTCTGGCGACGAG
GACTTCAGCTCTATCGCCGATATGGATTTCTCAGCCTTGCTGGGCTCTGGCAGCGGCAGCCGGGATTCCAGGGAAGGGATG
TTTTTGCCGAAGCCTGAGGCCGGCTCCGCTATTAGTGACGTGTTTGAGGGCCGCGAGGTGTGCCAGCCAAAACGAATCCGG
CCATTTCATCCTCCAGGAAGTCCATGGGCCAACCGCCCACTCCCCGCCAGCCTCGCACCAACACCAACCGGTCCAGTACAT
GAGCCAGTCGGGTCACTGACCCCGGCACCAGTCCCTCAGCCACTGGATCCAGCGCCCGCAGTGACTCCCGAGGCCAGTCAC
CTGTTGGAGGATCCCGATGAAGAGACGAGCCAGGCTGTCAAAGCCCTTCGGGAGATGGCCGATACTGTGATTCCCCAGAAG
GAAGAGGCTGCAATCTGTGGCCAAATGGACCTTTCCCATCCGCCCCCAAGGGGCCATCTGGATGAGCTGACAACCACACTT
GAGTCCATGACCGAGGATCTGAACCTGGACTCACCCCTGACCCCGGAATTGAACGAGATTCTGGATACCTTCCTGAACGAC
GAGTGCCTCTTGCATGCCATGCATATCAGCACAGGACTGTCCATCTTCGACACATCTCTGTTTTAA
>HaloTag VPR NLS
```

**pCMV-AzoTag16-miRFP670**

```
>Amino acid sequence
MAQDSTSDLIPAPPLSKVPLQQNFQDNQFQGKWYIVGYAGNDFLREDKDPFKMWARIYELKEDKSYNVTGVFFRKKKCYYH
IHTFVPGSQPGEFTLGNIKSYPGYTSQLVRVVSTNYNQHAMVFTKYVYQNREMFYIVLYGRTKELTSELKENFIRFSKSLG
LPENHIVFPVPIDQCIDGSGGSPPVATMVAGHASGSPAFGTASHSNCEHEEIHLAGSIQPHGALLVVSEHDHRVIQASANA
AEFLNLGSVLGVPLAEIDGDLLIKILPHLDPTAEGMPVAVRCRIGNPSTEYCGLMHRPPEGGLIIELERAGPSIDLSGTLA
PALERIRTAGSLRALCDDTVLLFQQCTGYDRVMVYRFDEQGHGLVFSECHVPGLESYFGNRYPSSTVPQMARQLYVRQRVR
VLVDVTYQPVPLEPRLSPLTGRDLDMSGCFLRSMSPCHLQFLKDMGVRATLAVSLVVGGKLWGLVVCHHYLPRFIRFELRA
ICKRLAERIATRITALES*
>DNA sequence
ATGGCCCAAGATAGTACCAGCGATCTGATTCCGGCGCCGCCGCTGAGCAAGGTGCCGCTGCAGCAAAACTTCCAAGACAAC
CAGTTTCAGGGTAAATGGTACATCGTTGGTTACGCGGGCAACGACTTCCTGCGTGAGGACAAGGACCCGTTCAAAATGTGG
GCGCGTATTTATGAGCTGAAGGAAGACAAAAGCTACAACGTGACCGGTGTTTTCTTCCGTAAGAAGAAATGCTACTATCAT
ATTCATACCTTCGTGCCGGGTAGCCAACCGGGCGAATTTACCCTGGGTAACATTAAGAGCTACCCGGGCTACACCAGCCAG
CTGGTTCGTGTGGTTAGCACCAACTATAACCAGCACGCGATGGTGTTCACTAAGTACGTTTACCAAAACCGTGAGATGTTC
TACATTGTTCTGTATGGCCGTACCAAGGAACTGACCAGCGAGCTGAAAGAAAACTTCATCCGTTTTAGCAAAAGCCTGGGT
CTGCCGGAAAACCACATTGTGTTTCCGGTTCCGATCGACCAATGCATTGATGGCTCTGGTGGTAGTCCACCGGTCGCCACC
ATGGTAGCAGGTCATGCCTCTGGCAGCCCCGCATTCGGGACCGCCTCTCATTCGAATTGCGAACATGAAGAGATCCACCTC
GCCGGCTCGATCCAGCCGCATGGCGCGCTTCTGGTCGTCAGCGAACATGATCATCGCGTCATCCAGGCCAGCGCCAACGCC
GCGGAATTTCTGAATCTCGGAAGCGTACTCGGCGTTCCGCTCGCCGAGATCGACGGCGATCTGTTGATCAAGATCCTGCCG
CATCTCGATCCCACCGCCGAAGGCATGCCGGTCGCGGTGCGCTGCCGGATCGGCAATCCCTCTACGGAGTACTGCGGTCTG
ATGCATCGGCCTCCGGAAGGCGGGCTGATCATCGAACTCGAACGTGCCGGCCCGTCGATCGATCTGTCAGGCACGCTGGCG
CCGGCGCTGGAGCGGATCCGCACGGCGGGTTCACTGCGCGCGCTGTGCGATGACACCGTGCTGCTGTTTCAGCAGTGCACC
GGCTACGACCGGGTGATGGTGTATCGTTTCGATGAGCAAGGCCACGGCCTGGTATTCTCCGAGTGCCATGTGCCTGGGCTC
GAATCCTATTTCGGCAACCGCTATCCGTCGTCGACTGTCCCGCAGATGGCGCGGCAGCTGTACGTGCGGCAGCGCGTCCGC
GTGCTGGTCGACGTCACCTATCAGCCGGTGCCGCTGGAGCCGCGGCTGTCGCCGCTGACCGGGCGCGATCTCGACATGTCG
GGCTGCTTCCTGCGCTCGATGTCGCCGTGCCATCTGCAGTTCCTGAAGGACATGGGCGTGCGCGCCACCCTGGCGGTGTCG
CTGGTGGTCGGCGGCAAGCTGTGGGGCCTGGTTGTCTGTCACCATTATCTGCCGCGCTTCATCCGTTTCGAGCTGCGGGCG
ATCTGCAAACGGCTCGCCGAAAGGATCGCGACGCGGATCACCGCGCTTGAGAGCTAA
>AzoTag16 miRFP670
```

**pCMV-TOMM20-mCherry-HaloTag**

```
>Amino acid sequence
MVGRNSAIAAGVCGALFIGYCIYFDRKRRSDPNFKNRLRERRKKQKLAKERAGLSKLPDLKDAEAVQKFFLEEIQLGEELL
AQGEYEKGVDHLTNAIAVCGQPQQLLQVLQQTLPPPVFQMLLTKLPTISQRIVSAQSLAEDDVEAAASAPPVATMVSKGEE
DNMAIIKEFMRFKVHMEGSVNGHEFEIEGEGEGRPYEGTQTAKLKVTKGGPLPFAWDILSPQFMYGSKAYVKHPADIPDYL
KLSFPEGFKWERVMNFEDGGVVTVTQDSSLQDGEFIYKVKLRGTNFPSDGPVMQKKTMGWEASSERMYPEDGALKGEIKQR
LKLKDGGHYDAEVKTTYKAKKPVQLPGAYNVNIKLDITSHNEDYTIVEQYERAEGRHSTGGMDELYKGSAGSMAEIGTGFP
FDPHYVEVLGERMHYVDVGPRDGTPVLFLHGNPTSSYVWRNIIPHVAPTHRCIAPDLIGMGKSDKPDLGYFFDDHVRFMDA
FIEALGLEEVVLVIHDWGSALGFHWAKRNPERVKGIAFMEFIRPIPTWDEWPEFARETFQAFRTTDVGRKLIIDQNVFIEG
TLPMGVVRPLTEVEMDHYREPFLNPVDREPLWRFPNELPIAGEPANIVALVEEYMDWLHQSPVPKLLFWGTPGVLIPPAEA
```

ARLAKSLPNCKAVDIGPGLNLLQEDNPDLIGSEIARWLSTLEISG*
>DNA sequence
ATGGTGGGTCGGAACAGCGCCATCGCCGCCGGTGTATGCGGAGCCCTTTTCATTGGGTACTGCATCTACTTCGACCGCAAA
AGACGAAGTGACCCCAACTTCAAGAACAGGCTTCGAGAACGAAGAAAGAAACAGAAGCTGGCCAAGGAGAGAGCTGGGCTT
TCCAAGTTACCTGACCTTAAAGATGCTGAAGCTGTTCAGAAGTTCTTCCTTGAAGAAATACAGCTTGGTGAAGAGTTACTA
GCTCAAGGTGAATATGAGAAGGGCGTAGACCATCTGACAAATGCAATAGCTGTGTGTGGACAGCCACAGCAGTTACTGCAA
GTCTTACAGCAAACTCTTCCACCACCAGTGTTCCAGATGCTTCTGACTAAGCTCCCAACAATTAGTCAGAGAATTGTAAGT
GCTCAGAGCTTGGCTGAAGATGATGTGGAAGCGGCCGCTTCGGCTCCACCGGTCGCCACCATGGTGAGCAAGGGCGAGGAG
GATAACATGGCCATCATCAAGGAGTTCATGCGCTTCAAGGTGCACATGGAGGGCTCCGTGAACGGCCACGAGTTCGAGATC
GAGGGCGAGGGCGAGGGCCGCCCCTACGAGGGCACCCAGACCGCCAAGCTGAAGGTGACCAAGGGTGGCCCCCTGCCCTTC
GCCTGGGACATCCTGTCCCCTCAGTTCATGTACGGCTCCAAGGCCTACGTGAAGCACCCCGCCGACATCCCCGACTACTTG
AAGCTGTCCTTCCCCGAGGGCTTCAAGTGGGAGCGCGTGATGAACTTCGAGGACGGCGGCGTGGTGACCGTGACCCAGGAC
TCCTCCCTGCAGGACGGCGAGTTCATCTACAAGGTGAAGCTGCGCGGCACCAACTTCCCCTCCGACGGCCCCGTAATGCAG
AAGAAGACCATGGGCTGGGAGGCCTCCTCCGAGCGGATGTACCCCGAGGACGGCGCCCTGAAGGGCGAGATCAAGCAGAGG
CTGAAGCTGAAGGACGGCGGCCACTACGACGCTGAGGTCAAGACCACCTACAAGGCCAAGAAGCCCGTGCAGCTGCCCGGC
GCCTACAACGTCAACATCAAGTTGGACATCACCTCCCACAACGAGGACTACACCATCGTGGAACAGTACGAACGCGCCGAG
GGCCGCCACTCCACCGGCGGCATGGACGAGCTGTACAAGGGTAGCGCTGGATCCATGGCAGAAATCGGTACTGGCTTTCCA
TTCGACCCCCATTATGTGGAAGTCCTGGGCGAGCGCATGCACTACGTCGATGTTGGTCCGCGCGATGGCACCCCTGTGCTG
TTCCTGCACGGTAACCCGACCTCCTCCTACGTGTGGCGCAACATCATCCCGCATGTTGCACCGACCCATCGCTGCATTGCT
CCAGACCTGATCGGTATGGGCAAATCCGACAAACCAGACCTGGGTTATTTCTTCGACGACCACGTCCGCTTCATGGATGCC
TTCATCGAAGCCCTGGGTCTGGAAGAGGTCGTCCTGGTCATTCACGACTGGGGCTCCGCTCTGGGTTTCCACTGGGCCAAG
CGCAATCCAGAGCGCGTCAAAGGTATTGCATTTATGGAGTTCATCCGCCCTATCCCGACCTGGGACGAATGGCCAGAATTT
GCCCGCGAGACCTTCCAGGCCTTCCGCACCACCGACGTCGGCCGCAAGCTGATCATCGATCAGAACGTTTTTATCGAGGGT
ACGCTGCCGATGGGTGTCGTCCGCCCGCTGACTGAAGTCGAGATGGACCATTACCGCGAGCCGTTCCTGAATCCTGTTGAC
CGCGAGCCACTGTGGCGCTTCCCAAACGAGCTGCCAATCGCCGGTGAGCCAGCGAACATCGTCGCGCTGGTCGAAGAATAC
ATGGACTGGCTGCACCAGTCCCCTGTCCCGAAGCTGCTGTTCTGGGGCACCCCAGGCGTTCTGATCCCACCGGCCGAAGCC
GCTCGCCTGGCCAAAAGCCTGCCTAACTGCAAGGCTGTGGACATCGGCCCAGGTCTGAATCTGCTGCAAGAAGACAACCCG
GACCTGATCGGCAGCGAGATCGCGCGCTGGCTGTCCACGCTGGAGATTTCCGGCTAG
>TOMM20 mCherry HaloTag

**pCMV-mCherry-HaloTag-NLS(x3)**
>Amino acid sequence
MVSKGEEDNMAIIKEFMRFKVHMEGSVNGHEFEIEGEGEGRPYEGTQTAKLKVTKGGPLPFAWDILSPQFMYGSKAYVKHP
ADIPDYLKLSFPEGFKWERVMNFEDGGVVTVTQDSSLQDGEFIYKVKLRGTNFPSDGPVMQKKTMGWEASSERMYPEDGAL
KGEIKQRLKLKDGGHYDAEVKTTYKAKKPVQLPGAYNVNIKLDITSHNEDYTIVEQYERAEGRHSTGGMDELYKSGLRSRA
QASNSAMAEIGTGFPFDPHYVEVLGERMHYVDVGPRDGTPVLFLHGNPTSSYVWRNIIPHVAPTHRCIAPDLIGMGKSDKP
DLGYFFDDHVRFMDAFIEALGLEEVVLVIHDWGSALGFHWAKRNPERVKGIAFMEFIRPIPTWDEWPEFARETFQAFRTTD
VGRKLIIDQNVFIEGTLPMGVVRPLTEVEMDHYREPFLNPVDREPLWRFPNELPIAGEPANIVALVEEYMDWLHQSPVPKL
LFWGTPGVLIPPAEAARLAKSLPNCKAVDIGPGLNLLQEDNPDLIGSEIARWLSTLEISDGTGSAGEPKKKRKVEPKKKRK
VEPKKKRKVGSTGSR*
>DNA sequence
ATGGTGAGCAAGGGCGAGGAGGATAACATGGCCATCATCAAGGAGTTCATGCGCTTCAAGGTGCACATGGAGGGCTCCGTG
AACGGCCACGAGTTCGAGATCGAGGGCGAGGGCGAGGGCCGCCCCTACGAGGGCACCCAGACCGCCAAGCTGAAGGTGACC
AAGGGTGGCCCCCTGCCCTTCGCCTGGGACATCCTGTCCCCTCAGTTCATGTACGGCTCCAAGGCCTACGTGAAGCACCCC
GCCGACATCCCCGACTACTTGAAGCTGTCCTTCCCCGAGGGCTTCAAGTGGGAGCGCGTGATGAACTTCGAGGACGGCGGC
GTGGTGACCGTGACCCAGGACTCCTCCCTGCAGGACGGCGAGTTCATCTACAAGGTGAAGCTGCGCGGCACCAACTTCCCC
TCCGACGGCCCCGTAATGCAGAAGAAGACCATGGGCTGGGAGGCCTCCTCCGAGCGGATGTACCCCGAGGACGGCGCCCTG
AAGGGCGAGATCAAGCAGAGGCTGAAGCTGAAGGACGGCGGCCACTACGACGCTGAGGTCAAGACCACCTACAAGGCCAAG
AAGCCCGTGCAGCTGCCCGGCGCCTACAACGTCAACATCAAGTTGGACATCACCTCCCACAACGAGGACTACACCATCGTG
GAACAGTACGAACGCGCCGAGGGCCGCCACTCCACCGGCGGCATGGACGAGCTGTACAAGTCCGGACTCAGATCTCGAGCT
CAAGCTTCGAATTCTGCAATGGCAGAAATCGGTACTGGCTTTCCATTCGACCCCCATTATGTGGAAGTCCTGGGCGAGCGC
ATGCACTACGTCGATGTTGGTCCGCGCGATGGCACCCCTGTGCTGTTCCTGCACGGTAACCCGACCTCCTCCTACGTGTGG
CGCAACATCATCCCGCATGTTGCACCGACCCATCGCTGCATTGCTCCAGACCTGATCGGTATGGGCAAATCCGACAAACCA
GACCTGGGTTATTTCTTCGACGACCACGTCCGCTTCATGGATGCCTTCATCGAAGCCCTGGGTCTGGAAGAGGTCGTCCTG
GTCATTCACGACTGGGGCTCCGCTCTGGGTTTCCACTGGGCCAAGCGCAATCCAGAGCGCGTCAAAGGTATTGCATTTATG
GAGTTCATCCGCCCTATCCCGACCTGGGACGAATGGCCAGAATTTGCCCGCGAGACCTTCCAGGCCTTCCGCACCACCGAC
GTCGGCCGCAAGCTGATCATCGATCAGAACGTTTTTATCGAGGGTACGCTGCCGATGGGTGTCGTCCGCCCGCTGACTGAA
GTCGAGATGGACCATTACCGCGAGCCGTTCCTGAATCCTGTTGACCGCGAGCCACTGTGGCGCTTCCCAAACGAGCTGCCA
ATCGCCGGTGAGCCAGCGAACATCGTCGCGCTGGTCGAAGAATACATGGACTGGCTGCACCAGTCCCCTGTCCCGAAGCTG
CTGTTCTGGGGCACCCCAGGCGTTCTGATCCCACCGGCCGAAGCCGCTCGCCTGGCCAAAAGCCTGCCTAACTGCAAGGCT
GTGGACATCGGCCCGGGTCTGAATCTGCTGCAAGAAGACAACCCGGACCTGATCGGCAGCGAGATCGCGCGCTGGCTGTCG
ACGCTCGAGATTTCCGACGGTACCGGATCTGCTGGAGAACCCAAGAAGAAGCGCAAGGTTGAGCCAAAGAAGAAAAGAAAG
GTGGAGCCCAAAAAGAAGCGCAAAGTGGGATCCACCGGATCTAGATAA
>mCherry HaloTag NLS

**pCMV-HaloTag-mCherry-Cb5**
>Amino acid sequence

MAEIGTGFPFDPHYVEVLGERMHYVDVGPRDGTPVLFLHGNPTSSYVWRNIIPHVAPTHRCIAPDLIGMGKSDKPDLGYFF
DDHVRFMDAFIEALGLEEVVLVIHDWGSALGFHWAKRNPERVKGIAFMEFIRPIPTWDEWPEFARETFQAFRTTDVGRKLI
IDQNVFIEGTLPMGVVRPLTEVEMDHYREPFLNPVDREPLWRFPNELPIAGEPANIVALVEEYMDWLHQSPVPKLLFWGTP
GVLIPPAEAARLAKSLPNCKAVDIGPGLNLLQEDNPDLIGSEIARWLSTLEISGAAASAPPVATMVSKGEEDNMAIIKEFM
RFKVHMEGSVNGHEFEIEGEGEGRPYEGTQTAKLKVTKGGPLPFAWDILSPQFMYGSKAYVKHPADIPDYLKLSFPEGFKW
ERVMNFEDGGVVTVTQDSSLQDGEFIYKVKLRGTNFPSDGPVMQKKTMGWEASSERMYPEDGALKGEIKQRLKLKDGGHYD
AEVKTTYKAKKPVQLPGAYNVNIKLDITSHNEDYTIVEQYERAEGRHSTGGMDELYKGTGGGGGSGGGGARAITTVESNSS
WWTNWVIPAISALVVALMYRLYMAED*
>DNA sequence
ATGGCAGAAATCGGTACTGGCTTTCCATTCGACCCCCATTATGTGGAAGTCCTGGGCGAGCGCATGCACTACGTCGATGTT
GGTCCGCGCGATGGCACCCCTGTGCTGTTCCTGCACGGTAACCCGACCTCCTCCTACGTGTGGCGCAACATCATCCCGCAT
GTTGCACCGACCCATCGCTGCATTGCTCCAGACCTGATCGGTATGGGCAAATCCGACAAACCAGACCTGGGTTATTTCTTC
GACGACCACGTCCGCTTCATGGATGCCTTCATCGAAGCCCTGGGTCTGGAAGAGGTCGTCCTGGTCATTCACGACTGGGGC
TCCGCTCTGGGTTTCCACTGGGCCAAGCGCAATCCAGAGCGCGTCAAAGGTATTGCATTTATGGAGTTCATCCGCCCTATC
CCGACCTGGGACGAATGGCCAGAATTTGCCCGCGAGACCTTCCAGGCCTTCCGCACCACCGACGTCGGCCGCAAGCTGATC
ATCGATCAGAACGTTTTTATCGAGGGTACGCTGCCGATGGGTGTCGTCCGCCCGCTGACTGAAGTCGAGATGGACCATTAC
CGCGAGCCGTTCCTGAATCCTGTTGACCGCGAGCCACTGTGGCGCTTCCCAAACGAGCTGCCAATCGCCGGTGAGCCAGCG
AACATCGTCGCGCTGGTCGAAGAATACATGGACTGGCTGCACCAGTCCCCTGTCCCGAAGCTGCTGTTCTGGGGCACCCCA
GGCGTTCTGATCCCACCGGCCGAAGCCGCTCGCCTGGCCAAAAGCCTGCCTAACTGCAAGGCTGTGGACATCGGCCCAGGT
CTGAATCTGCTGCAAGAAGACAACCCGGACCTGATCGGCAGCGAGATCGCGCGCTGGCTGTCCACGCTGGAGATTTCCGGC
GCGGCCGCTTCGGCTCCACCGGTCGCCACCATGGTGAGCAAGGGCGAGGAGGATAACATGGCCATCATCAAGGAGTTCATG
CGCTTCAAGGTGCACATGGAGGGCTCCGTGAACGGCCACGAGTTCGAGATCGAGGGCGAGGGCGAGGGCCGCCCCTACGAG
GGCACCCAGACCGCCAAGCTGAAGGTGACCAAGGGTGGCCCCCTGCCCTTCGCCTGGGACATCCTGTCCCCTCAGTTCATG
TACGGCTCCAAGGCCTACGTGAAGCACCCCGCCGACATCCCCGACTACTTGAAGCTGTCCTTCCCCGAGGGCTTCAAGTGG
GAGCGCGTGATGAACTTCGAGGACGGCGGCGTGGTGACCGTGACCCAGGACTCCTCCCTGCAGGACGGCGAGTTCATCTAC
AAGGTGAAGCTGCGCGGCACCAACTTCCCCTCCGACGGCCCCGTAATGCAGAAGAAGACCATGGGCTGGGAGGCCTCCTCC
GAGCGGATGTACCCCGAGGACGGCGCCCTGAAGGGCGAGATCAAGCAGAGGCTGAAGCTGAAGGACGGCGGCCACTACGAC
GCTGAGGTCAAGACCACCTACAAGGCCAAGAAGCCCGTGCAGCTGCCCGGCGCCTACAACGTCAACATCAAGTTGGACATC
ACCTCCCACAACGAGGACTACACCATCGTGGAACAGTACGAACGCGCCGAGGGCCGCCACTCCACCGGCGGCATGGACGAG
CTGTACAAGGGTACCGGAGGCGGAGGTGGGTCGGGTGGCGGCGGAGCTCGAGCTATCACCACCGTGGAGTCCAACTCCTCC
TGGTGGACCAACTGGGTGATCCCCGCCATCTCCGCCCTGGTGGTGGCCCTGATGTACCGCCTGTACATGGCCGAGGACTAG
>HaloTag mCherry Cb5(amino acids 100-134 of cytochrome *b*$_5$)

**pCMV-HaloTag-mCherry-Giantin**

>Amino acid sequence
MAEIGTGFPFDPHYVEVLGERMHYVDVGPRDGTPVLFLHGNPTSSYVWRNIIPHVAPTHRCIAPDLIGMGKSDKPDLGYFF
DDHVRFMDAFIEALGLEEVVLVIHDWGSALGFHWAKRNPERVKGIAFMEFIRPIPTWDEWPEFARETFQAFRTTDVGRKLI
IDQNVFIEGTLPMGVVRPLTEVEMDHYREPFLNPVDREPLWRFPNELPIAGEPANIVALVEEYMDWLHQSPVPKLLFWGTP
GVLIPPAEAARLAKSLPNCKAVDIGPGLNLLQEDNPDLIGSEIARWLSTLEISGAAASAPPVATMVSKGEEDNMAIIKEFM
RFKVHMEGSVNGHEFEIEGEGEGRPYEGTQTAKLKVTKGGPLPFAWDILSPQFMYGSKAYVKHPADIPDYLKLSFPEGFKW
ERVMNFEDGGVVTVTQDSSLQDGEFIYKVKLRGTNFPSDGPVMQKKTMGWEASSERMYPEDGALKGEIKQRLKLKDGGHYD
AEVKTTYKAKKPVQLPGAYNVNIKLDITSHNEDYTIVEQYERAEGRHSTGGMDELYKRSRGEPQQSFSEAQQQLCNTRQEV
NELRKLLEEERDQRVAAENALSVAEEQIRRLEHSEWDSSRTPIIGSCGTQEQALLIDLTSNSCRRTRSGVGWKRVLRSLCH
SRTRVPLLAAIYFLMIHVLLILCFTGHL*
>DNA sequence
ATGGCAGAAATCGGTACTGGCTTTCCATTCGACCCCCATTATGTGGAAGTCCTGGGCGAGCGCATGCACTACGTCGATGTT
GGTCCGCGCGATGGCACCCCTGTGCTGTTCCTGCACGGTAACCCGACCTCCTCCTACGTGTGGCGCAACATCATCCCGCAT
GTTGCACCGACCCATCGCTGCATTGCTCCAGACCTGATCGGTATGGGCAAATCCGACAAACCAGACCTGGGTTATTTCTTC
GACGACCACGTCCGCTTCATGGATGCCTTCATCGAAGCCCTGGGTCTGGAAGAGGTCGTCCTGGTCATTCACGACTGGGGC
TCCGCTCTGGGTTTCCACTGGGCCAAGCGCAATCCAGAGCGCGTCAAAGGTATTGCATTTATGGAGTTCATCCGCCCTATC
CCGACCTGGGACGAATGGCCAGAATTTGCCCGCGAGACCTTCCAGGCCTTCCGCACCACCGACGTCGGCCGCAAGCTGATC
ATCGATCAGAACGTTTTTATCGAGGGTACGCTGCCGATGGGTGTCGTCCGCCCGCTGACTGAAGTCGAGATGGACCATTAC
CGCGAGCCGTTCCTGAATCCTGTTGACCGCGAGCCACTGTGGCGCTTCCCAAACGAGCTGCCAATCGCCGGTGAGCCAGCG
AACATCGTCGCGCTGGTCGAAGAATACATGGACTGGCTGCACCAGTCCCCTGTCCCGAAGCTGCTGTTCTGGGGCACCCCA
GGCGTTCTGATCCCACCGGCCGAAGCCGCTCGCCTGGCCAAAAGCCTGCCTAACTGCAAGGCTGTGGACATCGGCCCAGGT
CTGAATCTGCTGCAAGAAGACAACCCGGACCTGATCGGCAGCGAGATCGCGCGCTGGCTGTCCACGCTGGAGATTTCCGGC
GCGGCCGCTTCGGCTCCACCGGTCGCCACCATGGTGAGCAAGGGCGAGGAGGATAACATGGCCATCATCAAGGAGTTCATG
CGCTTCAAGGTGCACATGGAGGGCTCCGTGAACGGCCACGAGTTCGAGATCGAGGGCGAGGGCGAGGGCCGCCCCTACGAG
GGCACCCAGACCGCCAAGCTGAAGGTGACCAAGGGTGGCCCCCTGCCCTTCGCCTGGGACATCCTGTCCCCTCAGTTCATG
TACGGCTCCAAGGCCTACGTGAAGCACCCCGCCGACATCCCCGACTACTTGAAGCTGTCCTTCCCCGAGGGCTTCAAGTGG
GAGCGCGTGATGAACTTCGAGGACGGCGGCGTGGTGACCGTGACCCAGGACTCCTCCCTGCAGGACGGCGAGTTCATCTAC
AAGGTGAAGCTGCGCGGCACCAACTTCCCCTCCGACGGCCCCGTAATGCAGAAGAAGACCATGGGCTGGGAGGCCTCCTCC
GAGCGGATGTACCCCGAGGACGGCGCCCTGAAGGGCGAGATCAAGCAGAGGCTGAAGCTGAAGGACGGCGGCCACTACGAC
GCTGAGGTCAAGACCACCTACAAGGCCAAGAAGCCCGTGCAGCTGCCCGGCGCCTACAACGTCAACATCAAGTTGGACATC
ACCTCCCACAACGAGGACTACACCATCGTGGAACAGTACGAACGCGCCGAGGGCCGCCACTCCACCGGCGGCATGGACGAG
CTGTACAAGAGATCTCGAGGAGAACCGCAGCAAAGCTTTTCTGAAGCTCAGCAGCAGCTATGCAACACCAGACAGGAAGTG
AATGAATTAAGGAAGCTGCTGGAAGAAGAACGAGACCAAAGAGTGGCTGCTGAGAATGCTCTCTCTGTGGCCGAGGAGCAG
ATCAGACGGTTAGAGCACAGTGAATGGGACTCTTCCCGGACTCCTATCATTGGCTCCTGTGGCACTCAGGAGCAGGCACTG

TTAATAGATCTTACAAGCAACAGTTGTCGAAGGACCCGGAGTGGCGTTGGATGGAAGCGAGTCCTGCGTTCACTCTGTCAT
TCACGGACCCGAGTGCCACTTCTAGCAGCCATCTACTTTCTAATGATTCATGTCCTGCTCATTCTGTGTTTTACGGGCCAT
CTATAG
>HaloTag mCherry Giantin(amino acids 3131-3259 of Giantin)

**pCMV-EMTB-HaloTag-mCherry**

>Amino acid sequence
MEQKLISEEDLGAPVRSETAPDSYKVQDKKNASSRPASAISGQNNNHSGNKPDPPPVLRVDDRQRLARERREEREKQLAAR
EIVWLEREERARQHYEKHLEERKKRLEEQRQKEERRRAAVEEKRRQRLEEDKERHEAVVRRTMERSQKPKQKHNRWSWGGS
LHGSPSIHSADPDRRSVSTMNLSKYVDPVISKRLSSSSATLLNSPDRARRLQLSPWESSVVNRLLTPTHSFLARSKSTAAL
SGEAASCSPIIMPYKAAHSRNSMDRPKLFVTPPEGSSDPWMAEIGTGFPFDPHYVEVLGERMHYVDVGPRDGTPVLFLHGN
PTSSYVWRNIIPHVAPTHRCIAPDLIGMGKSDKPDLGYFFDDHVRFMDAFIEALGLEEVVLVIHDWGSALGFHWAKRNPER
VKGIAFMEFIRPIPTWDEWPEFARETFQAFRTTDVGRKLIIDQNVFIEGTLPMGVVRPLTEVEMDHYREPFLNPVDREPLW
RFPNELPIAGEPANIVALVEEYMDWLHQSPVPKLLFWGTPGVLIPPAEAARLAKSLPNCKAVDIGPGLNLLQEDNPDLIGS
EIARWLSTLEISGAAASAPPVATMVSKGEEDNMAIIKEFMRFKVHMEGSVNGHEFEIEGEGEGRPYEGTQTAKLKVTKGGP
LPFAWDILSPQFMYGSKAYVKHPADIPDYLKLSFPEGFKWERVMNFEDGGVVTVTQDSSLQDGEFIYKVKLRGTNFPSDGP
VMQKKTMGWEASSERMYPEDGALKGEIKQRLKLKDGGHYDAEVKTTYKAKKPVQLPGAYNVNIKLDITSHNEDYTIVEQYE
RAEGRHSTGGMDELYK*
>DNA sequence
ATGGAGCAGAAGCTCATCTCAGAAGAAGACCTCGGCGCGCCAGTGCGAAGCGAAACAGCACCCGACAGCTACAAAGTGCAA
GATAAGAAAAATGCCTCCAGCCGCCCTGCCTCTGCAATTTCAGGACAAAATAACAACCACTCAGGAAATAAACCAGACCCT
CCGCCTGTGTTACGTGTTGATGACCGGCAGCGGCTGGCCCGGGAGCGACGTGAGGAACGGGAGAAACAGCTAGCTGCAAGA
GAAATAGTGTGGTTAGAAAGAGAAGAGCGAGCCAGGCAGCACTACGAGAAGCACCTGGAAGAGCGGAAGAAGAGGTTGGAG
GAGCAGAGGCAGAAGGAGGAGCGGAGGAGGGCTGCTGTGGAGGAGAAGCGGAGGCAGAGACTTGAGGAGGACAAAGAACGC
CACGAAGCTGTTGTACGGCGCACAATGGAAAGGAGCCAGAAGCCAAAACAGAAGCATAACCGTTGGTCGTGGGGAGGCTCT
CTCCATGGGAGCCCTAGCATCCACAGTGCAGATCCAGACAGGCGGTCAGTTTCCACCATGAATCTTTCGAAATATGTTGAT
CCCGTCATTAGCAAGCGGCTCTCCTCTTCATCTGCAACTTTACTAAATTCTCCAGATAGAGCTCGCCGCCTGCAGCTCAGC
CCATGGGAGAGCAGCGTTGTTAACAGACTCCTGACGCCCACACATTCGTTCCTGGCCAGAAGTAAAAGCACAGCTGCCTTG
TCTGGAGAAGCAGCATCTTGCAGCCCCATCATCATGCCCTACAAAGCTGCACACTCTAGAAATTCGATGGATCGACCAAAA
CTCTTTGTAACACCACCTGAGGGCTCTTCGGATCCATGGATGGCAGAAATCGGTACTGGCTTTCCATTCGACCCCCATTAT
GTGGAAGTCCTGGGCGAGCGCATGCACTACGTCGATGTTGGTCCGCGCGATGGCACCCCTGTGCTGTTCCTGCACGGTAAC
CCGACCTCCTCCTACGTGTGGCGCAACATCATCCCGCATGTTGCACCGACCCATCGCTGCATTGCTCCAGACCTGATCGGT
ATGGGCAAATCCGACAAACCAGACCTGGGTTATTTCTTCGACGACCACGTCCGCTTCATGGATGCCTTCATCGAAGCCCTG
GGTCTGGAAGAGGTCGTCCTGGTCATTCACGACTGGGGCTCCGCTCTGGGTTTCCACTGGGCCAAGCGCAATCCAGAGCGC
GTCAAAGGTATTGCATTTATGGAGTTCATCCGCCCTATCCCGACCTGGGACGAATGGCCAGAATTTGCCCGCGAGACCTTC
CAGGCCTTCCGCACCACCGACGTCGGCCGCAAGCTGATCATCGATCAGAACGTTTTTATCGAGGGTACGCTGCCGATGGGT
GTCGTCCGCCCGCTGACTGAAGTCGAGATGGACCATTACCGCGAGCCGTTCCTGAATCCTGTTGACCGCGAGCCACTGTGG
CGCTTCCCAAACGAGCTGCCAATCGCCGGTGAGCCAGCGAACATCGTCGCGCTGGTCGAAGAATACATGGACTGGCTGCAC
CAGTCCCCTGTCCCGAAGCTGCTGTTCTGGGGCACCCCAGGCGTTCTGATCCCACCGGCCGAAGCCGCTCGCCTGGCCAAA
AGCCTGCCTAACTGCAAGGCTGTGGACATCGGCCCAGGTCTGAATCTGCTGCAAGAAGACAACCCGGACCTGATCGGCAGC
GAGATCGCGCGCTGGCTGTCCACGCTGGAGATTTCCGGCGCGGCCGCTTCGGCTCCACCGGTCGCCACCATGGTGAGCAAG
GGCGAGGAGGATAACATGGCCATCATCAAGGAGTTCATGCGCTTCAAGGTGCACATGGAGGGCTCCGTGAACGGCCACGAG
TTCGAGATCGAGGGCGAGGGCGAGGGCCGCCCCTACGAGGGCACCCAGACCGCCAAGCTGAAGGTGACCAAGGGTGGCCCC
CTGCCCTTCGCCTGGGACATCCTGTCCCCTCAGTTCATGTACGGCTCCAAGGCCTACGTGAAGCACCCCGCCGACATCCCC
GACTACTTGAAGCTGTCCTTCCCCGAGGGCTTCAAGTGGGAGCGCGTGATGAACTTCGAGGACGGCGGCGTGGTGACCGTG
ACCCAGGACTCCTCCCTGCAGGACGGCGAGTTCATCTACAAGGTGAAGCTGCGCGGCACCAACTTCCCCTCCGACGGCCCC
GTAATGCAGAAGAAGACCATGGGCTGGGAGGCCTCCTCCGAGCGGATGTACCCCGAGGACGGCGCCCTGAAGGGCGAGATC
AAGCAGAGGCTGAAGCTGAAGGACGGCGGCCACTACGACGCTGAGGTCAAGACCACCTACAAGGCCAAGAAGCCCGTGCAG
CTGCCCGGCGCCTACAACGTCAACATCAAGTTGGACATCACCTCCCACAACGAGGACTACACCATCGTGGAACAGTACGAA
CGCGCCGAGGGCCGCCACTCCACCGGCGGCATGGACGAGCTGTACAAGTAA
>EMTB[the microtubule-binding domain (amino acids 18-283) of ensconsin] HaloTag mCherry
